# The neural basis of creative thought: An activation likelihood estimation meta-analysis involving over 17,000 participants

**DOI:** 10.1101/2025.02.19.639155

**Authors:** Melody M.Y. Chan, Matthew A. Lambon Ralph, Gail A. Robinson

## Abstract

Humans can create impressive art, make major scientific breakthroughs, and generate creative solutions that solve everyday life problems. But how are these creative activities supported by the brain? This question is of great scientific interest but remains unanswered. New hypotheses, grounded in converging evidence from cognitive neuroscience, are needed to guide breakthroughs in understanding the neural basis of creative thought. It has long been suggested that creativity is a distinct mental function. However, converging clinical-cognitive neuroscience evidence is starting to suggest that creative thought is not functionally distinct but, instead, might arise from general purpose cognitive mechanisms supporting semantic cognition, controlled episodic memory retrieval, and executive mechanisms (i.e., *Cognitive Cornerstones Hypothesis*; Chan et al., 2023). Results from this large-scale activation likelihood estimation (ALE) meta-analysis, based on 787 experiments with 10,357 foci from 17,228 healthy adult participants, showed that the brain regions implicated in creativity tasks heavily overlap with the brain regions implicated in the cognitive cornerstones of creative thought (i.e., pre-existing knowledge and executive mechanisms). These striking results suggest that innovative insights will arise from considering the roles of these fundamental cognitive functions and their interactions in supporting creative thought.

## 1. Introduction

Creativity, the ability to generate novel ideas that support human advances and everyday adaptive problem solving, is a complex behaviour that has attracted great interest and research in recent decades. To date, most research on this topic has taken a domain-specific viewpoint in which the cognitive processes that might underpin creative thought have been considered in detail. Thus, the earliest conceptual frameworks (Guilford, 1967; Mednick, 1962) define creativity as a mental function that arises from two major processes, a generative process [commonly known as “divergent thinking”; (Guilford, 1967)] and an evaluation-selection process [(Campbell, 1960; Mednick, 1962), sometimes known as “convergent thinking”; Cropley (2006)]. Accumulative evidence suggests that creativity involves two aspects, 1) pre-existing knowledge that provides raw materials for idea generation (Beaty & Kenett, 2023; Benedek et al., 2023; Cropley, 2006; Dane & Pratt, 2007; Ericsson et al., 2007; Sternberg, 1988), and 2) an operational system that flexibly manipulates ideas in a goal directed manner (Abraham, 2019; Benedek & Fink, 2019; Chrysikou, 2019; Flaherty, 2005; Gruber & Wallace, 1999; Lebuda & Benedek, 2023; Mekern et al., 2019).

To study the cognitive neuroscience of creativity, different cognitive tasks that evaluate how people produce novel and task-relevant (i.e., appropriate) ideas have been used to tap participants’ creative behaviours. During these tasks, activity in specific fronto-parieto-temporal brain regions (Boccia et al., 2015; Gonen-Yaacovi et al., 2013; Kuang et al., 2022) and functional connectivity between these regions (Beaty et al., 2015; Beaty et al., 2017) have been associated with creative thought. Intriguingly, these same fronto-parieto-temporal brain regions are implicated in many cognitive domains and specific processes, and are not specific to creative thought *per se* [e.g., “multiple demand system”; Duncan (2010)]. Consequently, previous attempts to map creativity to specific brain regions or processes in a “one-to-one” fashion, may be insufficient, with some researchers concluding that “creative thinking does not appear to critically depend on any single mental process” (Dietrich & Kanso, 2010).

Recently, we proposed a new perspective for understanding the cognitive neuroscience behind creativity – the *Cognitive Cornerstones Hypothesis* of creative thought (Chan et al., 2023). This hypothesis defines the two fundamental “cornerstones” of creative thought, “pre-existing knowledge” and “executive mechanisms”, based on converging findings from creativity, clinical neuroscience, and cognitive neuroscience research. The central idea is that creative thought, like many other complex behaviours, arises from general purpose cognitive mechanisms (i.e., cornerstones) that can be dynamically and differentially combined to support different creative activities (Patterson & Lambon Ralph, 1999; Humphreys et al., 2021). We further proposed that, based on converging creativity research findings, the general purpose cognitive mechanisms constituting the cornerstones of creative thought include those found to be underlying semantic cognition (Lambon Ralph et al., 2017), controlled episodic memory retrieval (Burgess, 2006; Tulving, 2002) and executive mechanisms (Stuss, 2011; Stuss & Alexander, 2007). Defining these prerequisites offers a framework for refining how we study the cognition behind creativity (e.g., fine-grained tasks used for assessing creative thought to further dissect the cognitive and neural machineries specific to the generation/evaluation/selection of novel verbal/nonverbal responses). This framework also provides a basis for more precise interpretation of the vast amount of creativity research data being generated.

This study tested the new domain-general hypothesis for creative thought through a large-scale activation likelihood estimation (ALE) analysis. Specifically, the ALE was used to 1) distil the core findings from functional neuroimaging studies of *<u>both</u>* the creativity and relevant clinical-cognitive neuroscience domains (i.e., semantic cognition, controlled episodic memory retrieval, and executive mechanisms), and then 2) formally converge and contrast the activation maps arising in relation to creativity and the collection of primary cognitive functions. By simultaneously mapping the neural bases of creativity and the cognitive cornerstones (i.e., “pre-existing knowledge” and “executive mechanisms”), it is possible to evaluate how much of creative thought reflects the collective action of domain-general cognitive processes (reflected by considerable overlap in the brain areas activated) vs. how much neurocognitive machinery is specific to creative thought (reflected in “selectively” active regions).

## 2. Methods

### 2.1 Inclusion/exclusion criteria

We included published fMRI or positron emission topography (PET) studies, that 1) involved healthy adult participants (mean age 18-40) actively performing cognitive tasks that tap creativity, semantic cognition, controlled episodic memory retrieval or executive mechanisms, 2) conducted whole-brain analysis on specific contrasts, with 3) statistical significant [threshold p ≤ .005; (Radua et al., 2012)] results reported as coordinates in standardized stereotaxic space (Talairach or MNI).

Within the creativity domain, we included studies that reported brain activations associated with two types of tasks: 1) novel response generation tasks, which instruct participants to generate multiple creative ideas (e.g., generating multiple creative responses to a daily object or nonverbal stimulus, creating stories/music, rhythmic pattern improvisation, imagining pictures based on a cue, or generating creative responses by recombining multiple verbal or nonverbal stimuli into meaningful words or objects); and 2) novel idea selection tasks, which instruct participants to select and produce the most novel and appropriate idea (e.g., creatively generating a metaphor or word based on one or more verbal or nonverbal stimuli, generating creative solutions to verbally or nonverbally presented problems, or deciding if two verbal or nonverbal stimuli can be meaningfully combined). We included both experimental and naturalistic tasks. The following contrasts were included: 1) generating the most/multiple creative response(s) > retrieving learned response(s), and 2) generating the most/multiple creative response(s) > retrieving learned response(s) > low-level baseline.

Within the cognitive cornerstones, we defined the tasks and contrasts to be included for each component based on previous literature (**Table 1**). It is worth noting that while the same task (e.g., the semantic fluency task) could appear in different domains (e.g., semantic representation and energization), the contrasts of interest differ. For example, in semantic representation, the primary contrast is semantic > nonsemantic (e.g., semantic fluency > phonological fluency) whereas for energization, the primary contrast is active response generation > word repetition (e.g., semantic fluency > repeat the word ‘nothing’).

**Table 1:**
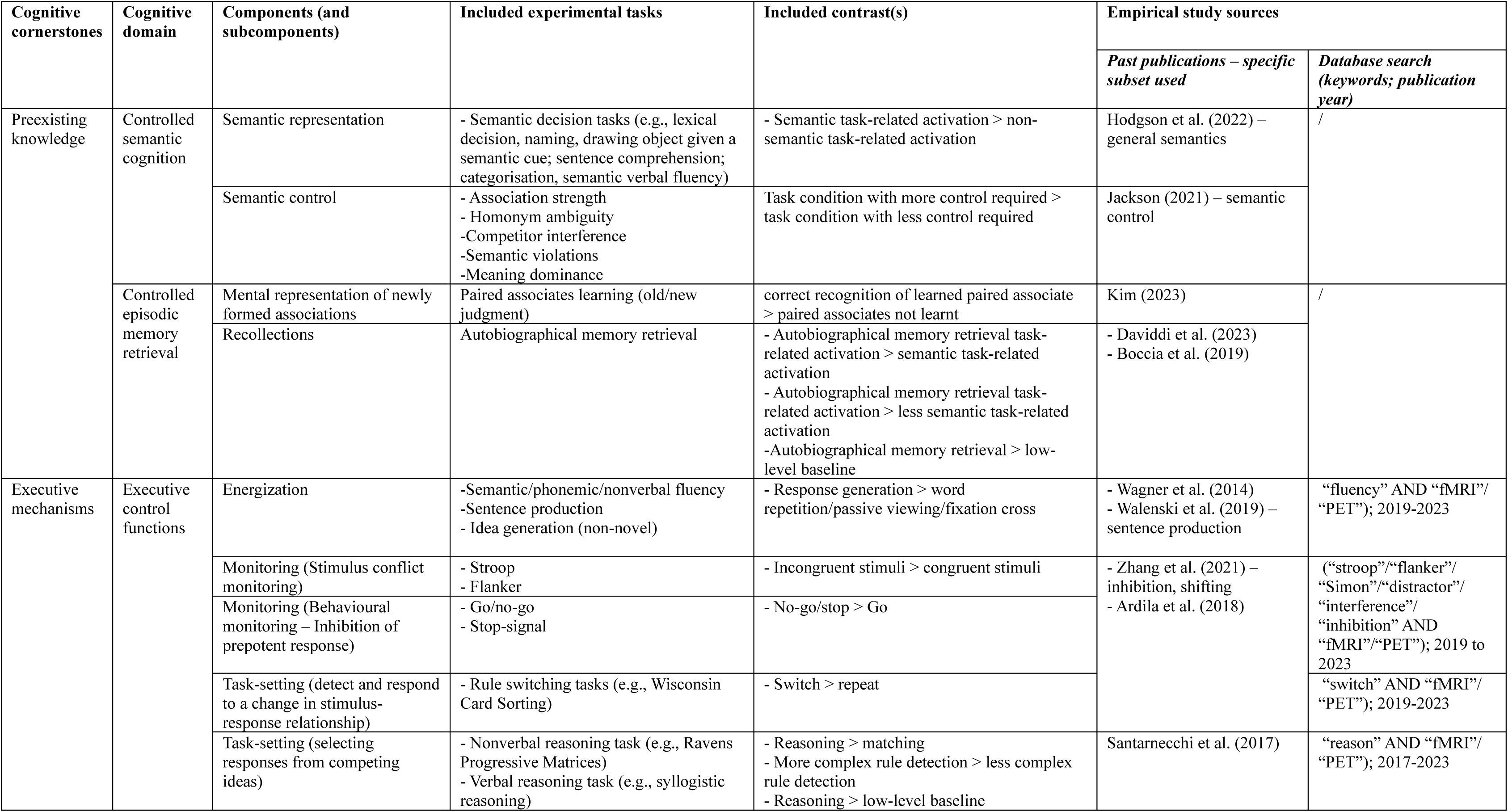
Clinical-cognitive neuroscience tasks and contrasts included in the meta-analysis.

To adjust for multiple contrasts derived from the same set of participants that could create dependence across experiment maps, coordinates generated from the same task or reflecting highly similar processes (e.g., autobiographical memory retrieval > math task across several TR frames) within the same sample were pooled into one experiment (Muller et al., 2018).

### 2.2 Study identification

Creativity studies (**Table S1**) were sourced from three previous ALE meta-analyses (Boccia et al., 2015; Gonen-Yaacovi et al., 2013; Kuang et al., 2022) and electronic database (Embase, MEDLINE, Web of Science, PsycINFO) searches were conducted with the following keywords: “creative* AND (process* OR thought* OR think* OR executive OR semantic OR cognit* OR control) AND (brain OR neural OR neuroscien* OR neurolog*)”, which were generated using a text mining software, PubReminer (O’Mara-Eves et al., 2015). No limitations were set on publication dates. Empirical studies supporting the two cognitive cornerstones of creative thought – “pre-existing knowledge” (**Table S2**) and “executive mechanisms” (**Table S3**) – were sourced from ten neuroimaging meta-analyses (Ardila et al., 2018; Boccia et al., 2019; Daviddi et al., 2023; Hodgson et al., 2023; Jackson, 2021; Kim, 2023; Santarnecchi et al., 2017; Wagner et al., 2014; Walenski et al., 2019; Zhang et al., 2021), and for executive mechanisms studies, additional database searches were performed using Google Scholar. As executive mechanisms involve multiple cognitive components, different keywords were used to identify studies that examined individual components (see **Table 1**). During Google Scholar keyword searches, the first 300 results yielded were screened (Haddaway et al., 2015).

The sources of empirical studies for each component are summarised in **Table 1**. With the inclusion/exclusion criteria applied, a total of 787 non-duplicated experiments (with 10,357 foci) representing 17,228 individuals were included in the analyses (**Figure 1a**).

**Figure 1.**
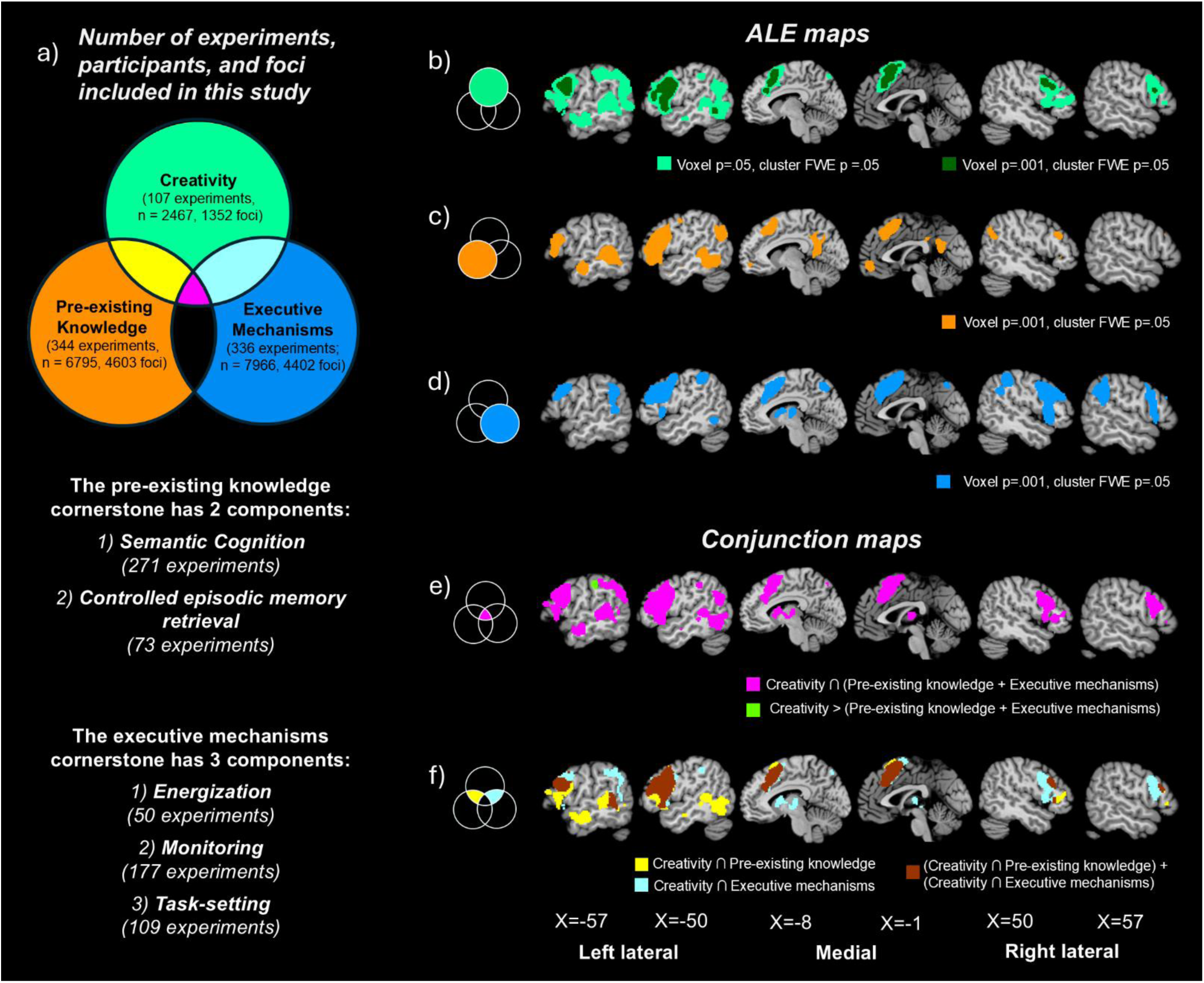
Panel a summarises the number of experiments and participants included in each domain. To address the study’s main aim, we primarily focussed on understanding the intersections of creativity and the two cognitive cornerstones – pre-existing knowledge, and executive mechanisms by conducting contrast analyses between domains. **Panels b to d are the ALE maps of each task type.** Panel b shows that creativity tasks consistently recruited fronto-parieto-temporal brain regions; these brain regions are also implicated in more domain general cognitive mechanisms, including c) controlled retrieval of pre-existing knowledge (i.e., semantic cognition and episodic memory combined) and d) executive mechanisms. Note that for panel b, we show the brain regions more likely to be activated by creativity tasks at two significance thresholds (voxel-level p = .001; dark green-shaded regions and p=.05; light green-shaded regions). **Panels e and f are the conjunction maps (voxel-level p = .001, cluster FWE p = .05).** In panel e, the magenta shaded regions show that the brain regions implicated in creativity tasks heavily overlap with those implicated in the cognitive cornerstones (i.e., pre-existing knowledge and executive mechanisms). The green-shaded region highlights the very limited number of small brain area significantly more likely to be activated during creativity tasks compared to cognitive neuroscience tasks tapping the cognitive cornerstones (see images available online for the other very small clusters reported in the results section). In panel f, the yellow-shaded regions denote the brain activation overlap between creativity and the pre-existing knowledge cornerstone; the aqua-shaded regions denote the brain activation overlap between creativity and the executive mechanisms cornerstone; the brown-shaded regions represent the overlap of both conjunction maps.

### 2.3 ALE meta-analysis

To map the brain regions of creativity and the two cognitive cornerstones of creative thought, three ALE maps were generated. These maps reveal the brain regions more likely to be recruited during tasks tapping creativity, as well as those engaged by tasks tapping the cognitive cornerstone components. To identify the overlapping neural bases of creativity and the cognitive cornerstones (see **Figure 1a**), conjunction analyses were performed to identify brain regions commonly recruited during i) creativity tasks and the cognitive cornerstones when combined (magenta-shaded intersection), ii) creativity tasks and the pre-existing knowledge cornerstone (yellow-shaded intersection) as well as iii) creativity tasks and the executive mechanisms cornerstone (aqua-shaded intersection). In addition, subtraction analyses (i.e., creativity >/< [Pre-existing knowledge + executive mechanisms] contrasts) were conducted, which denote brain regions that were more/less likely to be recruited during creativity tasks compared to the pre-existing knowledge/executive mechanisms cornerstone.

ALE, conjunction and subtraction analyses were performed using command-line GingerALE version 3.0.2 (Eickhoff et al., 2012; Eickhoff et al., 2017; Eickhoff et al., 2009; Turkeltaub et al., 2002). Data conversion was performed on the extracted Talairach space coordinate data before ALE meta-analysis. They were transformed to MNI space using the Lancaster transform (Lancaster et al., 2007), such that all analyses could be performed in MNI space at the whole-brain level. To ensure the activation probabilities represent true convergence but not random clustering, a Monte-Carlo permutation test (with 10000 permutations) was performed for all ALE analyses to remove nonsignificant clusters and to determine minimal cluster size to achieve statistical significance at family-wise error (FWE) corrected p = .05. For the generation of ALE maps that represent each cognitive cornerstone/component, the ALE values were thresholded at voxel level p = .001 for cluster-forming. Given the creativity dataset is highly heterogeneous (due to a great variety of naturalistic and experimental tasks used that probably recruits vastly different sets of cognitive mechanisms), to avoid type II error and better comparison with previous creativity ALE meta-analyses, an additional ALE map for creativity was generated at voxel level p<.05 and cluster-level FWE p<.05 (Gonen-Yaacovi et al., 2013). When generating ALE maps of each domain for contrast (conjunction/subtraction) analyses, ALE values were thresholded at voxel level p = √0.001, and the ALE maps for the pooled data (domain 1 + domain 2) required for the contrast analyses are thresholded at voxel p=.001 with cluster FWE corrected p=.05. The ALE maps were then compared using a conservative threshold of p = .001 with a minimal cluster volume of 100mm^3^ applied for cluster extraction.

## 3. Results

### 3.1 ALE maps

The brain regions with significant likelihood of activation during creativity tasks are shown in **Figure 1b** and the peak coordinates are reported in **Table 2**. Creativity tasks yielded bilateral frontal and left parieto-temporal activations with two significant clusters – 1) the larger left-lateralised cluster (83,768mm^3^) encompassing the left superior frontal gyri (extending to dorsal anterior cingulate; BA6,8,32), bilateral middle (BA9) and inferior frontal gyri (BA47; left BA44/45), anterior insula (BA13), anterior middle temporal gyrus (BA21), fusiform gyrus (BA37), and inferior parietal cortex (BA40); and 2) the smaller right hemisphere cluster (10,445mm^3^) involving only the right middle and inferior frontal gyri. Only the left medial and lateral frontal, as well as the right lateral frontal activations survived the standard threshold of voxel-level p=.001, cluster FWE p=.05.

**Table 2:** Activation likelihood of creativity.

| Cluster number<br>(size in mm<sup>3</sup>) | Hemisphere | Name of brain region | Brodmann<br>area | Max<br>ALE<br>value | Peak MNI coordinate |  |  |
| --- | --- | --- | --- | --- | --- | --- | --- |
|  |  |  |  |  | X | Y | Z |
| 1 (83,768) | Left | Middle Frontal Gyrus | BA9 | 0.065 | -50 | 10 | 22 |
|  |  | Inferior Frontal Gyrus | BA44 | 0.063 | -50 | 10 | 20 |
|  |  |  | BA45 | 0.037 | -48 | 18 | -2 |
|  |  |  | BA47 | 0.035 | -50 | 16 | -6 |
|  |  | Insula | BA13 | 0.048 | -48 | 12 | 14 |
|  |  | Superior Frontal Gyrus | BA6 | 0.063 | -2 | 16 | 54 |
|  |  |  | BA8 | 0.049 | -4 | 24 | 42 |
|  |  |  | BA32 | 0.049 | -2 | 16 | 47 |
|  |  | Anterior Cingulate<br>(Dorsal) | BA32 | 0.048 | -4 | 24 | 41 |
|  |  | Anterior middle temporal<br>gyrus | BA21 | 0.024 | -60 | 8 | -17 |
|  |  | Fusiform Gyrus | BA37 | 0.039 | -46 | -58 | -10 |
|  |  | Inferior Parietal Cortex | BA40 | 0.049 | -60 | -30 | 38 |
| 2 (10,445) | Right | Middle Frontal Gyrus | BA9 | 0.04 | 50 | 8 | 30 |
|  |  | Inferior Frontal Gyrus | BA47 | 0.016 | 34 | 36 | -24 |

Significance Threshold: voxel $p=.05$ , cluster formation at FWE corrected $p=.05$ ; Coordinates with bold text = significant at voxel $p=.001$ , cluster-FWE $p=.05$

The brain regions implicated in the pre-existing knowledge cornerstone are shown in **Figure 1c** and the peak coordinates are reported in **Table 3**. The pre-existing knowledge cognitive cornerstone recruited left-lateralised fronto-parieto-temporal brain regions, including left superior frontal gyri (extending to dorsal anterior cingulate and medial frontal pole, BA6,8,32,10), bilateral middle (BA9/46) and inferior frontal (BA44,45,47) gyri with a much more extensive cluster in the left hemisphere including bilateral anterior insula (BA13), left temporal pole (BA38), left posterior middle temporal gyrus (BA22 - extending to the fusiform gyrus BA37, bilateral parahippocampus BA28, and amygdala), left posterior cingulate (BA23) and left angular gyrus (BA39).

**Table 3:**
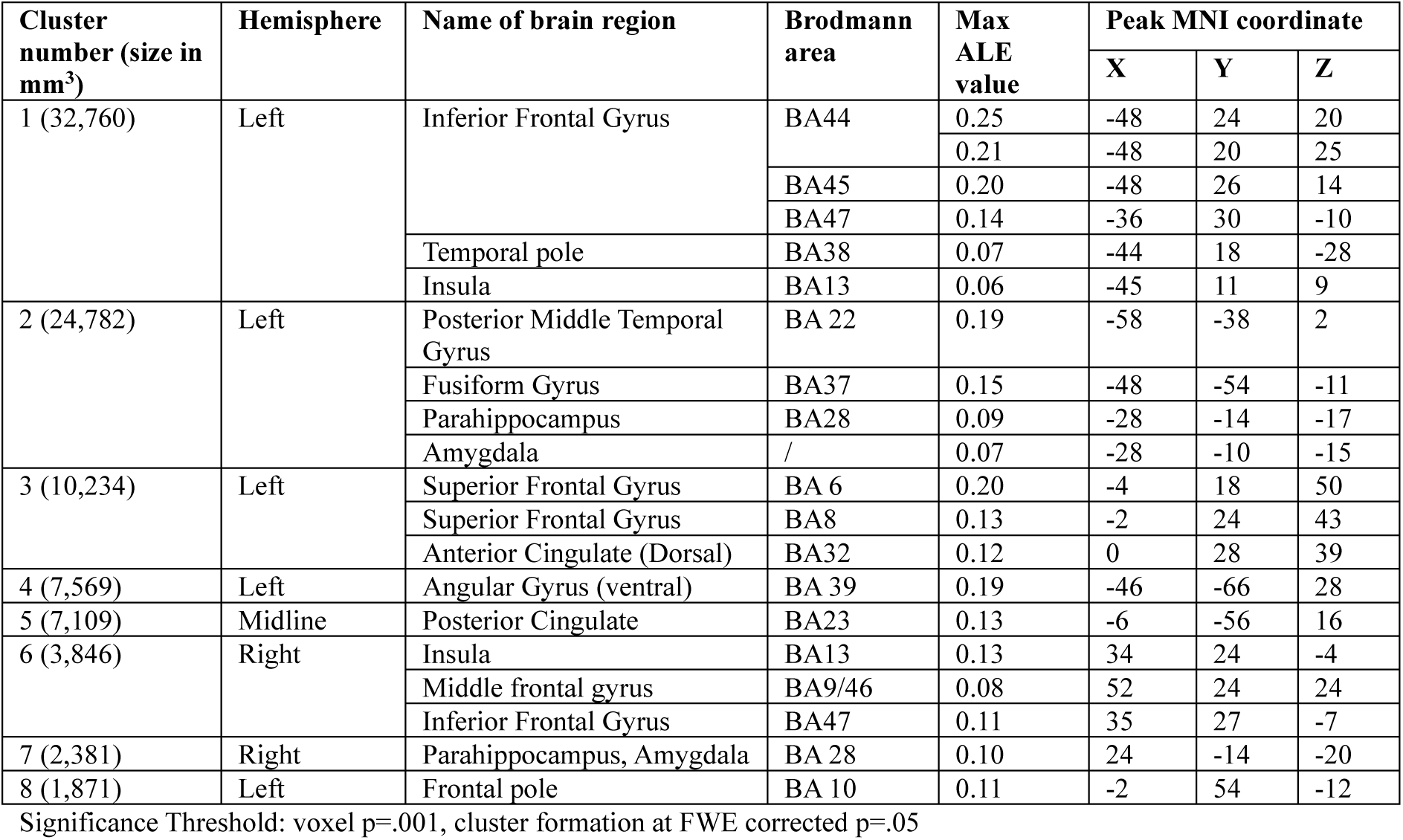
Activation likelihood of the “Pre-existing Knowledge” cornerstone.

The brain regions implicated in the executive mechanisms cornerstone are shown in **Figure 1d** and the peak coordinates are reported in **Table 4**. The executive mechanisms cognitive cornerstone primarily recruited bilateral frontoparietal brain regions, including bilateral superior frontal gyri BA6 (extending to dorsal anterior cingulate, BA32), left middle (BA9/46) and inferior (BA45; *pars triangularis*) frontal gyri, right inferior frontal gyrus *pars orbitalis* (BA47) extending to anterior insula (BA13) and lateral frontal pole (BA10), and bilateral supramarginal gyrus of the inferior parietal lobule (BA40). Left thalamus, fusiform gyrus (BA37), and right middle/inferior occipital gyrus (BA18/19) were also recruited.

**Table 4:**
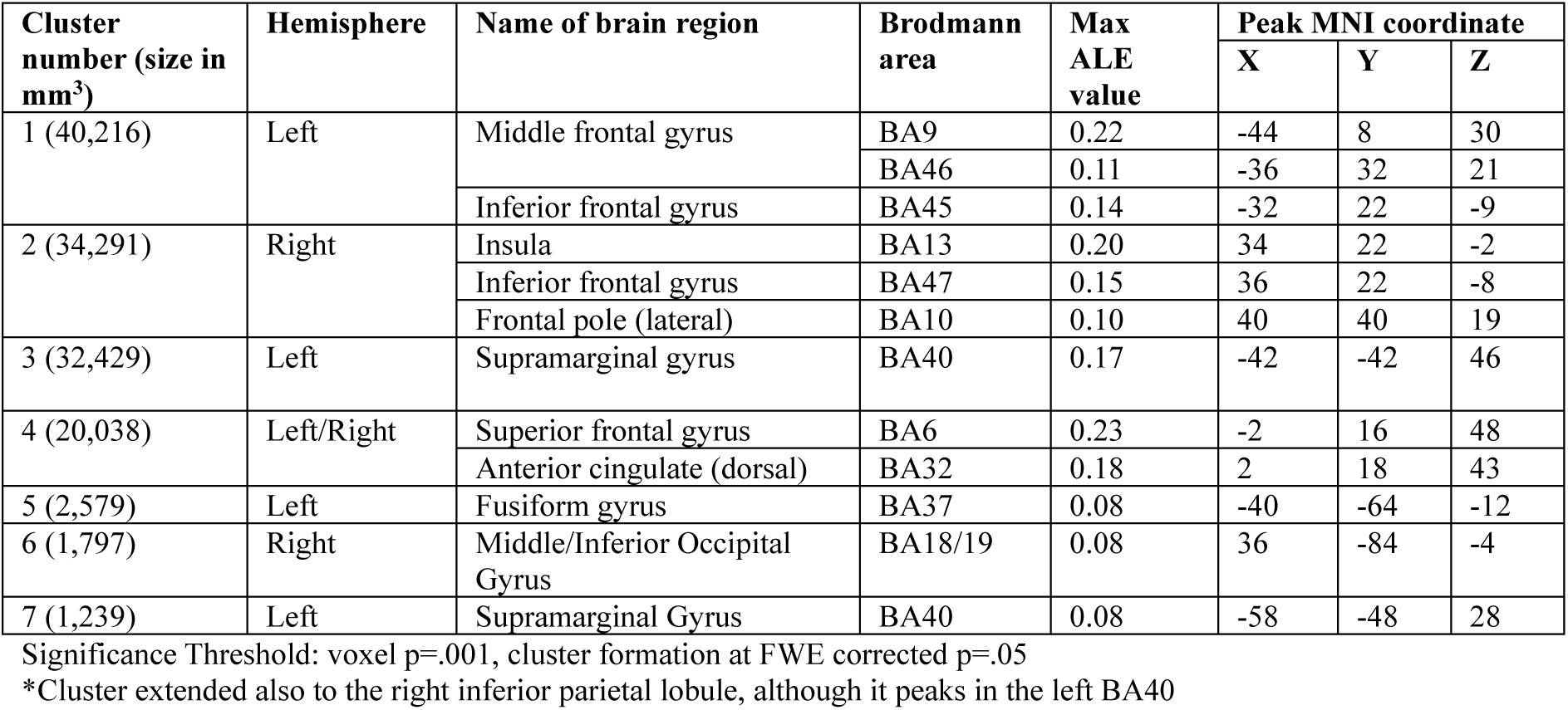
Activation likelihood of the “Executive Mechanisms” cornerstone.

| Cluster number (size in mm<sup>3</sup>) | Hemisphere | Name of brain region | Brodmann area | Max ALE value | Peak MNI coordinate |  |  |
| --- | --- | --- | --- | --- | --- | --- | --- |
|  |  |  |  |  | X | Y | Z |
| 1 (40,216) | Left | Middle frontal gyrus | BA9 | 0.22 | -44 | 8 | 30 |
|  |  |  | BA46 | 0.11 | -36 | 32 | 21 |
|  |  | Inferior frontal gyrus | BA45 | 0.14 | -32 | 22 | -9 |
| 2 (34,291) | Right | Insula | BA13 | 0.20 | 34 | 22 | -2 |
|  |  | Inferior frontal gyrus | BA47 | 0.15 | 36 | 22 | -8 |
|  |  | Frontal pole (lateral) | BA10 | 0.10 | 40 | 40 | 19 |
| 3 (32,429) | Left | Supramarginal gyrus | BA40 | 0.17 | -42 | -42 | 46 |
| 4 (20,038) | Left/Right | Superior frontal gyrus | BA6 | 0.23 | -2 | 16 | 48 |
|  |  | Anterior cingulate (dorsal) | BA32 | 0.18 | 2 | 18 | 43 |
| 5 (2,579) | Left | Fusiform gyrus | BA37 | 0.08 | -40 | -64 | -12 |
| 6 (1,797) | Right | Middle/Inferior Occipital Gyrus | BA18/19 | 0.08 | 36 | -84 | -4 |
| 7 (1,239) | Left | Supramarginal Gyrus | BA40 | 0.08 | -58 | -48 | 28 |

Significance Threshold: voxel $p=.001$ , cluster formation at FWE corrected $p=.05$
\*Cluster extended also to the right inferior parietal lobule, although it peaks in the left BA40

### 3.2 Conjunction maps

The brain regions commonly recruited by creativity and the two cognitive cornerstones combined are shown in **Figure 1e** and the peak coordinates are reported in **Table 5**. The areas in common were considerable, covering all of the regions obtained for the creative thought ALE map. This included the bilateral frontal regions encompassing bilateral middle and inferior frontal gyri (BA9,44,45,46,47) with the right hemisphere cluster extending into the anterior insula (BA13), left superior frontal gyrus involving dorsal anterior cingulate (BA6,8,32), left-lateralised temporo-parieto-occipital regions encompassing the left precuneus (BA7), left inferior parietal cortex including angular gyrus (BA39,40), left posterior middle/superior temporal gyri (BA22/37), superior/middle occipital gyri (BA19), as well as left subcortical regions (i.e., thalamus, caudate, globus pallidus and parahippocampus/amygdala). In contrast to the conjunction analyses, the subtraction analysis only revealed small left-lateralised clusters that were more likely to be recruited in creativity tasks than in semantic cognition/episodic memory/executive functions tasks, which included the left inferior parietal lobule (cluster size: 560mm^3^, BA40; peak: -59,-28,37; green-shaded cluster in **Figure 1e**), left amygdala (cluster size: 264mm^3^, peak: -28, -8, -13), left superior frontal gyrus (cluster size: 240mm^3^, BA6; peak: -20, 19, 48), left inferior frontal gyrus (cluster size: 128mm^3^, BA6/9, peak: -54, 2, 28) and left ventral angular gyrus (cluster size: 120mm^3^, BA39; peak: -48, -54,22). The reverse contrast, (Pre-existing knowledge + executive mechanisms) > creativity, revealed that the right anterior insula (cluster size: 848mm^3^, BA13; peak: 33, 21, -1), right thalamus (cluster size: 320mm3; peak: 14, -8, 10), left inferior frontal gyrus *par orbitalis* (cluster size: 312mm3, BA47; peak: -48, 30, -16) and posterior cingulate gyrus (cluster size: 304mm3, BA23; peak: 2, -25,27) were more likely to be activated in cognitive tasks tapping controlled retrieval of pre-existing knowledge and executive mechanisms. Consistent with the *Cognitive Cornerstones Hypothesis*, when we overlayed the creativity∩pre-existing knowledge (**Table 6**) and creativity∩executive mechanisms (**Table 7**) conjunction maps (**Figure 1f**), it revealed that the two cornerstones overlapped with two subsets of the brain regions that support creative thought. The intersection of creative thought with the “pre-existing knowledge” cornerstone (yellow-shaded regions) predominantly engaged a left-lateralised frontotemporal network encompassing the left superior/inferior frontal gyri, left anterior/posterior temporal lobes, and the amygdala; the intersection with the “executive mechanisms” cornerstone (aqua-shaded regions) primarily recruited bilateral frontal, left lateral parietal, left posterior superior temporal, and thalamic regions.

**Table 5:**
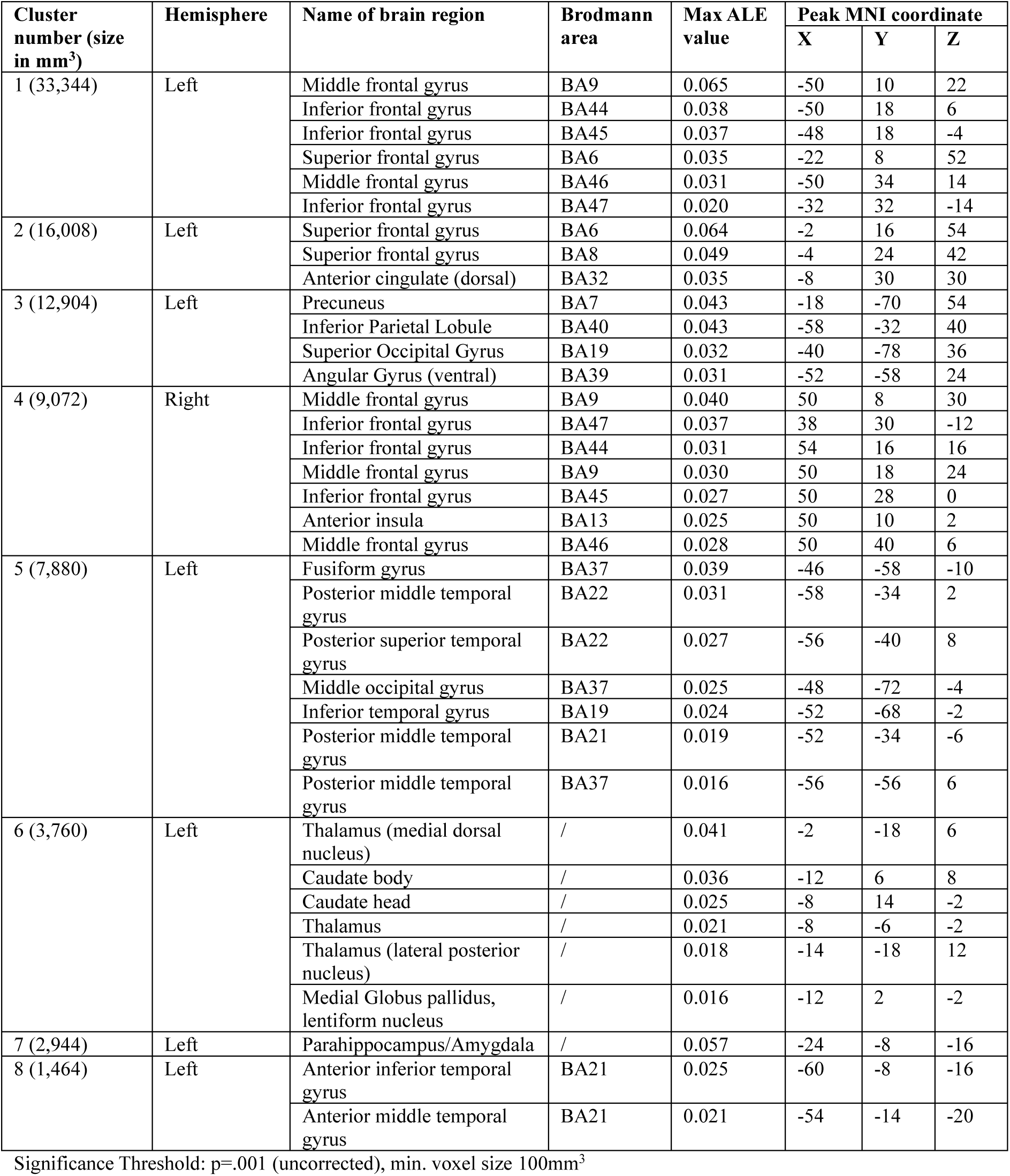
Brain regions commonly recruited by the creativity tasks, tasks tapping pre-existing knowledge, and tasks tapping executive mechanisms.

| Cluster number (size in mm<sup>3</sup>) | Hemisphere | Name of brain region | Brodmann area | Max ALE value | Peak MNI coordinate |  |  |
| --- | --- | --- | --- | --- | --- | --- | --- |
|  |  |  |  |  | X | Y | Z |
| 1 (33,344) | Left | Middle frontal gyrus | BA9 | 0.065 | -50 | 10 | 22 |
|  |  | Inferior frontal gyrus | BA44 | 0.038 | -50 | 18 | 6 |
|  |  | Inferior frontal gyrus | BA45 | 0.037 | -48 | 18 | -4 |
|  |  | Superior frontal gyrus | BA6 | 0.035 | -22 | 8 | 52 |
|  |  | Middle frontal gyrus | BA46 | 0.031 | -50 | 34 | 14 |
|  |  | Inferior frontal gyrus | BA47 | 0.020 | -32 | 32 | -14 |
| 2 (16,008) | Left | Superior frontal gyrus | BA6 | 0.064 | -2 | 16 | 54 |
|  |  | Superior frontal gyrus | BA8 | 0.049 | -4 | 24 | 42 |
|  |  | Anterior cingulate (dorsal) | BA32 | 0.035 | -8 | 30 | 30 |
| 3 (12,904) | Left | Precuneus | BA7 | 0.043 | -18 | -70 | 54 |
|  |  | Inferior Parietal Lobule | BA40 | 0.043 | -58 | -32 | 40 |
|  |  | Superior Occipital Gyrus | BA19 | 0.032 | -40 | -78 | 36 |
|  |  | Angular Gyrus (ventral) | BA39 | 0.031 | -52 | -58 | 24 |
| 4 (9,072) | Right | Middle frontal gyrus | BA9 | 0.040 | 50 | 8 | 30 |
|  |  | Inferior frontal gyrus | BA47 | 0.037 | 38 | 30 | -12 |
|  |  | Inferior frontal gyrus | BA44 | 0.031 | 54 | 16 | 16 |
|  |  | Middle frontal gyrus | BA9 | 0.030 | 50 | 18 | 24 |
|  |  | Inferior frontal gyrus | BA45 | 0.027 | 50 | 28 | 0 |
|  |  | Anterior insula | BA13 | 0.025 | 50 | 10 | 2 |
|  |  | Middle frontal gyrus | BA46 | 0.028 | 50 | 40 | 6 |
| 5 (7,880) | Left | Fusiform gyrus | BA37 | 0.039 | -46 | -58 | -10 |
|  |  | Posterior middle temporal gyrus | BA22 | 0.031 | -58 | -34 | 2 |
|  |  | Posterior superior temporal gyrus | BA22 | 0.027 | -56 | -40 | 8 |
|  |  | Middle occipital gyrus | BA37 | 0.025 | -48 | -72 | -4 |
|  |  | Inferior temporal gyrus | BA19 | 0.024 | -52 | -68 | -2 |
|  |  | Posterior middle temporal gyrus | BA21 | 0.019 | -52 | -34 | -6 |
|  |  | Posterior middle temporal gyrus | BA37 | 0.016 | -56 | -56 | 6 |
| 6 (3,760) | Left | Thalamus (medial dorsal nucleus) | / | 0.041 | -2 | -18 | 6 |
|  |  | Caudate body | / | 0.036 | -12 | 6 | 8 |
|  |  | Caudate head | / | 0.025 | -8 | 14 | -2 |
|  |  | Thalamus | / | 0.021 | -8 | -6 | -2 |
|  |  | Thalamus (lateral posterior nucleus) | / | 0.018 | -14 | -18 | 12 |
|  |  | Medial Globus pallidus, lentiform nucleus | / | 0.016 | -12 | 2 | -2 |
| 7 (2,944) | Left | Parahippocampus/Amygdala | / | 0.057 | -24 | -8 | -16 |
| 8 (1,464) | Left | Anterior inferior temporal gyrus | BA21 | 0.025 | -60 | -8 | -16 |
|  |  | Anterior middle temporal gyrus | BA21 | 0.021 | -54 | -14 | -20 |

Significance Threshold: $p=.001$ (uncorrected), min. voxel size 100mm<sup>3</sup>

**Table 6:** Brain regions commonly recruited by the creativity tasks and tasks tapping the “pre-existing knowledge” cornerstone.

| Cluster number (size in mm<sup>3</sup>) | Hemisphere | Name of brain region | Brodmann area | Max ALE value | Peak MNI coordinate |  |  |
| --- | --- | --- | --- | --- | --- | --- | --- |
|  |  |  |  |  | X | Y | Z |
| 1 (26,552) | Left | Middle frontal gyrus | BA9 | 0.065 | -50 | 10 | 22 |
|  |  | Inferior frontal gyrus | BA44 | 0.038 | -50 | 18 | 6 |
|  |  | Inferior frontal gyrus | BA45 | 0.037 | -48 | 18 | -4 |
|  |  | Middle frontal gyrus | BA46 | 0.031 | -54 | 34 | 14 |
|  |  | Middle frontal gyrus | BA6 | 0.027 | -44 | 8 | 46 |
|  |  | Inferior frontal gyrus | BA47 | 0.020 | -32 | 32 | -14 |
| 2 (14,176) | Left | Superior frontal gyrus | BA6 | 0.064 | -2 | 16 | 54 |
|  |  | Superior frontal gyrus | BA8 | 0.049 | -4 | 24 | 42 |
|  |  | Anterior cingulate (dorsal) | BA32 | 0.035 | -8 | 30 | 30 |
|  |  | Superior frontal gyrus | BA8 | 0.031 | -24 | 20 | 48 |
| 3 (7,744) | Left | Fusiform gyrus | BA37 | 0.039 | -46 | -58 | -10 |
|  |  | Middle temporal gyrus | BA22 | 0.031 | -58 | -34 | 2 |
|  |  | Inferior temporal gyrus | BA19 | 0.026 | -54 | -66 | 0 |
|  |  | Middle temporal gyrus | BA21 | 0.019 | -52 | -34 | -6 |
| 4 (3,560) | Left | Amygdala | / | 0.057 | -24 | -8 | -16 |
| 5 (2,144) | Left | Anterior middle/inferior temporal gyrus | BA21 | 0.025 | -60 | -8 | -16 |
|  |  | Temporal pole | BA38 | 0.024 | -54 | 2 | -14 |

Significance Threshold: $p=.001$ (uncorrected), min. voxel size 100mm<sup>3</sup>

**Table 7:** Brain regions commonly recruited by the creativity tasks and tasks tapping the “executive mechanisms” cornerstone.

| Cluster number (size in mm<sup>3</sup>) | Hemisphere | Name of brain region | Brodmann area | Max ALE value | Peak MNI coordinate |  |  |
| --- | --- | --- | --- | --- | --- | --- | --- |
|  |  |  |  |  | X | Y | Z |
| 1 (29,312) | Left | Middle frontal gyrus | BA9 | 0.065 | -50 | 10 | 22 |
|  |  | Inferior frontal gyrus | BA44 | 0.038 | -50 | 18 | 6 |
|  |  | Inferior frontal gyrus | BA45 | 0.037 | -48 | 18 | -4 |
|  |  | Superior frontal gyrus | BA6 | 0.036 | -22 | 10 | 50 |
|  |  | Middle frontal gyrus | BA46 | 0.031 | -50 | 34 | 14 |
| 2 (14,960) | Left | Superior frontal gyrus | BA6 | 0.064 | -2 | 16 | 54 |
|  |  | Superior frontal gyrus | BA8 | 0.049 | -4 | 24 | 42 |
|  |  | Anterior cingulate (dorsal) | BA32 | 0.035 | -8 | 30 | 30 |
| 3 (9,032) | Right | Middle frontal gyrus | BA9 | 0.040 | 50 | 8 | 30 |
|  |  | Inferior frontal gyrus | BA47 | 0.037 | 38 | 30 | -12 |
|  |  | Inferior frontal gyrus | BA44 | 0.031 | 54 | 16 | 16 |
|  |  | Middle frontal gyrus | BA9 | 0.030 | 50 | 18 | 24 |
|  |  | Inferior frontal gyrus | BA45 | 0.027 | 50 | 28 | 0 |
|  |  | Anterior insula | BA13 | 0.025 | 50 | 10 | 2 |
| 4 (8,848) | Left | Precuneus | BA7 | 0.043 | -18 | -70 | 54 |
|  |  | Angular gyrus | BA39 | 0.029 | -34 | -58 | 48 |
|  |  | Angular gyrus | BA39 | 0.022 | -54 | -54 | 26 |
|  |  | Inferior parietal lobule | BA40 | 0.027 | -38 | -52 | 46 |
|  |  | Inferior parietal lobule | BA40 | 0.028 | -54 | -34 | 40 |
| 5 (1,880) | Left | Caudate body | / | 0.036 | -12 | 6 | 8 |
|  |  | Caudate head | / | 0.025 | -8 | 14 | -2 |
|  |  | Thalamus (ventral anterior nucleus) | / | 0.016 | -8 | -4 | 4 |
| 6 (1,592) | Left | Thalamus (medial dorsal nucleus) | / | 0.039 | -4 | -18 | 6 |
|  |  | Thalamus (lateral posterior nucleus) | / | 0.018 | -14 | -18 | 12 |
| 7 (728) | Left | Posterior superior temporal gyrus | BA22 | 0.025 | -58 | -42 | 10 |

Significance Threshold: $p=.001$ (uncorrected), min. voxel size 100mm<sup>3</sup>

## 4. Discussion

This large-scale ALE analysis formally compared and contrasted the brain activation maps implicated in creativity and two potentially crucial cognitive cornerstones (i.e., pre-existing knowledge and executive mechanisms) that underpin creative thought. The conjunction analysis revealed striking and considerable overlap between the brain regions implicated in creativity and cognitive cornerstones of creative thought (**Figure 1e**; magenta-shaded regions). This result implies that creative thought is not a distinct cognitive function [as proposed by some creativity researchers e.g., Beaty et al. (2023); Benedek et al. (2023); Benedek and Fink (2019); Kenett and Faust (2019); Kleinmintz et al. (2019); Lebuda and Benedek (2023); Lin and Vartanian (2018)]. Instead, it seems more accurate to view creative thought functions as arising from the dynamic interplay between more domain-general cognitive cornerstones, with these cornerstones exhibiting both shared and distinct neural correlates.

The conjunction analyses between the brain maps of creativity and each cornerstone, (creativity *∩* pre-existing knowledge) and (creativity *∩* executive mechanisms), revealed three overlapping clusters – left middle/inferior frontal gyrus, left superior/medial frontal gyrus, and left lateral inferior temporal gyrus (**Figure 1f**; brown-shaded regions). This result suggests that these regions are involved in shared cognitive processes including but not limited to creative activities, consistent with the multiple-demand network hypothesis (Duncan, 2010), which highlights that these brain regions are consistently coactivated during all demanding cognitive tasks. What are the possible cognitive processes associated with these brain regions? Neuropsychological and neuroimaging findings suggest that these brain regions are implicated in a number of cognitive processes. The left lateral frontal cortex (encompassing both middle and inferior frontal gyri) is associated with the selection of task/goal-relevant representations amongst competing ideas (Stuss, 2011), with the dorsal part (left middle frontal gyrus) being associated with resolving task response conflict during unexpected conditions (Badre, 2008), and the ventral-anterior part (left inferior frontal gyrus) implicated in semantic control and controlled episodic memory retrieval (Vatansever et al., 2021). The left superior/medial frontal gyrus is implicated in self-initiated responses (Robinson et al., 2012) and estimation of cognitive effort (Hauser et al., 2017), while the left lateral posterior temporal cortex (BA37) is associated with semantic control (Hodgson et al., 2023) and executive control processes (Zhang et al., 2021). To explore these regions’ roles in creative activities in more detail and anatomical precision, future research based on new fMRI comparisons across tasks within the same participants will be needed, as recent studies have done for the middle/inferior frontal gyri (Assem et al., 2024; Chiou et al., 2025) and inferior parietal cortex (Humphreys et al., 2024).

Strikingly the subtraction analysis (creativity > cognitive cornerstones contrast) only identified a handful of small left-lateralised frontoparietal and subcortical clusters. These clusters included the inferior parietal lobule (BA39/40), precuneus (BA7), lateral superior/middle frontal gyrus (BA6/9), and amygdala. These small activation clusters either were extensions of the large overlapping brain regions or appeared as small hot spots within (much larger) overlapping clusters recruited by tasks tapping both creativity and the cognitive cornerstones. For example, the anteriorly extended BA39/40 cluster (**Figure 1e**; green-shaded region) might imply that creativity tasks engage a larger neuronal population supporting autobiographical memory retrieval and buffering of multimodal contextual information (Humphreys et al., 2024). These findings converge to emphasise that creativity tasks do not uniquely engage additional brain regions but instead recruit the very same networks activated by tasks tapping semantic cognition, episodic memory retrieval, and executive functions.

To the best of our knowledge, this is the largest ALE meta-analysis ever conducted to examine the overlap between brain networks implicated in creativity and the two possible neurocognitive mechanisms underpinning creative thought. Our findings provided powerful support for the *Cognitive Cornerstones Hypothesis* of creative thought, which posits that: 1) creative thought indeed comprises two cognitive cornerstones, pre-existing knowledge (i.e., the reservoir from which ideas are drawn from and synthesised for further processing) and executive mechanisms (i.e., the cognitive processes that allow goal-directed, flexible, and transformative use of preexisting knowledge); and 2) creative thought arises from general purpose cognitive mechanisms supporting semantic cognition, episodic memory, and executive control functions. Despite its strengths, this investigation has two limitations. First, activation-based analyses cannot identify the brain regions critical for a specific cognitive process. To determine the regions essential for creative thought, alternative methods, such as neuropsychological lesion studies and brain stimulation techniques, are required. Second, our meta-analysis revealed an intrinsic bias in the current literature: although we attempted to compile a balanced dataset generated from both verbal and nonverbal tasks across all domains, the majority of creativity tasks (75.8%; **Table S1**) and tasks tapping the pre-existing knowledge cornerstone (79.0%; **Table S2**) are verbal, whereas most tasks tapping executive mechanisms are nonverbal (56.8%; **Table S3**). Nevertheless, the findings from our meta-analysis can initiate and facilitate the crosstalk between creativity and clinical-cognitive neuroscience. By grounding creative thought in the cognitive cornerstones of semantic cognition, episodic memory retrieval, and executive mechanisms, we move away from treating creativity as a distinct cognitive function and toward understanding it as a dynamic capacity emerging from interactions among foundational cognitive processes.

## 5. Conclusion

Our large-scale functional neuroimaging meta-analysis suggests that the experimental tasks tapping creativity recruit frontoparietotemporal regions, which overlap with most of the brain regions implicated in fundamental cognitive components – semantic cognition, controlled episodic memory retrieval, and executive control functions. These findings provide powerful support for the *Cognitive Cornerstones Hypothesis* of creative thought, which posits that creative thought might not be a distinct mental capacity, but instead arises from the interplay of general purpose cognitive cornerstones (i.e., pre-existing knowledge and executive mechanisms). Future studies may explore how the cognitive cornerstone components play differential roles in supporting the diverse phenomena of creativity, from Monet’s impressionism and Einstein’s theories, to how we devise new solutions that solve everyday life problems, or why a release of creativity is reported to occur in some people with neurological disorders.

## 6. Data and code availability

All datasets and images used in this manuscript are available at https://github.com/melodymychan/CreativeThoughtALE.git.

## 7. Author contributions

M.Y.M.C.: conceptualisation, methodology, formal analysis, software, visualisation, writing – original draft; M.A.L.R.: conceptualisation, methodology, supervision, funding acquisition, writing – review and editing; G.A.R.: conceptualisation, supervision, funding acquisition, writing – review and editing.

## 8. Funding

This research is supported by the Australian Research Council grant (DP220103941) awarded to GAR and MALR. MALR is also supported by intramural funding from the UKRI-MRC (MC_UU_00005/18).

## 9. Declaration of competing interests

The authors declared no competing interests.

## 10. Acknowledgements

The authors thank Dr. Victoria Hodgson for her input in data analysis and Ms. Eugene Cho for her input in collecting and checking the data used in the analyses.

## Notes

https://github.com/melodymychan/CreativeThoughtALE.git

### Competing Interest Statement

The authors have declared no competing interest.

