## Supplementary material for "The neural basis of creative thought: An activation likelihood estimation meta-analysis involving over 17,000 participants": Table S1

**Table S1: Contrasts included in the creativity dataset**

| Reference | Contrast | DOI/Link | N | Foci | Task contrast | Input modality |
|---|---|---|---|---|---|---|
| Abraham et al., 2012 | 1 | https://doi.org/10.1016/j.neuropsychologia.2012.04.015 | 19 | 35 | AUT: divergent thinking contrast (Table S1) | Verbal |
| Asari et al., 2008 | 1 | 10.1016/j.neuroimage.2008.01.059 | 68 | 4 | Rorschach: unique > frequent | Nonverbal |
| Aziz-Zadeh et al., 2009 | 1 | 10.1002/hbm.20554 | 10 | 8 | Anagram: aha > non-Aha | Verbal |
| Aziz-Zadeh et al., 2013 | 1 | 10.1093/scan/nss021 | 13 | 10 | combine shapes into meaningful objects: creative > basic shapes | Nonverbal |
| Bechtereva et al., 2004 | 1 | https://doi.org/10.1016/j.ijpsycho.2004.01.001 | 16 | 7 | Create stories from 16 remotely associated words > memorize words | Verbal |
| Bengtsson et al., 2007 | 1 | 10.1162/jocn.2007.19.5.830 | 11 | 14 | music improvisation > play from memory | Nonverbal |
| Berkowitz and Ansari, 2008 | 1 | 10.1016/j.neuroimage.2008.02.028 | 13 | 13 | music improvisation: melodic improvisation > patterns | Nonverbal |
| Berkowitz and Ansari, 2008 | 2 | 10.1016/j.neuroimage.2008.02.028 | 13 | 7 | music improvisation: rhythm improvisation > metronome | Nonverbal |
| Cardillo et al., 2012 | 1 | https://doi.org/10.1016/j.neuroimage.2011.11.079 | 20 | 4 | metaphor comprehension: novel > familiar | Verbal |
| Chrysikou and Thompson-Schill 2011 | 1 | 10.1002/hbm.21056 | 24 | 3 | uncommon uses for daily objects: unusual > usual | Verbal |
| de Manzano and Ullen, 2012 | 1 | 10.1016/j.neuroimage.2011.07.016 | 18 | 8 | Music improvisation in classical pianists > music reading | Nonverbal |
| Ellamil et al., 2012 | 1 | https://doi.org/10.1016/j.neuroimage.2011.08.008 | 15 | 16 | Design book cover given a summary paragraph: generate > evaluate | Verbal |
| Fink et al., 2010 | 1 | 10.1016/j.neuroimage.2010.05.072 | 31 | 1 | alternative uses > object characteristics | Verbal |
| Fink et al., 2009 | 2 | 10.1002/hbm.20538 | 21 | 1 | alternative uses > object characteristics | Verbal |
| Fink et al., 2009 | 1 | 10.1002/hbm.20538 | 21 | 10 | alternative uses > fixation | Verbal |
| Geake and Hansen, 2005 | 1 | 10.1016/j.neuroimage.2005.01.035 | 12 | 15 | letter string analogy: greater analogical depth > lesser analogical depth | Verbal |
| Goel and Vartanian, 2005 | 1 | 10.1093/cercor/bhh217 | 13 | 3 | Conjunction: (Match problems task > baseline) and (successful trial - unsuccessful trial) | Nonverbal |
| Green et al., 2012 | 1 | 10.1037/a0025764 | 23 | 10 | Analogy: more remote > less remote | Verbal |
| Green et al., 2012 | 2 | 10.1037/a0025764 | 23 | 5 | Analogy: generation > rest | Verbal |
| Howard-Jones et al., 2005 | 1 | 10.1016/j.cogbrainres.2005.05.013 | 8 | 4 | Story generation from 3 words: creative > uncreative | Verbal |
| Huang et al., 2012 | 1 | 10.1002/hbm.22093 | 26 | 3 | Imagination of pictures: creative > uncreative | Nonverbal |

| Reference | Contrast | DOI/Link | N | Foci | Task contrast | Input modality |
|---|---|---|---|---|---|---|
| Jung-Beeman et al., 2004 | 1 | https://doi.org/10.1371/journal.pbio.0020097 | 18 | 7 | CRAT: Aha > no Aha | Verbal |
| Kounios et al., 2006 | 1 | 10.1111/j.1467-9280.2006.01798.x | 25 | 6 | CRAT: Aha > no Aha | Verbal |
| Kowatari et al., 2009 | 1 | 10.1002/hbm.20633 | 20 | 11 | Design of a new tool by expert > counting | Nonverbal |
| Kowatari et al., 2009 | 1 | 10.1002/hbm.20633 | 20 | 8 | Design of a new tool by novice > counting | Nonverbal |
| Kroeger et al., 2012 | 1 | http://doi.org/10.1016/j.brainres.2011.10.031 | 19 | 25 | Semantic judgement: judge whether the word pairs are unusual and appropriate | Verbal |
| Limb and Braun, 2008 | 1 | 10.1371/journal.pone.0001679 | 6 | 57 | Music improvisation > rest | Nonverbal |
| Luo et al., 2004b | 1 | https://doi.org/10.1007/978-3-540-68044-4_15 | 15 | 12 | insight problem metaphor comprehension Aha > no aha | Verbal |
| Mashal et al., 2007 | 1 | 10.1016/j.bandl.2005.10.005 | 15 | 3 | Semantic judgement: novel metaphor > unrelated words | Verbal |
| Qiu et al., 2010 | 1 | https://doi.org/10.1016/j.cortex.2009.06.006 | 16 | 19 | Chinese Logogriphs: Aha > no-Aha | Verbal |
| Rutter et al., 2012 | 1 | http://doi.org/10.1016/j.bandc.2011.11.002 | 18 | 7 | Semantic judgement: judge whether the word pairs are unusual and appropriate (conjunction for conceptual expansion; Table 3) | Verbal |
| Seger et al., 2000 | 1 | https://doi.org/10.1037/0894-4105.14.3.361 | 7 | 10 | Noun-verb generation: novel > repeated | Verbal |
| Seger et al., 2000 | 2 | https://doi.org/10.1037/0894-4105.14.3.361 | 7 | 14 | Noun-verb generation: unusual > first associate | Verbal |
| Shah et al., 2013 | 1 | 10.1002/hbm.21493 | 28 | 28 | Creative story writing: writing > fixation | Verbal |
| Shah et al., 2013 | 2 | 10.1002/hbm.21493 | 28 | 31 | Creative story thinking: thinking > fixation | Verbal |
| Sieborger et al., 2007 | 1 | http://doi.org/10.1016/j.brainres.2007.05.079 | 14 | 1 | Making inferences: much imagination required for seeing the connection between 2 sentences > unrelated sentences | Verbal |
| Sieborger et al., 2007 | 2 | http://doi.org/10.1016/j.brainres.2007.05.079 | 14 | 10 | Making inferences: 2 sentences that required imagination to see a connection between them > 2 semantically related sentences | Verbal |
| Tian et al., 2011 | 1 | 10.1016/j.bbr.2010.09.005 | 16 | 7 | Chinese Logogriphs: successful > unsuccessful problem solving | Verbal |
| Vartanian and Goel, 2005 | 1 | 10.1016/j.neuroimage.2005.05.016 | 15 | 11 | Anagrams: unconstrained > semantically constrained | Verbal |
| Vartanian and Goel, 2005 | 2 | 10.1016/j.neuroimage.2005.05.016 | 15 | 10 | Anagrams: unconstrained > baseline | Verbal |
| Liu et al., 2012 | 1 | 10.1038/srep00834 | 12 | 19 | Lyrics Improvisation > recite overlearned lyrics | Verbal |

| Reference | Contrast | DOI/Link | N | Foci | Task contrast | Input modality |
|---|---|---|---|---|---|---|
| Villarreal et al., 2013 | 1 | 10.1371/journal.pone.0075427 | 24 | 4 | Create rhythmic phrase > repeat rhythmic phrase (high creative) | Nonverbal |
| Villarreal et al., 2013 | 1 | 10.1371/journal.pone.0075427 | 24 | 2 | Create rhythmic phrase > repeat rhythmic phrase (low creative) | Nonverbal |
| Abraham et al., 2014 | 1 | 10.1007/s11682-013-9241-4 | 14 | 17 | males: divergent thinking (conjunction) | Verbal |
| Abraham et al., 2014 | 1 | 10.1007/s11682-013-9241-4 | 14 | 14 | females: divergent thinking (conjunction) | Verbal |
| Benedek et al., 2014a | 1 | http://doi.org/10.1016/j.neuroimage.2013.12.046 | 28 | 8 | word production: metaphor > literal synonym | Verbal |
| Benedek et al., 2014b | 1 | http://doi.org/10.1016/j.neuroimage.2013.11.021 | 35 | 1 | AUT: old and new idea > baseline | Verbal |
| Benedek et al., 2014b | 2 | http://doi.org/10.1016/j.neuroimage.2013.11.021 | 35 | 5 | AUT: new > old | Verbal |
| Fink et al., 2012 | 1 | https://doi.org/10.1002/hbm.21387 | 32 | 1 | Generation of idea with example: Original idea example> control (example given: meaningless word) | Verbal |
| Fink et al., 2012 | 2 | https://doi.org/10.1002/hbm.21387 | 32 | 3 | Generation of idea: Original idea example> common idea example | Verbal |
| Green et al., 2015 | 1 | 10.1002/hbm.22676 | 55 | 18 | Generation of verb: creative cue > ordinary cue | Verbal |
| Zhang et al., 2014 | 1 | http://doi.org/10.1016/j.cortex.2013.01.015 | 18 | 17 | creative analogical reasoning: NBFFAT > baseline + BFFAT > baseline | Verbal |
| Abraham et al., 2018 | 1 | https://doi.org/10.1016/j.neuropsychologia.2018.05.004 | 34 | 22 | alternative uses > recall object location | Verbal |
| Beaty et al., 2017 | 1 | http://doi.org/10.1016/j.neuroimage.2017.01.012 | 24 | 4 | Low constraint verb generation > recall | Verbal |
| Beaty et al., 2017 | 2 | http://doi.org/10.1016/j.neuroimage.2017.01.012 | 24 | 6 | high constraint verb generation > recall | Verbal |
| Beaty et al., 2018 | 1 | 10.1162/jocn_a_01327 | 29 | 15 | alternate uses > sentence generation with 2 words | Verbal |
| Benedek et al., 2018 | 1 | 10.1016/j.cortex.2017.10.024 | 42 | 6 | novel object uses: create original > recall common | Verbal |
| Benedek et al., 2020 | 1 | https://doi.org/10.1016/j.neuroimage.2020.116586 | 44 | 2 | original association > common association | Verbal |
| Benedek et al., 2020 | 2 | https://doi.org/10.1016/j.neuroimage.2020.116586 | 44 | 5 | bi association > common association | Verbal |
| Benedek et al., 2020 | 3 | https://doi.org/10.1016/j.neuroimage.2020.116586 | 44 | 3 | bi association > original association | Verbal |
| Fink et al., 2015 | 1 | 10.1002/hbm.22901 | 53 | 4 | alternate uses > generate facts given an | Verbal |

| Reference | Contrast | DOI/Link | N | Foci | Task contrast | Input modality |
|---|---|---|---|---|---|---|
| | | | | | objective (T1) | |
| Ivancovsky et al., 2018 | 1 | https://doi.org/10.1002/hbm.24288 | 36 | 1 | generate alternate uses original > generate common | Verbal |
| Kleibeuker et al., 2013 | 1 | https://doi.org/10.1016/j.dcn.2013.03.003 | 43 | 15 | creative use of tools > object characteristics | Verbal |
| Kleinmintz et al., 2018 | 1 | https://doi.org/10.1007/s00429-017-1500-5 | 18 | 3 | creative use of tools > object characteristics | Verbal |
| Mayseless et al., 2015 | 1 | https://doi.org/10.1016/j.neuroimage.2015.05.030 | 30 | 2 | creative use of tools > object characteristics | Verbal |
| Sun et al., 2016 | 1 | https://doi.org/10.1002/hbm.23246 | 14 | 1 | creative use of tools > object characteristics (pre-training) | Verbal |
| Sun et al., 2019 | 1 | 10.1093/cercor/bhy010 | 29 | 1 | creative use of tools > object characteristics (pre-training) | Verbal |
| Becker et al., 2019 | 1 | https://doi.org/10.1016/j.neuroimage.2019.116294 | 27 | 2 | modified CRA trial start: Aha > no Aha | Verbal |
| Becker et al., 2019 | 2 | https://doi.org/10.1016/j.neuroimage.2019.116294 | 27 | 2 | modified CRA solution stage for insight: Aha > no Aha | Verbal |
| Hao et al., 2013 | 1 | http://doi.org/10.1016/j.brainres.2013.08.041 | 17 | 2 | heuristic prototype: highlight feature > no highlighted feature | Verbal |
| Huang et al., 2015 | 1 | http://doi.org/10.1016/j.neuroimage.2015.03.030 | 15 | 19 | Chinese word chunk decomposition: creative > not creative | Verbal |
| Huang et al., 2018 | 1 | https://doi.org/10.1016/j.neuroimage.2018.01.070 | 20 | 15 | riddle problem solving: high novelty > low novelty + high appropriateness > low appropriateness | Verbal |
| Kizilirmak et al., 2016 | 1 | https://doi.org/10.3389/fpsyg.2016.01693 | 26 | 9 | solved German CRAT > not solved | Verbal |
| Kizilirmak et al., 2019 | 1 | https://doi.org/10.1016/j.concog.2019.01.005 | 23 | 21 | solved German CRAT > not solved | Verbal |
| Lin et al., 2020 | 1 | https://doi.org/10.1007/s11682-020-00337-z | 32 | 8 | Chinese word chunk decomposition: high creative > low creative (INPS contrast) | Verbal |
| Luo and Niki, 2003 | 1 | https://doi.org/10.1002/hipo.10069 | 7 | 39 | insight problem solving (riddle) > baseline | Verbal |
| Luo et al., 2004a | 1 | https://psycnet.apa.org/doi/10.1097/00001756-200409150-00004 | 13 | 21 | cerebral gymnastics puzzles > baseline | Verbal |
| Luo et al., 2006 | 1 | http://doi.org/10.1016/j.brainresbull.2006.07.005 | 13 | 19 | Chinese word chunk decomposition: tight chunking > loose chunking | Verbal |
| Luo et al., 2013 | 1 | https://doi.org/10.1371/journal.pone.0049231 | 17 | 1 | heuristic prototype: novel innovation > old innovation | Verbal |
| Pang et al., 2009 | 1 | 10.1109/ICNC.2009.631 | 13 | 3 | Chinese word chunk decomposition: tight | Verbal |

| Reference | Contrast | DOI/Link | N | Foci | Task contrast | Input modality |
|---|---|---|---|---|---|---|
| | | | | | chunking > loose chunking | |
| Sinitsyn et al., 2020 | 1 | 10.3390/bs10110170 | 32 | 6 | insight > rest (anagram) | Verbal |
| Subramaniam et al., 2009 | 1 | https://doi.org/10.1162/jocn.2009.21057 | 27 | 4 | CRA: insight > analytical (preparation) | Verbal |
| Subramaniam et al., 2009 | 2 | https://doi.org/10.1162/jocn.2009.21057 | 27 | 7 | CRA: insight > analytical (solution) | Verbal |
| Tang et al., 2015 | 1 | https://doi.org/10.1093/cercor/bhv113 | 22 | 23 | tight chunking > loose chunking | Verbal |
| Terai et al., 2013 | 1 | https://escholarship.org/uc/item/35d004q0 | 18 | 1 | chunk > nonchunk (correct trials) RAT | Verbal |
| Tik et al., 2018 | 1 | 10.1002/hbm.24073 | 29 | 3 | high insight > low insight RAT German | Verbal |
| Tong et al., 2013 | 1 | https://doi.org/10.3724/SP.J.1041.2013.00740 | 16 | 2 | heuristic prototype: related > unrelated | Verbal |
| Wu et al., 2013 | 1 | https://doi.org/10.1002/hbm.21501 | 14 | 24 | Chinese chunk decomposition: tight (unfamiliar) > loose (unfamiliar) | Verbal |
| Yu et al., 2019 | 1 | https://doi.org/10.3758/s13415-019-00702-6 | 20 | 15 | semantic judgement: metaphor > literal | Verbal |
| Zhao et al., 2014 | 1 | https://doi.org/10.1016/j.neuroscience.2013.10.019 | 17 | 7 | aha > no aha riddle problem solving | Verbal |
| Zhou et al., 2011 | 1 | https://doi.org/10.1109/CISP.2011.6100412 | 10 | 9 | Riddle/idiom problem: target words are new and semantically related with riddle (NR) > target words are new but unrelated with the riddle (NU) | Verbal |
| Zhou et al., 2011 | 2 | https://doi.org/10.1109/CISP.2011.6100412 | 10 | 9 | Riddle/idiom problem: target words are old but can form new meanings with the riddle (ON) > target words are old and have literal semantic meaning of the riddle (OL) | Verbal |
| Amir et al., 2016 | 1 | https://doi.org/10.3389/fnhum.2016.00597 | 13 | 7 | Cartoon caption generation > describe cartoon | Verbal |
| Beaty et al., 2015 | 1 | 10.1038/srep10964 | 25 | 25 | alternate uses task > object characteristics | Verbal |
| Bitsch et al., 2021 | 1 | https://doi.org/10.1038/s41598-021-89843-8 | 24 | 28 | cartoon caption generation: creative > typical | Verbal |
| Fu et al., 2019 | 1 | 10.1017/dsj.2019.21 | 18 | 1 | design generation: no example > with example | Nonverbal |
| Liu et al., 2015 | 1 | 10.1002/hbm.22849 | 27 | 42 | Generation of new poems > recite poems | Verbal |
| Heinomen et al., 2016 | 1 | 10.1371/journal.pone.0162234 | 20 | 3 | AUT: idea generation (production phase) > object identification (retrieval phase) | Verbal |
| Goucher-Lambert et al., 2019 | 1 | https://doi.org/10.1016/j.destud.2018.07.001 | 21 | 6 | design ideation (Creative a device that solve the given problem): far + near cues > cue words are words from the problem itself | Verbal |
| Hahm et al., 2017 | 1 | https://doi.org/10.12779/dnd.2017.16.2.48 | 25 | 31 | creative drawing imagery > line tracking | Nonverbal |
| Saggar et al., 2015 | 1 | 10.1038/srep10894 | 36 | 6 | word drawing > zigZag drawing | Nonverbal |
| Erhard et al., 2014 | 1 | http://doi.org/10.1016/j.neuroimage.2014.05.076 | 20 | 13 | creative writing > reading | Verbal |

| Reference | Contrast | DOI/Link | N | Foci | Task contrast | Input modality |
|---|---|---|---|---|---|---|
| De Aquino et al., 2019 | 1 | https://doi.org/10.1038/s41598-019-49405-5 | 19 | 15 | musicians: improvise > repeat | Nonverbal |
| De Aquino et al., 2019 | 1 | https://doi.org/10.1038/s41598-019-49405-5 | 21 | 8 | nonmusicians: improvise > repeat | Nonverbal |
| Dhakal et al., 2019 | 1 | 10.1089/brain.2017.0566 | 24 | 5 | music improvisation: improvise > repeat | Nonverbal |
| Chauvigne et al., 2018 | 1 | https://doi.org/10.1371/journal.pone.0191098 | 19 | 16 | improvised action > non-improvised action | Nonverbal |
| Donnay et al., 2014 | 1 | https://doi.org/10.1371/journal.pone.0088665 | 11 | 17 | Jazz improvisation: improvised > recited | Nonverbal |
| Vartanian et al., 2018 | 1 | https://doi.org/10.1016/j.neuropsychologia.2018.02.024 | 44 | 14 | alternate uses > object characteristics | Verbal |
| Becker et al., 2020 | 1 | https://doi.org/10.1016/j.neuroimage.2019.116294 | 30 | 9 | CRAT: greater semantic distance solution > lower semantic distance solution | Verbal |
| Wu et al., 2021a | 1 | https://psycnet.apa.org/doi/10.5406/amerjpsyc.134.3.033 3 | 30 | 5 | remote associations (correct > unsolved) | Verbal |
| Wu et al., 2021b | 1 | https://doi.org/10.3389/fpsyg.2021.672997 | 60 | 4 | Chinese compound remote association resolved > unresolved remote association | Verbal |
| Yu et al., 2021 | 1 | 10.1111/psyp.13886 | 46 | 6 | conjunction of insight problem solving: (high > low novelty) and (high > low appropriateness) | Verbal |
| Danek et al., 2019 | 1 | https://doi.org/10.3934/FNeuroscience.2019.2.60 | 32 | 42 | Magic trick problem solving: correct restructuring > incorrect restructuring | Nonverbal |
| Ren et al., 2020 | 1 | https://doi.org/10.1016/j.neuroimage.2020.116751 | 21 | 21 | novel and useful > familiar and useful | Nonverbal |
| Roberts et al., 2020 | 1 | https://doi.org/10.1016/j.neuroimage.2020.116758 | 27 | 9 | future thinking in response to cues: incongruent > congruent | Verbal |
| Zhang et al., 2020 | 1 | 10.1002/hipo.23253 | 24 | 16 | new combination of 2 objects: high + low creative > ordinary (early binding + late integration phase) | Nonverbal |
| Cai et al., 2018 | 1 | 10.1007/s11682-017-9689-8 | 16 | 11 | visual creative imagery task: mentally create unique and valuable products using 3 of the previously described items: imagery - control | Nonverbal |
| Mashal et al., 2013 | 1 | https://doi.org/10.1016/j.bandl.2012.11.012 | 14 | 2 | metaphor comprehension: novel > conventional metaphors | Verbal |
| Wu et al., 2016 | 1 | https://doi.org/10.1016/j.biopsycho.2016.03.011 | 20 | 17 | number problem solving - multiple solution > baseline + single solution < baseline | Nonverbal |
| Beaty et al., 2023 | 1 | https://doi.org/10.1037/aca0000603 | 51 | 14 | hypothesis generation and synonym generation | Verbal |
| Park et al., 2015 | 1 | https://doi.org/10.1016/j.neuropsychologia.2015.05.007 | 20 | 12 | Figural TTCT incomplete figures: Creative > control | Nonverbal |
| McPherson et al., | 1 | https://doi.org/10.1038/srep18460 | 20 | 5 | Music improvisation: positive > control | Nonverbal |

| **Reference** | **Contrast** | **DOI/Link** | **N** | **Foci** | **Task contrast** | **Input modality** |
|---|---|---|---|---|---|---|
| 2016 | | | | | | |
| McPherson et al., 2016 | 2 | https://doi.org/10.1038/srep18460 | 20 | 7 | Music improvisation: negative > control | Nonverbal |
| McPherson et al., 2016 | 3 | https://doi.org/10.1038/srep18460 | 20 | 2 | Music improvisation: ambiguous > control | Nonverbal |
