## Supplementary material for "The neural basis of creative thought: An activation likelihood estimation meta-analysis involving over 17,000 participants": Table S2

**Table S2: Contrasts included in the pre-existing knowledge dataset**

| Study | Contrast # | DOI | N | Foci | Contrast included | Input Modality | Subcomponent |
|---|---|---|---|---|---|---|---|
| Booth et al., 2006 | 1 | 10.1016/j.brainres.2005.11.097 | 13 | 3 | matched prior stimulus/not: Semantic decision > phonological (rhyme) | verbal | Semantic representation |
| Demonet et al., 1992 | 1 | 10.1093/brain/115.6.1753 | 9 | 9 | Semantic decision - word pair is positive attribute and small animal or not: Words > Phonemes | verbal | Semantic representation |
| Devlin et al., 2003 | 1 | 10.1162/089892903321107837 | 12 | 6 | Semantic decision > phonological decision | verbal | Semantic representation |
| Gesierich et al., 2012 | 1 | 10.1093/cercor/bhr286 | 21 | 40 | Same/different identity: Familiar faces > unfamiliar faces | nonverbal | Semantic representation |
| Gitelman et al., 2005 | 1 | 10.1016/j.neuroimage.2005.03.014 | 14 | 21 | semantic decision: semantic>control (include masked by phonological + orthographic processing) | verbal | Semantic representation |
| Gourovitch et al., 2000 | 1 | 10.1037/0894-4105.14.3.353 | 18 | 6 | Semantic generation > pho | verbal | Semantic representation |
| Kotz et al., 2002 | 1 | 10.1006/nimg.2002.1316 | 13 | 7 | lexical decision (auditory): Words > pseudowords | verbal | Semantic representation |
| Mechelli et al., 2007 | 1 | 10.1002/hbm.20272 | 20 | 8 | Reading aloud and naming: Semantically related > Phonologically related | verbal & nonverbal | Semantic representation |
| Mummery et al., 1998 | 1 | 10.1162/089892998563059 | 10 | 7 | Semantic decision > phonol | verbal | Semantic representation |
| Price et al., 1997 | 1 | 10.1162/jocn.1997.9.6.727. | 6 | 3 | Semantic decision > phonol | verbal | Semantic representation |
| Raposo et al., 2006 | 1 | 10.1016/j.neuropsychologia.2006.05.017 | 15 | 12 | semantic decision: unrelated > identity | verbal | Semantic representation |
| Roskies et al., 2001 | 1 | 10.1162/08989290152541485 | 20 | 22 | Semantic decision > phonological decision | verbal | Semantic representation |
| Seghier et al., 2010 | 1 | 10.1523/JNEUROSCI.3377-10.2010 | 94 | 4 | Semantics > perceptual decision | verbal & nonverbal | Semantic representation |
| Sugiura et al., 2006 | 1 | 10.1016/j.neuroimage.2006.01.002 | 24 | 17 | Familiarity judgments: personal > famous; famous > unfamiliar; personal > unfamiliar | verbal | Semantic representation |
| Wirth et al., 2011 | 1 | 10.1016/j.neuroimage.2010.10.039 | 21 | 7 | Living/non-living, syllable judgement: Semantic > phonological decision | verbal | Semantic representation |
| Gurd et al., 2002 | 1 | doi.org/10.1093/brain/awf093 | 11 | 8 | Category fluency > Rote Fluency | verbal | Semantic representation |
| Thioux et al., 2005 | 1 | 10.1016/j.cogbrainres.2005.02.009 | 6 | 2 | classification/comparison: animals>numbers (conjunction of 2 tasks) | verbal | Semantic representation |

| Study | Contrast # | DOI | N | Foci | Contrast included | Input Modality | Subcomponent |
|---|---|---|---|---|---|---|---|
| Xiao et al., 2005 | 1 | 10.1002/hbm.20105 | 14 | 3 | lexical decision: real >pseudowords | verbal | Semantic representation |
| Cappa et al., 1998 | 1 | 10.1006/nimg.1998.0368 | 13 | 6 | visual (size of tool/tail)/associative (kitchen tool/animal found in Italy) judgement: words>pseudowords | verbal | Semantic representation |
| Craik et al., 1999 | 1 | 10.1111/1467-9280.00102 | 8 | 12 | semantic judgement: how desirable/accurate the trait is>syllable | verbal | Semantic representation |
| Baumgaertner et al., 2007 | 1 | 10.1.1.906.1831 | 19 | 11 | acceptability judgement: action>non-action motion for sentences/videos | verbal | Semantic representation |
| Davis et al., 2004 | 1 | 10.1016/S0093-934X(03)00471-1 | 11 | 9 | One-back synonym monitoring: words> letter strings | verbal | Semantic representation |
| Devlin et al., 2002 | 1 | 10.1016/s0028-3932(01)00066-5 | 12 | 6 | any semantic category > letter detection | verbal | Semantic representation |
| Devlin et al., 2002 | 2 | 10.1016/s0028-3932(01)00066-5 | 8 | 33 | semantic categorisation (both animate + inanimate) > letter categorisation | verbal | Semantic representation |
| Ebisch et al., 2007 | 1 | 10.1093/cercor/bhm001 | 17 | 5 | semantic judgements - use two tools together for task, visuospatial - size of 1 vs other, control task - assess pseudoword/object: function + visuospatial>control (i.e. real objects/words>not) | verbal & nonverbal | Semantic representation |
| Giraud et al., 2001 | 1 | 10.1162/08989290152541421 | 12 | 3 | repeat/name object source of sound or 1/2 p's just say ok: words/meaningful sounds>noise /syllables | verbal & nonverbal | Semantic representation |
| Grossman et al., 2002a | 1 | 10.1006/nimg.2001.1028 | 16 | 3 | Pleasantness judgement - yes/no: all nouns>pseudowords | verbal | Semantic representation |
| Grossman et al., 2002b | 1 | 0.1002/hbm.10017 | 16 | 3 | pleasantness judgement (yes/no) or motion/cognition verbs: words>pseudowords | verbal | Semantic representation |
| Hagoort et al., 1999 | 1 | 10.1162/089892999563490 | 10 | 8 | words>pseudowords | verbal | Semantic representation |
| Henke et al., 1999 | 1 | 10.1162/089892999563490 | 12 | 7 | decide if two words inc. an unpleasant one or decide number of vowels across 2 words (</>6): semantic>orthographical | verbal | Semantic representation |
| Herbster et al., 1997 | 1 | 10.1002/(SICI)1097-0193 | 10 | 1 | Reading: regular > nonwords | verbal | Semantic representation |
| Binder et al., 1999 | 1 | 10.1162/089892999563265 | 30 | 4 | listen to a series of words, respond when heard animal in US and used by people = semantic, word with phonemes b and d =phono: semantic>phonology | verbal | Semantic representation |

| Study | Contrast # | DOI | N | Foci | Contrast included | Input Modality | Subcomponent |
|---|---|---|---|---|---|---|---|
| Joubert et al., 2004 | 1 | 10.1016/S0093-934X(03)00403-6 | 10 | 6 | reading: low frequency >nonwords | verbal | Semantic representation |
| Kuchinke et al., 2005 | 1 | 10.1016/j.neuroimage.2005.06.050 | 20 | 14 | Lexical decision: word>nonword | verbal | Semantic representation |
| Binder et al., 2003 | 1 | 10.1162/089892903321593108 | 24 | 35 | Lexical decision: word>nonword | verbal | Semantic representation |
| Noppeney et al., 2003b | 1 | 10.1016/S0093-934X(02)00525-4 | 9 | 7 | semantic decision (taste/colour/origin): normal>reversed words for all condition (taste, colour & origin) | verbal | Semantic representation |
| orfanidou et al., 2006 | 1 | 10.1162/jocn.2006.18.8.1237 | 13 | 33 | lexical decision: words>nonwords | verbal | Semantic representation |
| Perani et al., 1999 | 1 | 10.1016/S0028-3932(98)00073-6 | 11 | 6 | Same/different judgement: living/non-living objects > shape recognition | nonverbal | Semantic representation |
| Perani et al., 1999 | 2 | 10.1016/S0028-3932(98)00073-6 | 8 | 14 | Same/different judgement: living/non-living > nonwords | verbal | Semantic representation |
| Pilgrim et al., 2002 | 1 | 10.1006/nimg.2002.1105 | 14 | 16 | semantic categorisation: words > letter strings (orthographical cat) | verbal | Semantic representation |
| Rissman et al., 2003 | 1 | 10.1162/089892903322598120 | 15 | 5 | Lexical decision: words>nonwords | verbal | Semantic representation |
| Cai et al., 2007 | 1 | 10.1097/WNR.0b013e32810f2de7 | 15 | 2 | verbal repetition and syllable judgement: words>nonwords | verbal | Semantic representation |
| Damasio et al., 2001 | 1 | 10.1006/nimg.2001.077 | 10 | 44 | Naming: Actions (with/without implement) > control task | non-verbal | Semantic representation |
| Engelien et al., 2006 | 1 | 10.1007/s00702-005-0342-0 | 6 | 5 | Listening Meaningful sounds > meaningless sounds | non-verbal | Semantic representation |
| Rogers et al., 2006 | 1 | 10.3758/cabn.6.3.201 | 12 | 10 | Category decision: all semantic > baseline | non-verbal & verbal | Semantic representation |
| Tieleman et al., 2005 | 1 | 10.1016/j.neuroimage.2005.02.017 | 22 | 49 | self/fixed paced semantic categorisation (animal/object) task > perceptual (small/capital letter string) categorisation | verbal | Semantic representation |
| Bright et al., 2004 | 1 | 10.1016/j.bandl.2004.01.010 | 38 | 18 | lexical decision: conjunction of all semantic tasks > baseline | Verbal & nonverbal | Semantic representation |
| Bright et al., 2004 | 3 | 10.1016/j.bandl.2004.01.010 | 38 | 3 | Semantic decision: word only | verbal | Semantic representation |
| Bright et al., 2004 | 4 | 10.1016/j.bandl.2004.01.010 | 38 | 6 | semantic decision: picture only | non-verbal | Semantic representation |
| D'Arcy et al., 2007 | 1 | 10.1016/j.neures.2006.09.018 | 10 | 10 | Word to picture matching: (words + pictures) | nonverbal & | Semantic |

| Study | Contrast # | DOI | N | Foci | Contrast included | Input Modality | Subcomponent |
|---|---|---|---|---|---|---|---|
| | | | | | Semantic > baseline | verbal | representation |
| Devlin et al., 2000 | 1 | 10.1006/nimg.2000.059 | 16 | 6 | categorisation: semantic >baseline (PET) | verbal | Semantic representation |
| Devlin et al., 2000 | 2 | 10.1006/nimg.2000.059 | 16 | 7 | categorisation: semantic >baseline (fMRI) | verbal | Semantic representation |
| Gerlach et al., 1999 | 1 | 10.1093/brain/122.11.2159 | 15 | 18 | Object decision tasks > pattern discrimination | non-verbal | Semantic representation |
| Herbster et al., 1997 | 1 | 10.1002/(SICI)1097-0193(1997)5 | 10 | 6 | reading words: irregular and regular > say hiya | verbal | Semantic representation |
| Thierry et al., 2006 | 1 | 10.1162/jocn.2006.18.6.1018 | 12 | 5 | Semantic judgement (animal/ non animal) or ordered/not sequence: auditory sounds/words>scrambled | verbal + nonverbal | Semantic representation |
| Thierry et al., 2006 | 2 | 10.1162/jocn.2006.18.6.1018 | 12 | 5 | Semantic judgement (animal/ non animal) or ordered/not sequence: visual videos/words>distorted | verbal + nonverbal | Semantic representation |
| Tyler et al., 2003 | 1 | 10.1016/S1053-8119(02)00047-2 | 12 | 65 | Semantic judgement (related or unrelated) or orthographical judgement: tool/animal/tool action word/biological action > baseline | verbal | Semantic representation |
| Grabowski (1998 | 1 | 10.1006/nimg.1998.0324 | 9 | 3 | Naming animals/tools > say up/down | nonverbal | Semantic representation |
| Sergent et al., 1992 | 1 | doi.org/10.1093/brain/115.1.15 | 7 | 10 | semantic judgement: Identity > gender | non-verbal | Semantic representation |
| Sergent et al., 1992 | 2 | doi.org/10.1093/brain/115.1.15 | 7 | 8 | categorisation: Identity > gender | non-verbal | Semantic representation |
| Gorno-Tempini et al., 1998 | 1 | 10.1093/brain/121.11.2103 | 6 | 12 | match judgement: Famous Faces/names > controls; famous names > non-famous names | nonverbal + verbal | Semantic representation |
| Nakamura et al., 2000 | 1 | doi.org/10.1093/brain/123.9.1903 | 7 | 7 | face discrimination (familiar vs unfamiliar): Faces (fam vs unfamous task) > control | non-verbal | Semantic representation |
| Leveroni et al., 2000 | 1 | 10.1523/jneurosci.20-02-00878.2000 | 11 | 13 | pleasantness judgement (& remember for subsequent test): Familiar faces > unfamiliar | non-verbal | Semantic representation |
| sugiura et al., 2001 | 1 | 10.1006/nimg.2001.0747 | 5 | 19 | familiarity judgement: Identity discrimination (familiar or unfamiliar task) > control | non-verbal | Semantic representation |
| Grabowski et al., 2001 | 1 | 10.1002/hbm.1033.abs | 10 | 11 | naming people/landmarks>baseline (way up) tasks | non-verbal | Semantic representation |
| Damasio et al., 2004 | 1 | 10.1016/j.cognition.2002.07.001 | 68 | 25 | naming: people/animal/tools>scrambled | non-verbal | Semantic representation |
| Rothstein et al., 2005 | 1 | 10.1038/nn1370 | 20 | 4 | target face detection: Familiarity & identity change effect | non-verbal | Semantic representation |

| Study | Contrast # | DOI | N | Foci | Contrast included | Input Modality | Subcomponent |
|---|---|---|---|---|---|---|---|
| Elfgren et al., 2006 | 1 | 10.1016/j.neuroimage.2005.09.060 | 15 | 21 | familiarity judgement: Familiar > unfamiliar faces during identification | non-verbal | Semantic representation |
| Sugiura et al., 2008 | 1 | 10.1162/jocn.2008.21150 | 25 | 6 | familiarity detection: Familiar > unfamiliar | verbal | Semantic representation |
| Nielson et al., 2010 | 1 | 10.1016/j.bandc.2010.01.006 | 17 | 15 | familiarity judgement: Familiar > unfamiliar | verbal & nonverbal | Semantic representation |
| Brambati et al., 2010 | 1 | 10.1016/j.neuroimage.2010.06.045 | 12 | 20 | semantic categorisation: Specific judgement of occupation from faces | non-verbal | Semantic representation |
| Barense et al., 2011 | 1 | 10.1162/jocn_a_00010 | 18 | 8 | odd one out triad discrimination: Familiar faces > unfamiliar faces | non-verbal | Semantic representation |
| Gesierich et al., 2012 | 1 | 10.1093/cercor/bhr286 | 12 | 16 | judge if profession matches: Familiar > Unfamiliar | non-verbal | Semantic representation |
| Ross et al., 2012 | 1 | 10.1093/cercor/bhr274 | 16 | 10 | famousness judgement: Famous faces + landmarks > non famous | non-verbal | Semantic representation |
| Dreyer et al., 2017 | 1 | 10.1016/j.cortex.2017.10.021 | 28 | 12 | reading (silent) nouns>hashmarks | verbal | Semantic representation |
| Perrone-Bertolotti et al., 2017 | 1 | 10.3389/fnhum.2017.00325 | 24 | 7 | categorisation (living/non-living) words>unreadable font | verbal | Semantic representation |
| Liuzzi et al., 2017 | 1 | 10.1016/j.neuroimage.2017.02.032 | 18 | 14 | property verification>heard/saw stimuli? | verbal | Semantic representation |
| Redcay et al., 2016 | 1 | 10.1002/hbm.23251 | 24 | 7 | content matching judgement (e.g. include this word?): meaningful>meaningless | verbal & nonverbal | Semantic representation |
| Redcay et al., 2016 | 2 | 10.1002/hbm.23251 | 24 | 5 | content matching judgement (e.g. include this word?): communicative gesture>self adaptor gesture | nonverbal | Semantic representation |
| Sheldon et al., 2016 | 1 | 10.1016/j.neuropsychologia.2016.06.028 | 15 | 11 | semantic fluency>button press when see ‘X’ | verbal | Semantic representation |
| Haberling et al., 2016 | 1 | 10.1016/j.cortex.2016.06.003 | 91 | 33 | repetition judgement (i.e., seen before/not): pantomimes>(unknown) sign language/dog videos | nonverbal | Semantic representation |
| Haberling et al. (2016 | 2 | 10.1016/j.cortex.2016.06.003 | 91 | 16 | whether word pair is synonym/letter string matching: synonyms>letter strings | verbal | Semantic representation |
| Wang et al., 2016 | 1 | 10.3389/fpsyg.2016.00947 | 16 | 9 | lexical decision: WORDS>nonsense STROKES | verbal | Semantic representation |
| Kumar & Uttam (2016 | 1 | 10.1007/s10339-015-0738-1 | 20 | 8 | orthography judgment (include Matras/not): concrete and abstract words>nonwords | verbal | Semantic representation |
| Higuchi et al., 2015 | 1 | 10.1002/brb3.413 | 28 | 6 | text size judgement: real words > checkerboard | verbal | Semantic representation |

| Study | Contrast # | DOI | N | Foci | Contrast included | Input Modality | Subcomponent |
|---|---|---|---|---|---|---|---|
| Abdul Sabur et al., 2014 | 1 | 10.1016/j.cortex.2014.01.017 | 18 | 44 | Narrative processing > rhyme | verbal | Semantic representation |
| Slioussar et al., 2014 | 1 | 10.1016/j.bandl.2014.01.006 | 21 | 7 | word generation: real>non-verbs/non-nouns | verbal | Semantic representation |
| Bruffaerts et al., 2013 | 1 | 10.1523/JNEUROSCI.1548-13.2013 | 19 | 23 | property verification or pic/word for control task: pics + words > scrambled pics & consonants | Verbal & nonverbal | Semantic representation |
| Ludersdorfer et al., 2013 | 1 | 10.3389/fnhum.2013.00491 | 29 | 10 | one-back (visual): words>false fonts/pseudowords | verbal | Semantic representation |
| Ludersdorfer et al., 2013 | 2 | 10.3389/fnhum.2013.00491 | 29 | 2 | one-back (auditory): words>pseudowords/reversed speech | verbal | Semantic representation |
| Simard et al., 2013 | 1 | 10.1016/j.bandl.2011.08.002 | 14 | 41 | Wisconsin word sorting task: semantic>control/rhyme/syllable onset | verbal | Semantic representation |
| Straube et al., 2012 | 1 | 10.1371/journal.pone.0051207 | 16 | 13 | semantic judgement (incidental): known lang/iconic gesture>unknown lang/meaningless gesture | verbal + nonverbal | Semantic representation |
| Wende et al., 2012 | 1 | 10.1016/j.neuroimage.2012.06.003 | 18 | 10 | fluency: semantic fluency/association>phonological fluency | verbal | Semantic representation |
| Visser et al., 2012 | 1 | 10.1162/jocn_a_00244 | 15 | 4 | Camel & cactus/pyramid & palm trees: semantic>control | Verbal & nonverbal | Semantic representation |
| Visser et al., 2012 | 2 | 10.1162/jocn_a_00244 | 15 | 2 | Camel & cactus/pyramid & palm trees: picture>control | nonverbal | Semantic representation |
| Visser et al., 2012 | 3 | 10.1162/jocn_a_00244 | 15 | 7 | Camel & cactus/pyramid & palm trees: word>control | verbal | Semantic representation |
| Zhang et al., 2012 | 1 | 10.1016/j.ijpsycho.2012.02.013 | 14 | 6 | lexical decision; word>nonword | verbal | Semantic representation |
| Szlachta et al., 2012 | 1 | 10.1016/j.bandl.2012.02.007 | 21 | 40 | 1-back: words>musical rain, nouns, inflected nouns | verbal | Semantic representation |
| Carota et al., 2012 | 1 | 10.1162/jocn_a_00219 | 18 | 75 | silent reading: words (tools/animals/food) > hash mark strings | verbal | Semantic representation |
| Geranmayeh et al., 2012 | 1 | 10.1016/j.bandl.2012.02.005 | 19 | 10 | speech production: speech>tongue movements | verbal | Semantic representation |
| Welcome et al., 2012 | 1 | 10.1016/j.bandl.2011.12.011 | 20 | 7 | semantic judgement (animal or not): word meaning>word rhyme/word case | verbal | Semantic representation |
| Hervais-Adelman et al., 2012 | 1 | 10.1080/01690965.2012.662280 | 15 | 22 | target detection: clear/>noise vocoded speech (intelligibility) | verbal | Semantic representation |
| Hauk et al., 2011 | 1 | 10.3389/fnhum.2011.00149 | 21 | 10 | passive reading: reading>hashmarks | verbal | Semantic representation |

| Study | Contrast # | DOI | N | Foci | Contrast included | Input Modality | Subcomponent |
|---|---|---|---|---|---|---|---|
| Seghier et al., 2011 | 1 | 10.1093/cercor/bhq203 | 60 | 31 | word & picture pyramids & palm trees: semantic>perceptual matching on meaningless stimuli | verbal & nonverbal | Semantic representation |
| Hocking et al., 2011 | 1 | 10.1002/hbm.21040 | 13 | 8 | object attribute verification: environmental sounds > perceptual baseline | nonverbal | Semantic representation |
| Rapp et al., 2011 | 1 | 10.1162/jocn.2010.21507 | 10 | 10 | passive reading: words>checkerboards/consonant strings | verbal | Semantic representation |
| Khader et al., 201 | 1 | 10.1016/j.brainres.2009.12.082 | 16 | 19 | noun/verb generation > rhyme generation/letter search | verbal | Semantic representation |
| Kuchinke et al., 2009 | 1 | 10.1016/j.bandc.2008.07.014 | 15 | 7 | semantic judgement (taxonomic/sequential) > grammar judgement | verbal | Semantic representation |
| Cao et al., 2009 | 1 | 10.1002/hbm.20546 | 13 | 11 | relatedness judgement: >control (line judgement) | verbal | Semantic representation |
| Leff et al., 2008 | 1 | 10.1523/JNEUROSCI.2903-08.2008 | 26 | 3 | listen to speech judge gender: normal>reversed speech | verbal | Semantic representation |
| Alain et al., 2008 | 1 | 10.1162/jocn.2008.20014 | 16 | 8 | one-back: category>location judgement | verbal | Semantic representation |
| Sabri et al., 2008 | 1 | 10.1016/j.neuroimage.2007.09.052 | 28 | 5 | one-back: speech/word>rotated speech/pseudoword | verbal | Semantic representation |
| Vingerhoets (2008 | 1 | 10.1016/j.neuroimage.2007.12.058 | 14 | 4 | grasping orientation judgement: familiar>unfamiliar tools | nonverbal | Semantic representation |
| Garn et al., 2009 | 1 | 10.1016/j.cortex.2008.02.004 | 26 | 8 | picture naming: objects>scrambled | nonverbal | Semantic representation |
| Lin et al., 2015 | 1 | 10.1162/jocn_a_00852 | 20 | 3 | lexical decision: words>pseudowords | verbal | Semantic representation |
| Ludersdorfer et al., 2016 | 1 | 10.1016/j.neuroimage.2015.09.039 | 29 | 15 | semantic judgement (animacy): words>tones | verbal | Semantic representation |
| Segal & Petrides, 2012 | 1 | 10.1111/j.1460-9568.2011.07937.x | 90 | 14 | writing>copying | nonverbal | Semantic representation |
| Segal & Petrides, 2012 | 2 | 10.1111/j.1460-9568.2011.07937.x | 90 | 3 | copying: words>nonwords | verbal | Semantic representation |
| Carota et al., 2017 | 1 | 10.1093/cercor/bhw379 | 23 | 12 | detect typo: words>hashmarks | verbal | Semantic representation |
| Garbin et al., 2012 | 1 | 10.1371/journal.pone.0045091 | 12 | 12 | lexical decision: object or event noun/verb>pseudowords | verbal | Semantic representation |
| Vignali, 2019 | 1 | 10.1016/j.neuroimage.2018.08.061 | 21 | 22 | lexical decision (with flanker): words>pseudowords | verbal | Semantic representation |
| Wu et al., 2013 | 1 | 10.1038/srep02049 | 19 | 38 | passive reading: arm/leg/mouth | verbal | Semantic |

| Study | Contrast # | DOI | N | Foci | Contrast included | Input Modality | Subcomponent |
|---|---|---|---|---|---|---|---|
| | | | | | words>checkerboard baseline | | representation |
| Groussard et al., 2010 | 1 | 10.1016/j.neuroimage.2009.10.039 | 11 | 2 | proverb ending match judgement: verbal semantic>verbal reference | verbal | Semantic representation |
| Bozic et al., 2013 | 1 | 10.2298/PSI1304439B | 13 | 14 | gap detection: words>musical rain | verbal | Semantic representation |
| Bagga et al., 2013 | 1 | 10.1007/s12038-013-9387-7 | 18 | 5 | semantic judgement (concrete/abstract) or case judgement: semantic judgement>matching | verbal | Semantic representation |
| Chan et al., 2009 | 1 | 10.1016/j.neuroimage.2009.06.078 | 22 | 14 | passive reading: synonym/known language> pseudo characters/unknown language | verbal | Semantic representation |
| Malins et al., 2016 | 1 | 10.1016/j.neuropsychologia.2016.08.027 | 18 | 20 | passive reading: unrelated>false font/pseudoword | verbal | Semantic representation |
| Harrington et al., 2009 | 1 | 10.1016/j.cortex.2007.10.015 | 8 | 8 | drawing mental simulation: familiar>non-objects | nonverbal | Semantic representation |
| Taylor et al., 2009 | 1 | 10.1002/hbm.20646 | 10 | 24 | detect superimposition of all faces together: own/partner's/parent's face>unfamiliar | nonverbal | Semantic representation |
| Husain et al., 2012 | 1 | 10.1016/j.brainres.2012.08.029 | 16 | 22 | delayed match to sample of same gesture/same category gesture: meaningful iconic>meaningless gestures | nonverbal | Semantic representation |
| Lindenberg et al., 2012 | 1 | 10.1002/hbm.21258 | 20 | 8 | watch/imagine performing gesture: meaningful iconic>meaningless gestures | nonverbal | Semantic representation |
| Mashal et al., 2013 | 1 | 10.1016/j.bandl.2012.11.012 | 14 | 13 | covert relationship judgement: novel/real metaphor>unrelated words | verbal | Semantic representation |
| Christensen et al., 2008 | 1 | 10.1097/WNR.0b013e3283060a9d | 14 | 28 | gender and category judgement (e.g. female speaker, animal word): diotic/dichotic listening>reversed speech | verbal | Semantic representation |
| Emmorey et al., 2013 | 1 | 10.1016/j.bandl.2013.05.001 | 14 | 25 | concreteness judgement: semantic/phonological task words>false font | verbal | Semantic representation |
| Axmacher et al., 2009 | 1 | 10.1002/hbm.20645 | 32 | 3 | n-back: words>shapes | verbal | Semantic representation |
| Zaccarella et al., 2015 | 1 | 10.3389/fpsyg.2015.01818 | 22 | 3 | semantic judgement: words>pseudowords | verbal | Semantic representation |
| Jackson et al., 2015 | 1 | 10.1093/cercor/bhv003 | 24 | 18 | semantic relatedness: words>letter strings | verbal | Semantic representation |
| Zou et al., 2016 | 1 | 10.3339/fnhum.2015.00714 | 17 | 66 | morpheme matching after hearing speeches > tone | verbal | Semantic representation |
| Zhang et al., 2014 | 1 | 10.1016/j.cortex.2013.01.015 | 18 | 18 | nonbiological word relational task> asterisks | verbal | Semantic representation |
| Emmorey et al., 2010 | 1 | 10.1016/j.neuroimage.2009.08.001 | 14 | 8 | action observation: pantomimes>unknown | nonverbal | Semantic |

| Study | Contrast # | DOI | N | Foci | Contrast included | Input Modality | Subcomponent |
|---|---|---|---|---|---|---|---|
| | | | | | signs | | representation |
| Smith et al., 2012 | 1 | 10.1080/02643294.2012.706218 | 14 | 8 | property verification: semantic>control task | verbal | Semantic representation |
| Ryan et al., 2010 | 1 | 10.1002/hipo.20607 | 15 | 26 | semantic judgement task: spatial/nonspatial judgement > letter judgement/episodic judgement | verbal | Semantic representation |
| Roxbury et al., 2014 | 1 | 10.1186/1744-9081-10-34 | 17 | 7 | lexical decision: concrete/abstract word>pseudoword | verbal | Semantic representation |
| Hayashi et al., 2014 | 1 | 10.1016/j.neures.2013.10.007 | 16 | 22 | lexical decision: concrete/abstract words>asterisks | verbal | Semantic representation |
| Sachs et al., 2008 | 1 | 10.1016/j.neuropsychologia.2007.08.015 | 14 | 20 | categorisation: thematic/taxonomic relations>letters | verbal | Semantic representation |
| Obleser et al., 2008 | 1 | 10.1523/JNEUROSCI.1290-08.2008 | 16 | 4 | intelligibility rating: comprehension modulation in vocoded speech | verbal | Semantic representation |
| Jensen et al., 2011 | 1 | 10.1016/j.eplepsyres.2010.12.003 | 12 | 12 | lexical decision: words>nonwords | verbal | Semantic representation |
| Barros-Loscertales et al., 2012 | 1 | 10.1093/cercor/bhr324 | 59 | 11 | passive reading: words>hashmarks | verbal | Semantic representation |
| Raettig & Kotz, 2008 | 1 | 10.1016/j.neuroimage.2007.09.030 | 16 | 4 | lexical decision: words>pseudowords | verbal | Semantic representation |
| Moseley et al., 2012 | 1 | 10.1093/cercor/bhr238 | 18 | 32 | passive reading: emo words>hashmarks | verbal | Semantic representation |
| Weiss et al., 2015 | 1 | 10.1016/j.neuroimage.2015.07.029 | 18 | 30 | read aloud: pointed words>asterisks/say pass | verbal | Semantic representation |
| Chouinard et al., 2008 | 1 | 10.1016/j.neuroimage.2008.02.011 | 14 | 10 | naming intact objects > counting scrambled objects | nonverbal | Semantic representation |
| Zvyagintsev et al., 2013 | 1 | 10.1111/ejn.12140 | 15 | 7 | visual imagery of object >counting | nonverbal | Semantic representation |
| Bick et al., 2008 | 1 | 10.1162/jocn.2008.20.3.406 | 14 | 27 | semantic judgement (relatedness/morphological/orthographic/phonological): semantic task>pseudowords/visual control | verbal | Semantic representation |
| Raposo et al., 2016 | 1 | 10.1016/j.neuropsychologia.2016.06.036 | 18 | 8 | pleasantness judgement: semantic>perceptual task | verbal | Semantic representation |
| Heim et al., 2008 | 1 | 10.1016/j.neuroimage.2008.01.009 | 28 | 2 | semantic fluency >phonological fluency | verbal instruction prior to task | Semantic representation |
| Gutchess et al., 2010 | 1 | 10.1093/scan/nsp059 | 20 | 10 | semantic relatedness: relationships>same | verbal | Semantic |

| Study | Contrast # | DOI | N | Foci | Contrast included | Input Modality | Subcomponent |
|---|---|---|---|---|---|---|---|
| | | | | | words | | representation |
| Marques et al., 2008 | 1 | 10.1016/j.brainres.2007.11.070 | 21 | 8 | motion/visual feature judgement over press for plus signs: conjunction of visual and motion features >control task | verbal | Semantic representation |
| Menz et al., 2010 | 1 | 10.1016/j.neuroimage.2010.03.050 | 20 | 4 | familiarity judgement: known>unknown objects | nonverbal (except instructions) | Semantic representation |
| Ryan et al., 2008 | 1 | 10.1016/j.neuropsychologia.2008.02.030 | 10 | 16 | semantic fluency>button press when see “X” | verbal (cue only) | Semantic representation |
| Grindrod et al., 2014 | 1 | 10.1016/j.bandl.2014.10.001 | 23 | 11 | lexical decision: words>nonwords | verbal | Semantic representation |
| Chiao et al., 2009 | 1 | 10.1016/j.neuropsychologia.2008.09.023 | 12 | 32 | social status judgement: uniform/face/car status>colour change detection | nonverbal | Semantic representation |
| Jeon et al., 2009 | 1 | 10.1016/j.neuroimage.2009.06.049 | 16 | 8 | word generation/reading>nonword reading | verbal | Semantic representation |
| Sun et al., 2017 | 1 | 10.1016/j.jmr.2016.12.012 | 11 | 3 | semantic>orthography | verbal | Semantic representation |
| Liu et al., 2009 | 1 | 10.1162/jocn.2009.21141 | 16 | 24 | semantic judgement: words>slashes | verbal | Semantic representation |
| Liu et al., 2009 | 2 | 10.1162/jocn.2009.21141 | 16 | 20 | semantic judgement: words>tones | verbal | Semantic representation |
| Diaz & McCarthy. 2009 | 1 | 10.1016/j.brainres.2009.05.043 | 16 | 11 | match-to-sample nonwords: words>nonwords | verbal | Semantic representation |
| Taminato et al., 2014 | 1 | 10.1016/j.neures.2014.09.001 | 35 | 21 | object recognition >visual perception (before and after object is recognized) | nonverbal | Semantic representation |
| Wright et al., 2008 | 1 | 10.1002/hbm.20443 | 34 | 2 | n-back (1-back): semantic>visual matching | verbal & nonverbal | Semantic representation |
| Wright et al., 2008 | 2 | 10.1002/hbm.20443 | 34 | 2 | n-back (1-back): semantic>visual matching | verbal | Semantic representation |
| Wright et al., 2008 | 3 | 10.1002/hbm.20443 | 34 | 2 | n-back (1-back): semantic>visual matching | nonverbal | Semantic representation |
| Leung & Alain, 2010 | 1 | 10.1016/j.neuroimage.2010.12.055 | 16 | 6 | n-back (1/2-back): category>location matching | nonverbal | Semantic representation |
| Soch et al., 2017 | 1 | 10.1093/cercor/bhw206 | 110 | 9 | self/other attribute judgement>syllable counting | verbal | Semantic representation |
| Chou et al., 2009 | 1 | 10.1007/s00221-009-1942-y | 31 | 7 | relatedness judgement > character match judgement | verbal | Semantic representation |
| Wright et al., 2011 | 1 | 10.1162/jocn.2010.21450 | 14 | 6 | lexical decision: speech>musical rain | verbal | Semantic |

| Study | Contrast # | DOI | N | Foci | Contrast included | Input Modality | Subcomponent |
|---|---|---|---|---|---|---|---|
| | | | | | | | representation |
| Abraham, 2018 | 1 | 10.1016/j.neuropsychologia.2018.05.004 | 34 | 23 | Name uses of objects > name typical objects for location | verbal | Semantic control |
| Bitan et al., 2017 | 1 | 10.1037/neu0000357 | 23 | 4 | Subordinate>dominant meanings | verbal | Semantic control |
| Canini et al., 2016 | 1 | 10.1002/hbm.23304 | 24 | 2 | Interference time between exemplars of same category | nonverbal | Semantic control |
| Krieger-Redwood et al., 2015 | 1 | 10.1016/j.neuropsychologia.2015.02.030 | 22 | 35 | Weak>strong associations between words/pictures | verbal | Semantic control |
| Krieger-Redwood et al., 2015 | 2 | 10.1016/j.neuropsychologia.2015.02.030 | 22 | 18 | Strong>weak associations between words/pictures | verbal | Semantic control |
| Abraham et al., 2012 | 1 | 10.1016/j.neuropsychologia.2012.04.015 | 19 | 15 | High > low flexibility | verbal | Semantic control |
| Snyder et al., 2011 | 1 | 10.1162/jocn_a_00023 | 18 | 58 | High > low competition (with low/high association strength) | verbal | Semantic control |
| Balthasar et al., 2011 | 1 | 10.1016/j.brainres.2011.06.054 | 18 | 33 | Homonyms (auditory) > identification | verbal | Semantic control |
| Balthasar et al., 2011 | 2 | 10.1016/j.brainres.2011.06.054 | 18 | 26 | Homonyms (visual) > identification | verbal | Semantic control |
| Hsu et al., 2011 | 1 | 10.1162/jocn.2011.21619 | 12 | 13 | Low > high distance to foil (auditory) | verbal | Semantic control |
| Hsu et al., 2011 | 2 | 10.1162/jocn.2011.21619 | 12 | 13 | Low > high distance to foil (visual) | verbal | Semantic control |
| Rodd et al., 2010 | 1 | 10.1016/j.neuropsychologia.2009.12.035 | 14 | 1 | Semantic ambiguity > no ambiguity | verbal | Semantic control |
| Rodd et al., 2010 | 2 | 10.1016/j.neuropsychologia.2009.12.035 | 14 | 1 | Semantic ambiguity > syntactic ambiguity | verbal | Semantic control |
| Schnur et al., 2009 | 1 | 10.1073/pnas.0805874106 | 16 | 7 | More > less interference | nonverbal | Semantic control |
| Sun et al., 2016 | 1 | 10.1002/hbm.23246 | 28 | 1 | Alternative uses task > name typical objects for location | verbal | Semantic control |
| Grindrod et al., 2008 | 1 | 10.1016/j.brainres.2008.07.017 | 15 | 4 | Ambiguous incongruent meanings>single meaning | verbal | Semantic control |
| Jackson et al., 2015 | 1 | 10.1093/cercor/bhv003 | 24 | 8 | High>low association strength | verbal | Semantic control |
| Madore, 2019 | 1 | 10.1093/cercor/bhx312 | 32 | 59 | Alternate uses > associations after induction | verbal | Semantic control |
| Musz & Thompson- | 1 | 10.1016/j.bandl.2016.11.002 | 13 | 2 | Subordinate > dominant homonym meaning | verbal | Semantic |

| Study | Contrast # | DOI | N | Foci | Contrast included | Input Modality | Subcomponent |
|---|---|---|---|---|---|---|---|
| Schill, 2017 | | | | | | | control |
| Musz & Thompson-Schill, 2017 | 2 | 10.1016/j.bandl.2016.11.002 | 13 | | Delayed > immediate subordinate homonym meaning | verbal | Semantic control |
| Vitello et al., 2014 | 1 | 10.3389/fnhum.2014.00530 | 20 | 3 | Ambiguous word in sentence > not | verbal | Semantic control |
| Hallam et al., 2016 | 1 | 10.1016/j.neuropsychologia.2016.09.012 | 18 | 8 | Weak > strong association | verbal | Semantic control |
| Jeon, 2012 | 1 | 10.1097/WNR.0b013e32835a19ae | 14 | 4 | Ambiguous homonym > unambiguous word | verbal | Semantic control |
| Mestres-Misse et al., 2016 | 1 | 10.1007/s00429-014-0966-7 | 23 | 6 | Ambiguous homonym > unambiguous word | verbal | Semantic control |
| Hoenig & Scheef, 2009 | 1 | 10.1016/j.neuroimage.2008.12.044 | 22 | 15 | Ambiguous homonym > unambiguous word | verbal | Semantic control |
| Grindrod et al., 2014 | 1 | 10.1016/j.bandl.2014.10.001 | 23 | 10 | Ambiguous homonym > unambiguous word | verbal | Semantic control |
| Rodd et al., 2012 | 1 | 10.1093/cercor/bhr252 | 15 | 16 | Ambiguity across time | verbal | Semantic control |
| Tahmasebi et al., 2012 | 1 | 10.1093/cercor/bhr205 | 24 | 2 | Ambiguous homonym > unambiguous word | verbal | Semantic control |
| Hargreaves et al., 2011 | 1 | 10.1027/1618-3169/a000062 | 20 | 2 | Ambiguous > not | verbal | Semantic control |
| Bekinschtein et al., 2011 | 1 | 10.1523/JNEUROSCI.5058-10.2011 | 12 | 2 | Ambiguous homonym > unambiguous word | verbal | Semantic control |
| Satpute et al., 2014 | 1 | 10.1093/cercor/bhs408 | 33 | 6 | Weakly-related > strongly-related target | verbal | Semantic control |
| Tylen et al., 2015 | 1 | 10.1016/j.neuroimage.2015.07.047 | 24 | 12 | Incoherent > coherent story | verbal | Semantic control |
| Li et al., 2014 | 1 | 10.1093/scan/nst091 | 24 | 2 | Incongruent > congruent sentences | verbal | Semantic control |
| Mestres-Misse et al., 2014 | 1 | 10.1016/j.neuroimage.2014.05.002 | 23 | 1 | judge sentence congruency: subordinate > dominant homonym meaning | verbal | Semantic control |
| Mestres-Misse et al., 2014 | 2 | 10.1016/j.neuroimage.2014.05.002 | 23 | 15 | judge sentence congruency: Ambiguous > unambiguous word in sentences | verbal | Semantic control |
| Mestres-Misse et al., 2014 | 3 | 10.1016/j.neuroimage.2014.05.002 | 23 | 2 | judge sentence congruency: Incongruent>congruent sentence | verbal | Semantic control |
| Deen & McCarthy, 2010 | 1 | 10.1016/j.neuropsychologia.2010.01.028 | 15 | 3 | Incongruent>congruent story end | verbal | Semantic control |
| Smirnov et al., 2014 | 1 | 10.1016/j.neuropsychologia.2014.0 | 20 | 1 | Miscue > cue | both | Semantic |

| Study | Contrast # | DOI | N | Foci | Contrast included | Input Modality | Subcomponent |
|---|---|---|---|---|---|---|---|
| | | 9.007 | | | | | control |
| Newman et al., 2010 | 1 | 10.1016/j.bandl.2010.02.001 | 20 | 11 | Unrelated > related sentence | verbal | Semantic control |
| Mano et al., 2009 | 1 | 10.1016/j.neuropsychologia.2008.12.011 | 18 | 2 | Less > more coherent texts | verbal | Semantic control |
| Tune et al., 2016 | 1 | 10.1016/j.neuroimage.2016.05.020 | 18 | 21 | Easy/Hard anomalous > normal sentences | verbal | Semantic control |
| Peelle et al., 2009 | 1 | 10.1016/j.neuropsychologia.2008.10.027 | 25 | 15 | Inconsistent > consistent description for nominal kinds/nouns | verbal | Semantic control |
| Zhu et al., 2012 | 1 | 10.1016/j.neuroimage.2012.02.036 | 27 | 14 | Incongruence (cloze probability) in reading/semantic/font task | verbal | Semantic control |
| Willems et al., 2016 | 1 | 10.1093/cercor/bhv075 | 24 | 12 | Word context surprisal | verbal | Semantic control |
| Carter et al., 2019 | 1 | 10.1016/j.neuroimage.2019.01.018 | 41 | 10 | More > less surprisal of words in different contexts | verbal | Semantic control |
| Obleser & Kotz, 2010 | 1 | 10.1093/cercor/bhp128 | 16 | 6 | Low > high cloze probability | verbal | Semantic control |
| Scharinger et al., 2016 | 1 | 10.1002/hbm.23060 | 22 | 4 | Unpredictable > predictable word for incomplete > complete sentence | verbal | Semantic control |
| Huang et al., 2012 | 1 | 10.1016/j.brainres.2011.11.060 | 23 | 5 | Unexpected > expected word | verbal | Semantic control |
| Kambara et al., 2013 | 1 | 10.1016/j.langsci.2012.07.003 | 38 | 4 | Semantic violations > normal sentences | verbal | Semantic control |
| Rothermich & Kotz, 2013 | 1 | 10.1016/j.neuroimage.2012.12.013 | 16 | 5 | Unpredictable > predictable word when listening to sentence with abnormal/normal intonation | verbal | Semantic control |
| Nieuwland et al., 2012 | 1 | 10.1002/hbm.21377 | 24 | 9 | Semantic anomalies > normal sentences | verbal | Semantic control |
| Ye et al., 2014 | 1 | 10.1002/hbm.22182 | 20 | 3 | Incongruent > congruent sentences | verbal | Semantic control |
| van de Meerendonk et al., 2013 | 1 | 10.1016/j.bandl.2013.07.004 | 24 | 14 | Less congruent > more congruent words | verbal | Semantic control |
| van de Meerendonk et al., 2013 | 2 | 10.1016/j.bandl.2013.07.004 | 24 | 6 | Incongruent > congruent words | verbal | Semantic control |
| Willems, Ozyurek, Hagoort, 2008 | 1 | 10.1162/jocn.2008.20085 | 19 | 4 | Incongruent > congruent (words and sentences) | verbal | Semantic control |
| Willems et al., 2008 | 2 | 10.1162/jocn.2008.20085 | 19 | 1 | Incongruent > congruent (pictures nad sentences) | nonverbal | Semantic control |

| Study | Contrast # | DOI | N | Foci | Contrast included | Input Modality | Subcomponent |
|---|---|---|---|---|---|---|---|
| Raposo et al., 2012 | 1 | 10.1016/j.neuroimage.2011.08.072 | 17 | 1 | Rarer > prototypical features | verbal | Semantic control |
| Raposo et al., 2012 | 2 | 10.1016/j.neuroimage.2011.08.072 | 17 | 3 | Rarer > prototypical features of basic item | verbal | Semantic control |
| Kroeger et al., 2012 | 1 | 10.1016/j.brainres.2011.10.031 | 19 | 14 | Appropriate and unusual > inappropriate and usual uses | verbal | Semantic control |
| Rutter et al., 2012 | 1 | 10.1016/j.bandc.2011.11.002 | 18 | 7 | Appropriate and unusual > inappropriate and usual word choices | verbal | Semantic control |
| Zhu et al., 2013 | 1 | 10.1016/j.neuroimage.2013.02.060 | 26 | 11 | Violation > low cloze > high cloze parametric modulator in orthographic/semantic task | verbal | Semantic control |
| Sitnikova et al., 2014 | 1 | 10.1016/j.neuroimage.2014.09.012 | 16 | 3 | Novel > typical object use | nonverbal | Semantic control |
| Moberget et al., 2014 | 1 | 10.1523/JNEUROSCI.2264-13.2014 | 32 | 6 | Incongruent > congruent sentences | verbal | Semantic control |
| Clos et al., 2014 | 1 | 10.1002/hbm.22151 | 29 | 1 | Cue different > same sentence | verbal | Semantic control |
| Van Ettinger-Veenstra et al., 2016 | 1 | 10.3389/fnhum.2016.00110 | 27 | 2 | Incongruent > congruent sentences | verbal | Semantic control |
| Tobia et al., 2017 | 1 | 10.14814/phy2.13078 | 16 | 4 | Subordinate > prototypical uses | nonverbal | Semantic control |
| Ferstl et al, 2005 | 1 | 10.1162/0898929053747658 | 20 | 1 | Incongruent > congruent | verbal | Semantic control |
| Allen et al., | 1 | 10.1162/jocn.2008.20107 | 15 | 5 | Inappropriate > appropriate | verbal | Semantic control |
| Badre et al., 2005 | 1 | 10.1016/j.neuron.2005.07.023 | 22 | 19 | Weak > strong association | verbal | Semantic control |
| Badre et al., 2005 | 2 | 10.1016/j.neuron.2005.07.023 | 22 | 10 | Incongruent > congruent relations | verbal | Semantic control |
| Bedny et al, 2008 | 1 | 10.1093/cercor/bhn018 | 20 | 11 | Ambiguous > not ambiguous | verbal | Semantic control |
| Bunge, 2005 | 1 | https://academic.oup.com/cercor/article/15/3/239/375113 | 20 | 1 | Low > high relation between probe and target | verbal | Semantic control |
| Chan et al., 2004 | 1 | 10.1016/j.neuroimage.2004.02.034 | 8 | 17 | Semantic ambiguity > non-ambiguous words | verbal | Semantic control |
| de Zubicaray et al., 2000 | 1 | 10.1016/S0028-3932(00)00026-9 | 8 | 14 | Inappropriate > appropriate responses | verbal | Semantic control |
| Gennari et al, 2007 | 1 | 10.1016/j.neuroimage.2007.01.015 | 17 | 4 | Ambiguous > unambiguous | verbal | Semantic control |

| Study | Contrast # | DOI | N | Foci | Contrast included | Input Modality | Subcomponent |
|---|---|---|---|---|---|---|---|
| Gurd et al., 2002 | 1 | 10.1093/brain/awf093 | 11 | 2 | Switching > not switching | verbal | Semantic control |
| Hirshorn & Thompson Schill, 2006 | 1 | 10.1016/j.neuropsychologia.2006.03.035 | 10 | 15 | Switching > free generation | verbal | Semantic control |
| Hirshorn & Thompson Schill, 2006 | 2 | 10.1016/j.neuropsychologia.2006.03.035 | 10 | 21 | Switching > clustering | verbal | Semantic control |
| Ketteler et al., 2008 | 1 | 10.1016/j.neuroimage.2007.10.023 | 12 | 16 | Weak > strong target relation and strong > weak distractor relation | verbal | Semantic control |
| Nagel et al, 2008 | 1 | 10.1016/j.neuroimage.2008.07.017 | 14 | 3 | High > low selection | verbal | Semantic control |
| Nelson et al, 2009 | 1 | 10.1016/j.brainres.2008.12.001 | 17 | 5 | Many > few associates | verbal | Semantic control |
| Noppeney et al., 2004 | 2 | 10.1016/j.neuroimage.2003.12.010 | 15 | 9 | Difficult > easy judgements | verbal | Semantic control |
| Persson et al., 2004 | 1 | 10.1016/j.neuroimage.2004.08.004 | 22 | 4 | High > low selection | verbal | Semantic control |
| Race et al., 2009 | 1 | 10.1162/jocn.2009.21132 | 26 | 17 | DIFFERENT>SAME ATTRIBUTE/reversed/novel decision > repeated | verbal | Semantic control |
| Roskies et al., 2001 | 1 | 10.1162/08989290152541485 | 20 | 2 | Low > high category typicality | verbal | Semantic control |
| Snyder et al., 2007 | 1 | 10.1162/jocn.2007.19.5.761 | 14 | 13 | Specific attribute related > globally semantically related | verbal | Semantic control |
| Spalek et al., 2008 | 1 | 10.1016/j.bandl.2008.05.005 | 21 | 2 | Semantically related > unrelated distractors | both | Semantic control |
| Thompson schill et al., 1997 | 1 | 10.1073/pnas.94.26.14792 | 6 | 8 | High > low selection (pictures) | nonverbal | Semantic control |
| Thompson schill et al., 1997 | 2 | 10.1073/pnas.94.26.14792 | 6 | 8 | High > low selection (words) | verbal | Semantic control |
| Thompson Schill et al., 1999 | 4 | 10.1016/S0896-6273(00)80804-1 | 8 | 1 | Different > same attribute | verbal | Semantic control |
| Wagner et al., 2001 | 1 | 10.1016/S0896-6273(01)00359-2 | 14 | 12 | Weak > strong association | verbal | Semantic control |
| Wagner et al., 2001 | 2 | 10.1016/S0896-6273(01)00359-2 | 14 | 6 | More > less foils | verbal | Semantic control |
| Whitney et al., 2009 | 1 | 10.1093/cercor/bhp007 | 18 | 7 | Subordinate > dominant word ambiguity | verbal | Semantic control |
| Wig et al., 2009 | 1 | 10.1152/jn.91213.2008 | 27 | | Reversed > different decision | nonverbal | Semantic |

| Study | Contrast # | DOI | N | Foci | Contrast included | Input Modality | Subcomponent |
|---|---|---|---|---|---|---|---|
| | | | | | | | control |
| Zhang et al., 2004 | 1 | 10.1016/j.neuroimage.2004.07.008 | 14 | 7 | High conflict > low conflict/neutral words | verbal | Semantic control |
| Collette et al., 2001 | 1 | 10.1006/nimg.2001.0846 | 12 | 22 | Inhibit>initiate | verbal | Semantic control |
| Mason & Just, 2007 | 1 | 10.1016/j.brainres.2007.02.076 | 12 | 21 | Ambiguous > unambiguous | verbal | Semantic control |
| Zempleni et al., 2007 | 1 | 10.1016/j.neuroimage.2006.09.048 | 16 | 5 | Ambiguous > unambiguous | verbal | Semantic control |
| Liu et al., 2009 | 1 | 10.1162/jocn.2009.21141 | 16 | 2 | Weak > strong association | verbal | Semantic control |
| Liu et al., 2009 | 2 | 10.1162/jocn.2009.21141 | 16 | 2 | Weak > strong association | verbal | Semantic control |
| Addis et al., 2007 | 1 | https://doi.org/10.1016/j.neuropsychologia.2006.10.016 | 14 | 23 | event elaboration (autobiographical memory covert elaboration of past/future event with written cue) > retrieve two words related to the cued word then generate a sentence/think of 2 objects related to the stimulus and visualize all 3 objects in a triangular arrangement | Verbal | Recollections |
| Addis et al., 2011 | 1 | https://doi.org/10.1002/hipo.20870 | 15 | 9 | conjunction of autobiographical tasks (covert generation of autobiographical memory based on written cue)> imagery/semantic decision tasks | Verbal | Recollections |
| Addis et al., 2012 | 1 | https://doi.org/10.1016/j.neuroimage.2011.09.066 | 15 | 17 | event elaboration (autobiographical memory covert elaboration of past/future event with written cue) > retrieve two words related to the cued word then generate a sentence/think of 2 objects related to the stimulus and visualize all 3 objects in a triangular arrangement | Verbal | Recollections |
| Audrain et al., 2022 | 1 | https://doi.org/10.1523/JNEUROSCI.0832-22.2022 | 40 | 25 | overtly generate the recalled autobiographical event after choosing from 2 pictures > overtly describe the picture | Nonverbal | Recollections |
| Compere et al., 2016 | 1 | https://doi.org/10.3389/fnhum.2016.00285 | 20 | 4 | men: covertly recall autobiographical event > sentence completion and visualize the sentence they completed | Verbal | Recollections |
| Compere et al., 2016 | 2 | https://doi.org/10.3389/fnhum.2016.00285 | 18 | 11 | women: covertly recall autobiographical event > sentence completion and visualize the | Verbal | Recollections |

| Study | Contrast # | DOI | N | Foci | Contrast included | Input Modality | Subcomponent |
|---|---|---|---|---|---|---|---|
| | | | | | sentence they completed | | |
| D'Argembeau et al., 2014 | 1 | https://doi.org/10.1093/scan/nst028 | 24 | 13 | EAM remembering > EAM reasoning | Verbal | Recollections |
| D'Argembeau et al., 2014 | 2 | https://doi.org/10.1093/scan/nst028 | 24 | 7 | EAM reasoning > EAM remembering | Verbal | Recollections |
| Daselaar et al., 2008 | 1 | https://doi.org/10.1093/cercor/bhm048 | 17 | 9 | Access > elaboration | Verbal | Recollections |
| Daselaar et al., 2008 | 2 | https://doi.org/10.1093/cercor/bhm048 | 17 | 10 | Elaboration > access | Verbal | Recollections |
| Denkova et al., 2015 | 1 | https://doi.org/10.1093/scan/nsu039 | 18 | 26 | overtly generating autobiographical response (written cue) > semantic verbal fluency | Verbal | Recollections |
| Ford et al., 2011 | 1 | https://doi.org/10.1016/j.neuropsychologia.2011.04.032 | 18 | 21 | autobiographical memory search upon musical cue > select an adjective that describe the music and give a definition to the adjective selected | Verbal | Recollections |
| Ford et al., 2016 | 1 | https://doi.org/10.1037/pmu0000152 | 16 | 19 | All EAM > control search (young adults) | Verbal | Recollections |
| Gardini et al., 2006 | 1 | https://doi.org/10.1016/j.neuroimage.2005.10.012 | 14 | 20 | EAM >pseudoword reading (silent) | Verbal | Recollections |
| Gilmore et al., 2021 | 1 | https://doi.org/10.1523/JNEUROSCI.1490-20.2020 | 46 | 16 | autobiographical recall > picture description | Nonverbal | Recollections |
| Gurguryan & Sheldon, 2019 | 1 | https://doi.org/10.1016/j.neuroimage.2019.05.077 | 28 | 13 | retrieval orientation (conceptual and contextual) > number detection task | Verbal | Recollections |
| Gurguryan & Sheldon, 2019 | 2 | https://doi.org/10.1016/j.neuroimage.2019.05.077 | 28 | 12 | stage of retrieval (re-oriented and oriented) > number detection task | Verbal | Recollections |
| Holland et al., 2011 | 1 | https://doi.org/10.1016/j.neuropsychologia.2011.07.015 | 31 | 9 | covertly generate autobiographical memory from written cue > sentence generation with given format (x is smaller than y is smaller than z) | Verbal | Recollections |
| Ino et al., 2011 | 1 | https://doi.org/10.2174/1874440001105010014 | 21 | 13 | overtly generate autobiographical memory > rest and autobiographical memory > semantic memory | Verbal | Recollections |
| Iriye & St Jacques, 2020 | 1 | https://doi.org/10.1016/j.cortex.2020.05.007 | 25 | 16 | negative EAM > baseline (time lag 3) | Nonverbal | Recollections |
| Iriye & St Jacques, 2020 | 2 | https://doi.org/10.1016/j.cortex.2020.05.007 | 25 | 21 | negative EAM > baseline (time lag 4) | Nonverbal | Recollections |
| Iriye & St Jacques, 2020 | 3 | https://doi.org/10.1016/j.cortex.2020.05.007 | 25 | 31 | negative EAM > baseline (time lag 5) | Nonverbal | Recollections |

| Study | Contrast # | DOI | N | Foci | Contrast included | Input Modality | Subcomponent |
|---|---|---|---|---|---|---|---|
| Iriye & St Jacques, 2020 | 4 | https://doi.org/10.1016/j.cortex.2020.05.007 | 25 | 23 | negative EAM > baseline (time lag 6) | Nonverbal | Recollections |
| Martinelli et al., 2013 | 1 | https://doi.org/10.1371/journal.pone.0082385 | 20 | 9 | young: overt autobiographical memory retrieval > overt sentence completion | Verbal | Recollections |
| McCormick et al., 2015 | 1 | https://doi.org/10.1093/cercor/bht324 | 18 | 5 | EAM > math task (lag 1) | Verbal | Recollections |
| McCormick et al., 2015 | 2 | https://doi.org/10.1093/cercor/bht324 | 18 | 13 | EAM > math task (lag 2) | Verbal | Recollections |
| McCormick et al., 2015 | 3 | https://doi.org/10.1093/cercor/bht324 | 18 | 7 | EAM > math task (lag 3) | Verbal | Recollections |
| McCormick et al., 2015 | 4 | https://doi.org/10.1093/cercor/bht324 | 18 | 11 | EAM > math task (lag 4) | Verbal | Recollections |
| McCormick et al., 2015 | 5 | https://doi.org/10.1093/cercor/bht324 | 18 | 11 | EAM > math task (lag 5) | Verbal | Recollections |
| McCormick et al., 2015 | 6 | https://doi.org/10.1093/cercor/bht324 | 18 | 13 | EAM > math task (lag 6) | Verbal | Recollections |
| McCormick et al., 2015 | 7 | https://doi.org/10.1093/cercor/bht324 | 18 | 11 | EAM > math task (lag 7) | Verbal | Recollections |
| Monge et al., 2018 | 1 | https://doi.org/10.1016/j.neuropsychologia.2017.07.030 | 18 | 7 | autobiographical memory > laboratory memory recovery | Nonverbal | Recollections |
| Rabin et al., 2009 | 1 | https://doi.org/10.1162/jocn.2009.21344 | 18 | 10 | EAM > TOM construction | Nonverbal | Recollections |
| Rabin et al., 2009 | 2 | https://doi.org/10.1162/jocn.2009.21344 | 18 | 9 | EAM > TOM elaboration | Nonverbal | Recollections |
| Rabin & Rosenbaum, 2012 | 1 | https://doi.org/10.1016/j.neuroimage.2012.05.002 | 18 | 4 | EAM > pTOM (TR 1, 2) | Nonverbal | Recollections |
| Rabin & Rosenbaum, 2012 | 2 | https://doi.org/10.1016/j.neuroimage.2012.05.002 | 18 | 1 | EAM > pTOM (TR 4, 5) | Nonverbal | Recollections |
| Sheldon & Moscovitch, 2011 | 1 | https://doi.org/10.1002/hipo.20985 | 20 | 7 | overtly generate words after reading written autobiographical cue > visuomotor baseline task | Verbal | Recollections |
| Speer et al., 2014 | 1 | https://doi.org/10.1016/j.neuron.2014.09.028 | 19 | 27 | positive > neutral EAM | Verbal | Recollections |
| Speer et al., 2014 | 2 | https://doi.org/10.1016/j.neuron.2014.09.028 | 19 | 1 | neutral > positive EAM | Verbal | Recollections |
| Sperduti et al., 2013 | 1 | https://doi.org/10.3389/fnbeh.2013.00041 | 20 | 35 | covertly generating descriptions that describe the autobiographical memory after written cue > rest | Verbal | Recollections |

| Study | Contrast # | DOI | N | Foci | Contrast included | Input Modality | Subcomponent |
|---|---|---|---|---|---|---|---|
| St Jacques et al., 2012 | 1 | https://doi.org/10.1016/j.neurobiologing.2010.11.007 | 17 | 35 | activation during overt generation of autobiographical memory > rest | Verbal | Recollections |
| Summerfield et al., 2009 | 1 | https://doi.org/10.1016/j.neuroimage.2008.09.033 | 18 | 18 | EAM (real + imagined) > films and new events (real + imagined) | Verbal | Recollections |
| Addis et al., 2004a | 1 | https://doi.org/10.1016/j.neuroimage.2004.08.007 | 14 | 4 | specific EAM > general EAM (lag 2) | Verbal | Recollections |
| Addis et al., 2004a | 2 | https://doi.org/10.1016/j.neuroimage.2004.08.007 | 14 | 16 | specific EAM > general EAM (lag 3) | Verbal | Recollections |
| Addis et al., 2004b | 1 | https://doi.org/10.1002/hipo.10215 | 14 | 16 | specific and general AM retrieval > sentence completion and size discrimination | Verbal | Recollections |
| Arshamian et al., 2013 | 1 | https://doi.org/10.1016/j.neuropsychologia.2012.10.023 | 15 | 4 | odour cued EAM > control odours | Nonverbal | Recollections |
| Arshamian et al., 2013 | 2 | https://doi.org/10.1016/j.neuropsychologia.2012.10.023 | 15 | 12 | word cued EAM > control words | Verbal | Recollections |
| Botzung et al., 2008a | 1 | https://doi.org/10.1080/09658210801931222 | 10 | 34 | EAM > semantic memory | Verbal | Recollections |
| Botzung et al., 2008b | 1 | https://doi.org/10.1016/j.bandc.2007.07.011 | 10 | 11 | past events evocation main effect | Verbal | Recollections |
| Cabeza et al., 2004 | 1 | https://doi.org/10.1162/0898929042568578 | 13 | 10 | Conjunction: EAM triggered by photos taken by self and EAM triggered by photos taken by others | Nonverbal | Recollections |
| Chen et al., 2017 | 1 | https://doi.org/10.1523/JNEUROSCI.1534-16.2017 | 31 | 30 | EAM: life memory > picture memory | Nonverbal | Recollections |
| Denkova et al., 2006 | 1 | https://doi.org/10.1016/j.brainres.2006.01.061 | 10 | 26 | EAM (relatives and friends' faces)> control (face identification; yes for familiar faces and no for unknown faces) | Nonverbal | Recollections |
| Detour et al., 2011 | 1 | https://doi.org/10.1016/j.brainres.2011.05.024 | 8 | 9 | presented personal photographs mixed with photographs of unknown people: discrimination (remember/know/new) trial > observation trial | Nonverbal | Recollections |
| Donix et al., 2010 | 1 | https://doi.org/10.1093/arclin/acq037 | 15 | 17 | EAM > semantic memory (younger subjects) | Verbal | Recollections |
| Eich et al., 2009 | 1 | https://doi.org/10.1016/j.neuropsychologia.2009.02.019 | 16 | 22 | first person perspective memory retrieval (field perspective) main effect | Nonverbal | Recollections |
| Fleischer et al., 2019 | 1 | https://doi.org/10.1016/j.bbr.2018.06.024 | 33 | 8 | EAM > arithmetic (placebo condition) | Verbal | Recollections |
| Gilboa et al., 2004 | 1 | https://doi.org/10.1093/cercor/bhh082 | 9 | 18 | main effect of vividly reexperienced events | Nonverbal | Recollections |

| Study | Contrast # | DOI | N | Foci | Contrast included | Input Modality | Subcomponent |
|---|---|---|---|---|---|---|---|
| Grol et al., 2017 | 1 | https://doi.org/10.1016/j.bandc.2016.09.014 | 27 | 21 | imagery of memory from field perspective > visual search (numbers) | Verbal | Recollections |
| Hoscheidt et al., 2010 | 1 | https://doi.org/10.1016/j.bbr.2010.04.010 | 17 | 36 | EAM spatial > read ungrammatical sentences | Verbal | Recollections |
| Hoscheidt et al., 2010 | 2 | https://doi.org/10.1016/j.bbr.2010.04.010 | 17 | 35 | EAM nonspatial > read ungrammatical sentences | Verbal | Recollections |
| Lux et al., 2013 | 1 | https://doi.org/10.1080/13554794.2013.860174 | 13 | 5 | EAM with personalized sentence cues: main effect of temporal and locational context | Verbal | Recollections |
| Maguire and Frith, 2003 | 1 | https://doi.org/10.1093/brain/awg157 | 12 | 11 | EAM > word recognition control task | Verbal | Recollections |
| Muscatell et al., 2010 | 1 | https://doi.org/10.1093/scan/nsp043 | 13 | 13 | EAM > sentence reading | Verbal | Recollections |
| Niki and Luo, 2002 | 1 | https://doi.org/10.1162/089892902317362010 | 9 | 20 | recall the paces where participants visited 7 years ago or within 2 years when they were presented with landmarks of the places (remote > recent memories) | Verbal | Recollections |
| Noreen et al., 2016 | 1 | https://doi.org/10.3389/fpsyg.2016.00379 | 22 | 24 | EAM > EAM suppression (major and minor clusters) | Verbal | Recollections |
| Oddo et al., 2010 | 1 | https://doi.org/10.1016/j.cortex.2008.07.003 | 15 | 6 | EAM > semantic memory | Verbal | Recollections |
| Rekkas and Constable, 2005 | 1 | https://doi.org/10.1162/089892905775008652 | 12 | 24 | retrieval of remote EAM main effect | Verbal | Recollections |
| St Jacques et al., 2013 | 1 | https://doi.org/10.1073/pnas.1319630110 | 35 | 3 | recognition memory: subsequent true memory for novel encoding | Nonverbal | Recollections |
| St Jacques et al., 2018 | 1 | https://doi.org/10.1016/j.neuropsychologia.2017.06.015 | 29 | 7 | EAM > counterfactual simulation | Verbal | Recollections |
| Svoboda and Levine, 2009 | 1 | https://doi.org/10.1523/JNEUROSCI.3452-08.2009 | 11 | 27 | main effect: EAM rehearsal (1st rehearsal) | Verbal | Recollections |
| Young et al., 2013 | 1 | https://doi.org/10.1002/hbm.22144 | 40 | 18 | specific AM recall > example generation (male + female) | Verbal | Recollections |
| Achim & Lepage, 2005a | 1 | https://doi.org/10.1162/0898929053467578 | 18 | 3 | Associative recognition > item recognition | Nonverbal | Paired associates |
| Achim & Lepage, 2005b | 1 | https://doi.org/10.1016/j.neuroimage.2004.10.036 | 18 | 3 | associative recognition: intact > rearranged | Nonverbal | Paired associates |
| Achim & Lepage, 2005b | 2 | https://doi.org/10.1016/j.neuroimage.2004.10.036 | 18 | 9 | associative recognition: rearranged > intact | Nonverbal | Paired associates |
| Bader et al., 2014 | 1 | https://doi.org/10.1016/j.neuropsychologia.2014.06.006 | 20 | 4 | PAL: general recognition (same > new) | Verbal | Paired associates |
| Bisby et al., 2016 | 1 | https://doi.org/10.1093/scan/nsw02 | 20 | 8 | main effect of associative memory (faces) | Nonverbal | Paired |

| Study | Contrast # | DOI | N | Foci | Contrast included | Input Modality | Subcomponent |
|---|---|---|---|---|---|---|---|
| | | 8 | | | | | associates |
| Burgess et al., 2001 | 1 | https://doi.org/10.1006/nimg.2001.0806 | 13 | 13 | person-object association main effect (Table 4) | Nonverbal | Paired associates |
| Burgess et al., 2001 | 2 | https://doi.org/10.1006/nimg.2001.0806 | 13 | 18 | person-place association main effect (Table 6) | Nonverbal | Paired associates |
| Cabeza et al., 1997 | 1 | https://doi.org/10.1162/jocn.1997.9.2.254 | 12 | 6 | PAL recognition > reading | Verbal | Paired associates |
| Dennis et al., 2014 | 1 | https://doi.org/10.1016/j.bandc.2014.04.009 | 18 | 8 | PAL: recognition (correct recognition > correct rejection) | Verbal | Paired associates |
| Ford et al., 2010 | 1 | https://doi.org/10.1016/j.neuropsychologia.2010.06.010 | 18 | 7 | word pair: unrelated word and compound word (intact > recombined) | Verbal | Paired associates |
| Giovanello et al., 2004 | 1 | https://doi.org/10.1002/hipo.10182 | 16 | 22 | word pair associative > item recognition | Verbal | Paired associates |
| Hannula et al., 2009 | 1 | https://doi.org/10.1016/j.neuron.2009.08.025 | 14 | 11 | scene-face pair: correct > incorrect recognition (Table S1) | Nonverbal | Paired associates |
| Holdstock et al., 2010 | 1 | https://doi.org/10.1016/j.neuropsychologia.2010.08.018 | 12 | 43 | face-laugh/face-face recognition > perceptual baseline | Nonverbal | Paired associates |
| Huijbers et al., 2013 | 1 | https://doi.org/10.1162/jocn_a_00366 | 45 | 13 | face-name pairs: hit (categorize intact pairs as old ones)> missed (categorized intact pairs as rearranged/new one) recognition | Verbal and nonverbal | Paired associates |
| King et al., 2005 | 1 | https://doi.org/10.1016/j.neuroimage.2005.05.057 | 13 | 19 | person-object association main effect | Nonverbal | Paired associates |
| Lepage et al., 2003 | 1 | https://doi.org/10.1016/S0304-3940(03)00578-0 | 6 | 10 | line-drawing pairs: intact +rearranged pairs > new pairs | Nonverbal | Paired associates |
| Li et al., 2016 | 1 | https://doi.org/10.1155/2016/9860604 | 16 | 12 | object pair: contextual > noncontextual | Nonverbal | Paired associates |
| Li et al., 2016 | 2 | https://doi.org/10.1155/2016/9860604 | 16 | 17 | shape pair: contextual > noncontextual | Nonverbal | Paired associates |
| Okada et al., 2011 | 1 | https://doi.org/10.1371/journal.pone.0024862 | 15 | 14 | adjective + face pair: recognition > word | Nonverbal | Paired associates |
| Onoda et al., 2009 | 1 | https://doi.org/10.1016/j.neures.2009.01.008 | 25 | 7 | word pair (neutral) > baseline | Verbal | Paired associates |
| Onoda et al., 2009 | 2 | https://doi.org/10.1016/j.neures.2009.01.008 | 25 | 7 | word pair (negative) > baseline | Verbal | Paired associates |
| Park et al., 2014 | 1 | https://doi.org/10.1016/j.neulet.2014.08.024 | 19 | 12 | object pair: associative recognition (hit > miss) | Nonverbal | Paired associates |
| Sherman et al., 2015 | 1 | https://doi.org/10.1016/j.neuropsychologia.2015.07.020 | 41 | 7 | successful associative memory retrieval (correct memory for intact vs control task) | Verbal | Paired associates |
