## Supplementary material for "The neural basis of creative thought: An activation likelihood estimation meta-analysis involving over 17,000 participants": Table S3

**Table S3: Contrasts included in the executive mechanisms dataset**

| Reference and year | Contrast # | DOI | N | Foci | Task name, brief description and contrast | Input modality | Subcomponent |
|---|---|---|---|---|---|---|---|
| Warburton et al., 1996 | 1 | 10.1093/brain/119.1.159 | 9 | 14 | generation of verb given a noun: generation > rest | Verbal | Energization |
| Hwang et al., 2009 | 1 | 10.1016/j.neuroimage.2009.06.042 | 13 | 6 | semantic verbal fluency>word repetition | Verbal | Energization |
| Birn et al., 2011 | 1 | 10.1016/j.neuroimage.2009.07.036 | 14 | 10 | fluency (category) >recite months | verbal | Energization |
| Gurd et al (2002) | 1 | doi.org/10.1093/brain/awf093 | 11 | 8 | Category fluency > recite months | verbal | Energization |
| Badzakova-Trajkov et al., 2011 | 1 | https://doi.org/10.1016/j.neuropsychologia.2011.06.016 | 139 | 10 | covert word generation (phonemic fluency) > fixation | Verbal | Energization |
| Kircher et al., 2011 | 1 | https://doi.org/10.1016/j.brainres.2011.03.054 | 15 | 12 | phonemic VF > silent rest | Verbal | Energization |
| Audenaert et al., 2000 | 1 | 10.1007/s002590000351 | 10 | 7 | phonemic VF > recite month/days | Verbal | Energization |
| Weiss et al., 2003 | 1 | 10.1016/j.schres.2004.01.010 | 20 | 8 | phonemic VF > silent rest | Verbal | Energization |
| Dye et al., 1999 | 1 | 10.1192/bjp.175.4.367 | 10 | 10 | phonemic VF > repeat noun/verb | Verbal | Energization |
| Phelps et al., 1997 | 1 | 10.1097/00001756-199701200-00036 | 11 | 8 | phonemic VF > repeat cue word | Verbal | Energization |
| Schlösser et al., 2003 | 1 | 10.1136/jnnp.64.4.492 | 6 | 23 | female - phonemic VF > count forward | Verbal | Energization |
| Schlösser et al., 2003 | 2 | 10.1136/jnnp.64.4.492 | 6 | 18 | male - phonemic VF > count forward | Verbal | Energization |
| Abrahams et al., 2003 | 1 | 10.1002/hbm.10126 | 18 | 22 | phonemic VF > repeat word rest | Verbal | Energization |
| Lurito et al., 2000 | 1 | https://doi.org/10.1002/1097-0193(200007)10:3<99::aid-hbm10>3.0.co;2-q | 5 | 15 | phonemic VF > fixation of a symbol | Verbal | Energization |
| Brammer et al., 2000 | 1 | 10.1016/S0730-725X(97)00135-5 | 6 | 6 | phonemic VF > repeat word rest | Verbal | Energization |
| Hutchinson et al., 1999 | 1 | 10.1016/S0730-725X(99)00093-4 | 12 | 6 | phonemic VF > silently count forward | Verbal | Energization |
| Fu et al., 2001 | 1 | 10.1016/S1053-8119(02)91189-4 | 11 | 29 | phonemic VF > repeat word rest | Verbal | Energization |
| Okada et al., | 1 | 10.1159/000068871 | 10 | 8 | phonemic VF > repeat word rest | Verbal | Energization |

| Reference and year | Contrast # | DOI | N | Foci | Task name, brief description and contrast | Input modality | Subcomponent |
|---|---|---|---|---|---|---|---|
| 2003 | | | | | | | |
| Curtis et al., 1998 | 1 | 10.1176/ajp.155.8.1056 | 5 | 9 | phonemic VF > repeat word rest | Verbal | Energization |
| Nosarti et al., 2009 | 1 | 10.1016/j.neuroimage.2009.04.041 | 26 | 4 | phonemic VF > repeat word rest | Verbal | Energization |
| Halari et al., 2006 | 1 | 10.1007/s00221-005-0118-7 | 9 | 12 | males - phonemic VF > repeat word rest | Verbal | Energization |
| Halari et al., 2006 | 2 | 10.1007/s00221-005-0118-7 | 10 | 3 | females - phonemic VF > repeat word rest | Verbal | Energization |
| Weiss et al., 2004 | 1 | 10.1016/j.schres.2004.01.010 | 9 | 8 | females - phonemic VF > repeat word rest | Verbal | Energization |
| Bonelli et al., 2011 | 1 | https://doi.org/10.1016/j.eplepsyres.2011.04.007 | 22 | 4 | phonemic VF > fixation cross | Verbal | Energization |
| Meinzer et al., 2009 | 1 | 10.1162/jocn.2009.21219 | 16 | 7 | phonemic VF > repeat word pause | Verbal | Energization |
| Ragland et al., 2008 | 1 | 10.1016/j.schres.2007.11.017 | 14 | 7 | semantic VF > repeat word rest | Verbal | Energization |
| Kircher et al., 2011 | 2 | https://doi.org/10.1016/j.brainres.2011.03.054 | 15 | 15 | semantic VF > silent rest | Verbal | Energization |
| Krug et al., 2011 | 1 | 10.1002/hbm.21005 | 91 | 5 | semantic VF > read aloud nouns | Verbal | Energization |
| Basho et al., 2007 | 1 | 10.1016/j.neuropsychologia.2007.01.007 | 12 | 6 | semantic VF > read the word nothing | Verbal | Energization |
| Gaillard et al., 2003 | 1 | https://doi.org/10.1002/hbm.10091 | 29 | 14 | semantic VF > silent rest | Verbal | Energization |
| Meinzer et al., 2009 | 1 | 10.1162/jocn.2009.21219 | 16 | 7 | semantic VF > repeat word pause | Verbal | Energization |
| Meinzer et al., 2012 | 1 | 10.1016/j.neurobiolaging.2010.06.020 | 20 | 8 | semantic VF > repeat word rest | Verbal | Energization |
| Audenaert et al., 2000 | 2 | 10.1007/s002590000351 | 10 | 6 | semantic VF > recite month/days | Verbal | Energization |
| Rodriguez-Aranda et al., 2020 | 1 | https://doi.org/10.3389/fnhum.2020.00203 | 15 | 3 | young group semantic fluency > rest (controlled for MMSE score and age) | Verbal | Energization |
| Li et al., 2017 | 1 | https://doi.org/10.3389/fnbeh.2017.00131 | 21 | 9 | covert semantic fluency > fixation | Verbal | Energization |
| Li et al., 2017 | 2 | https://doi.org/10.3389/fnbeh.2017.00131 | 21 | 11 | covert phonemic fluency > fixation | Verbal | Energization |

| Reference and year | Contrast # | DOI | N | Foci | Task name, brief description and contrast | Input modality | Subcomponent |
|---|---|---|---|---|---|---|---|
| Gawda and Szepietowska., 2016 | 1 | https://doi.org/10.3389/fnbeh.2016.00010 | 35 | 14 | phonemic fluency > fixation | Verbal | Energization |
| Gawda and Szepietowska., 2016 | 2 | https://doi.org/10.3389/fnbeh.2016.00010 | 35 | 20 | semantic fluency > fixation | Verbal | Energization |
| Marsolais et al., 2015 | 1 | 10.1016/j.bandl.2014.10.010 | 28 | 16 | main effect of task (semantic/orthographic fluency > name month of year) | Verbal | Energization |
| Gourovitch et al., 2000 | 1 | https://psycnet.apa.org/doi/10.1037/0894-4105.14.3.353 | 18 | 11 | phonemic fluency task > control | Verbal | Energization |
| Gourovitch et al., 2000 | 2 | https://psycnet.apa.org/doi/10.1037/0894-4105.14.3.353 | 18 | 10 | semantic fluency task > control | Verbal | Energization |
| Haller et al., 2005 | 1 | https://doi.org/10.1016/j.neuropsychologia.2004.09.007 | 15 | 9 | SVO sentence generation > word reading | Verbal | Energization |
| Haller et al., 2005 | 2 | https://doi.org/10.1016/j.neuropsychologia.2004.09.007 | 15 | 7 | SVO sentence generation > sentence reading | Verbal | Energization |
| Kemeny et al., 2005 | 1 | https://doi.org/10.1002/hbm.20078 | 6 | 10 | sentence that must contain a verb > syllable | Verbal | Energization |
| Troiani et al., 2008 | 1 | https://doi.org/10.1016/j.neuroimage.2007.12.002 | 15 | 5 | narrative (unconstrained) > story viewing (during nonsense word production) | Nonverbal | Energization |
| Tremblay et al., 2011 | 1 | https://doi.org/10.3389/fpsyg.2011.00253 | 20 | 18 | describe an object (line drawing) > picture viewing | Nonverbal | Energization |
| Grande et al., 2012 | 1 | https://doi.org/10.1016/j.neuroimage.2012.03.087 | 18 | 7 | unimpaired language (picture description task) > baseline | Nonverbal | Energization |
| Menenti et al., 2012a | 1 | https://doi.org/10.1016/j.bandl.2012.04.012 | 20 | 30 | picture description task: semantics, words, syntax (table 1) | Nonverbal | Energization |
| Menenti et al., 2012b | 1 | 10.3389/fpsyg.2011.00384 | 24 | 12 | sentence suppression and reference suppression | Nonverbal | Energization |
| Geranmayeh et al., 2014 | 1 | 10.1523/JNEUROSCI.0428-14.2014 | 24 | 12 | speech > counting | Nonverbal | Energization |
| Schonberger et al., 2014 | 1 | 10.3389/fpsyg.2014.00246 | 15 | 9 | simple complete > simple incomplete | Nonverbal | Energization |
| Matchin & Hickok, 2016 | 1 | 10.3389/fpsyg.2016.00241 | 20 | 4 | produce sentences > read word lists | Nonverbal | Energization |
| Rubia et al., 2006 | 1 | 10.1002/hbm.20237 | 22 | 9 | Simon: Adults- incongruent > congruent | nonverbal | Monitoring |
| Schilbach et | 1 | 10.1093/scan/nsq067 | 23 | 13 | Social gaze: Incongruent > congruent | nonverbal | Monitoring |

| Reference and year | Contrast # | DOI | N | Foci | Task name, brief description and contrast | Input modality | Subcomponent |
|---|---|---|---|---|---|---|---|
| al., 2011 | | | | | | | |
| Dong et al., 2014 | 1 | 10.1186/1744-9081-10-4 | 30 | 1 | Stroop (color-word; for two consecutive trials, either in repeated/different condition) : Incongruent (congruent – incongruent) > congruent (incongruent – incongruent) | verbal | Monitoring |
| Van Veen & Carter, 2005 | 1 | 10.1016/j.neuroimage.2005.04.042 | 14 | 6 | Stroop (color-word): Incongruent > congruent | verbal | Monitoring |
| Okayasu et al., 2023 | 1 | 10.1038/s41467-022-35397-w | 33 | 76 | Stroop (color-word): Incongruent > congruent | verbal | Monitoring |
| Verdolini et al., 2023 | 1 | 10.1016/j.jad.2023.02.132 | 28 | 4 | Stroop (number-word) : Incongruent > congruent | verbal | Monitoring |
| Jalalvandi et al., 2020 | 1 | 10.31661/jbpe.v0i0.2003-1084 | 20 | 6 | Stroop (color-word): Incongruent > congruent | verbal | Monitoring |
| Duggirala et al., 2022 | 1 | 10.3758/s13415-022-01025-9 | 23 | 9 | Flanker: Incongruent > congruent | nonverbal | Monitoring |
| Bernal & Altman 2009 | 1 | 10.1080/00207450802333029 | 18 | 40 | Stroop (color-word): Incongruent > congruent | verbal | Monitoring |
| Aarts et al., 2008 | 1 | 10.1523/JNEUROSCI.4400-07.2008 | 12 | 32 | Stroop (color-word) : Incongruent > congruent | verbal | Monitoring |
| Aarts & Roelofs, 2011 | 1 | 10.1162/jocn.2010.21435 | 20 | 19 | Stroop with certainty altered between trials (arrow-word) : Incongruent > congruent | verbal | Monitoring |
| Bastern et al., 2011 | 1 | 10.1162/jocn_a_00003 | 46 | 11 | Stroop (color-word): Incongruent > congruent | verbal | Monitoring |
| Becker et al., 2008 | 1 | 10.1038/sj.npp.1301673 | 17 | 9 | Stroop (color-word): Incongruent > congruent | verbal | Monitoring |
| Brass et al., 2005 | 1 | 10.1016/j.neuropsychologia.2004.06.018 | 10 | 7 | Stroop (color-word) : Incongruent > congruent | verbal | Monitoring |
| Coderre et al., 2008 | 1 | 10.1016/j.bandl.2008.01.011 | 9 | 8 | Stroop (Japanese: kana-kanji) : Incongruent > congruent | verbal | Monitoring |
| Compton et al., 2003 | 1 | 10.3758/CABN.3.2.81 | 12 | 5 | Stroop (color-word, also with semantically unrelated words) : More interference (color-word) > neutral (non-color-word) | verbal | Monitoring |
| Fan et al., 2003 | 1 | 10.1006/nimg.2002.1319 | 12 | 14 | Stroop (color-word) : Incongruent > congruent | verbal | Monitoring |
| Grandjean et | 1 | 10.1016/j.neuropsychologia.2013.02.015 | 25 | 7 | Stroop (color-word) : Incongruent > | verbal | Monitoring |

| Reference and year | Contrast # | DOI | N | Foci | Task name, brief description and contrast | Input modality | Subcomponent |
|---|---|---|---|---|---|---|---|
| al., 2013 | | | | | congruent | | |
| Jeong et al., 2005 | 1 | 10.1016/j.pscychresns.2004.01.008 | 10 | 9 | Stroop (color-word): Incongruent > congruent | verbal | Monitoring |
| Kerns et al., 2005 | 1 | 10.1176/appi.ajp.162.10.1833 | 13 | 13 | Stroop (color-word): Incongruent > congruent | verbal | Monitoring |
| Kim et al., 2010 | 1 | 10.1016/j.neulet.2010.04.019 | 12 | 5 | Stroop (arrow-word) : perceptual Incongruent > perceptual congruent | verbal | Monitoring |
| Kim et al., 2010 | 2 | 10.1016/j.neulet.2010.04.019 | 12 | 6 | Stroop (arrow-word) : response Incongruent > response congruent | verbal | Monitoring |
| Kim et al., 2012 | 1 | 10.1016/j.brainres.2012.06.013 | 16 | 22 | Stroop (color-word) : Incongruent > congruent | verbal | Monitoring |
| Mayer et al., 2012 | 1 | 10.1002/hbm.21405 | 24 | 14 | Stroop (number-word): Incongruent > congruent | verbal | Monitoring |
| Mead et al., 2002 | 1 | 10.1017/S1355617702860015 | 18 | 1 | Stroop (color-word) : Incongruent > congruent | verbal | Monitoring |
| Melcher & Gruber, 2006 | 1 | 10.1016/j.brainres.2006.08.120 | 9 | 8 | Stroop (font size-word): Incongruent > congruent | verbal | Monitoring |
| Milham et al., 2002 | 1 | 10.1006/brcg.2001.1501 | 12 | 6 | Stroop (color-word): Incongruent > congruent and neutral (old, young) | verbal | Monitoring |
| Milham et al., 2005 | 1 | 10.1002/hbm.20110 | 18 | 23 | Stroop (color-word): Incongruent > congruent | verbal | Monitoring |
| Nakao et al., 2005 | 1 | 10.1016/j.pscychresns.2004.12.004 | 14 | 27 | Stroop (color-word): Incongruent > congruent | verbal | Monitoring |
| Ochsner et al., 2005 | 1 | 10.1162/jocn.2009.21129 | 16 | 16 | Flanker (with mood-neutral word): Incongruent > congruent (neutral) | verbal | Monitoring |
| Ochsner et al., 2005 | 2 | 10.1162/jocn.2009.21129 | 16 | 13 | Flanker (with affective word): incongruent > congruent (affective) | verbal | Monitoring |
| Peterson et al., 1999 | 1 | 10.1016/S0006-3223(99)00056-6 | 34 | 40 | Stroop (color-word): Incongruent > congruent | verbal | Monitoring |
| Piai et al., 2013 | 1 | 10.3389/fnhum.2013.00832 | 23 | 5 | Stroop (color-word): Incongruent > congruent | verbal | Monitoring |
| Purmann et al., 2015 | 1 | 10.3389/fnhum.2015.00088 | 18 | 7 | Stroop (color-word): Incongruent (incompatible) > congruent (compatible) | verbal | Monitoring |
| Roelofs et al., 2006 | 1 | 10.1073/pnas.0606265103 | 12 | 11 | Stroop (arrow-word) : Incongruent > congruent | verbal | Monitoring |
| Terry et al., 2012 | 1 | 10.3109/02699052.2012.722259 | 20 | 5 | Stroop (color-word): Incongruent > congruent | verbal | Monitoring |
| Ye & Zhou, | 1 | 10.1016/j.neuroimage.2009.06.032 | 19 | 14 | Stroop (color-word) : Incongruent > | verbal | Monitoring |

| Reference and year | Contrast # | DOI | N | Foci | Task name, brief description and contrast | Input modality | Subcomponent |
|---|---|---|---|---|---|---|---|
| 2009 | | | | | congruent | | |
| Zoccatelli et al., 2010 | 1 | 10.1007/s00221-010-2433-x | 10 | 13 | Stroop (color-word): Incongruent > congruent | verbal | Monitoring |
| Zysset et al., 2001 | 1 | 10.1006/nimg.2000.0665 | 9 | 4 | Stroop (color-word): Incongruent > congruent | verbal | Monitoring |
| Kelley et al., 2013 | 2 | 10.1016/j.neuroimage.2013.04.094 | 15 | 1 | Flanker: Incongruent > congruent | nonverbal | Monitoring |
| Yoon et al., 2012 | 1 | 10.1162/jocn_a_00256 | 17 | 2 | View videos of objects used in congruent or incongruent actions: Incongruent > congruent | nonverbal | Monitoring |
| Berron et al.., 2015 | 1 | 10.1371/journal.pone.0120582 | 24 | 10 | Flanker: Incongruent > congruent | nonverbal | Monitoring |
| Böckler et al., 2016 | 1 | 10.1080/17588928.2015.1053442 | 21 | 7 | Eye gaze (flanker variant) : Incongruent > congruent | nonverbal | Monitoring |
| Christakou et al., 2009 | 1 | 10.1016/j.neuroimage.2009.06.070 | 63 | 5 | Simon: Incongruent > congruent | nonverbal | Monitoring |
| Fan et al., 2003 | 1 | 10.1006/nimg.2002.1319 | 12 | 14 | Flanker: Incongruent > congruent | nonverbal | Monitoring |
| Fan et al., 2012 | 1 | 10.1002/brb3.90 | 12 | 34 | Flanker: Incongruent > congruent | nonverbal | Monitoring |
| Frühholz et al., 2011 | 1 | 10.1016/j.neuroimage.2010.07.071 | 24 | 15 | Simon and Flanker (conjunction): Incongruent colour and position > congruent colour and position | nonverbal | Monitoring |
| Georgiou-Karistianis et al., 2012 | 1 | 10.1016/j.bandc.2012.02.011 | 14 | 27 | Simon: Incongruent > congruent | nonverbal | Monitoring |
| Hazeltine et al., 2000 | 1 | 10.1162/089892900563984 | 8 | 4 | Flanker : Incongruent > congruent | nonverbal | Monitoring |
| Kellermann et al., 2011 | 1 | 10.1016/j.brainres.2011.02.023 | 15 | 9 | Flanker: Incongruent > congruent | nonverbal | Monitoring |
| Kerns, 2006 | 1 | 10.1016/j.neuroimage.2006.06.012 | 26 | 5 | Simon: Incongruent > all trials | nonverbal | Monitoring |
| King et al., 2012 | 1 | 10.1523/JNEUROSCI.0934-12.2012 | 25 | 9 | Flanker: Incongruent > congruent | nonverbal | Monitoring |
| Korsch et al., 2014 | 1 | 10.3389/fnagi.2014.00057 | 19 | 9 | Flanker: Incongruent > congruent | nonverbal | Monitoring |
| Leibovich et al., 2016 | 1 | 10.1162/jocn_a_00887 | 19 | 2 | Simon (variant): Incongruent > congruent | nonverbal | Monitoring |
| Liu et al., | 1 | 10.1016/j.neuroimage.2004.02.033 | 11 | 34 | Simon: Incongruent > congruent | nonverbal | Monitoring |

| Reference and year | Contrast # | DOI | N | Foci | Task name, brief description and contrast | Input modality | Subcomponent |
|---|---|---|---|---|---|---|---|
| 2004 | | | | | | | |
| Rusted et al., 2013 | 1 | 10.1016/j.neuroimage.2012.10.010 | 40 | 5 | Rapid visual processing : Incongruent cue > congruent cue | nonverbal | Monitoring |
| Siemann et al., 2016 | 1 | 10.1016/j.neuroimage.2016.05.036 | 19 | 10 | Flanker: Incongruent > congruent | nonverbal | Monitoring |
| Ullsperger & von Cramon, 2001 | 1 | 10.1006/nimg.2001.0935 | 9 | 34 | Flanker : Incompatible > compatible | nonverbal | Monitoring |
| vel Grajewska et al., 2011 | 1 | 10.1016/j.brainres.2011.09.022 | 18 | 6 | Flanker: Incompatible > compatible | nonverbal | Monitoring |
| Zhu et al., 2010 | 1 | 10.1016/j.neuroimage.2009.12.087 | 22 | 8 | Flanker: Incongruent > congruent (young only) | nonverbal | Monitoring |
| Zoccatelli et al., 2010 | 2 | 10.1007/s00221-010-2433-x | 10 | 13 | Stroop (arrow direction- arrow position): Incongruent > congruent | nonverbal | Monitoring |
| Mennigen et al., 2014 | 1 | 10.1016/j.neuropsychologia.2014.06.022 | 185 | 14 | Flanker: Incongruent > congruent | nonverbal | Monitoring |
| Rodehacke et al., 2014 | 1 | 10.1371/journal.pone.0088957 | 213 | 9 | Flanker : Incongruent > congruent | nonverbal | Monitoring |
| Sylvester et al., 2003 | 1 | 10.1016/S0028-3932(02)00167-7 | 14 | 11 | Flanker with stimulus-response incompatible trials : Incompatible > compatible | nonverbal | Monitoring |
| Sylvester et al., 2003 | 2 | 10.1016/S0028-3932(02)00167-7 | 14 | 12 | Flanker: Incongruent > congruent | nonverbal | Monitoring |
| Mulder et al., 2008 | 1 | 10.1097/chi.0b013e31815a56dc | 12 | 6 | Go/no-go variant (with unexpected/expected times) : Unexpected stimulus > expected time | nonverbal | Monitoring |
| Mulder et al., 2008 | 2 | 10.1097/chi.0b013e31815a56dc | 12 | 4 | Go/no-go variant (with unexpected/expected times) : Expected stimulus > unexpected time | nonverbal | Monitoring |
| Mulder et al., 2008 | 3 | 10.1097/chi.0b013e31815a56dc | 12 | 6 | Go/no-go variant (with unexpected/expected times) : Unexpected stimulus > unexpected time | nonverbal | Monitoring |
| Afyouni et al., 2021 | 1 | https://doi.org/10.1093/texcom/tgab057 | 19 | 7 | Anti-saccade: Adult- antisaccade > prosaccade | nonverbal | Monitoring |
| Gaillard et al., 2020 | 1 | https://doi.org/10.1016/j.bbr.2020.112586 | 48 | 5 | Stop-signal: Signal Inhibit (i.e., successful stop trial) > go | nonverbal | Monitoring |
| Cao & Cannon, 2021 | 1 | https://doi.org/10.1002%2Fhbm.25347 | 119 | 15 | Stop-signal: Correct response in stop signal > hit (i.e., successful concurrent | nonverbal | Monitoring |

| Reference and year | Contrast # | DOI | N | Foci | Task name, brief description and contrast | Input modality | Subcomponent |
|---|---|---|---|---|---|---|---|
| | | | | | cognitive control) | | |
| Swann et al., 2012 | 1 | https://doi.org/10.1016/j.neuroimage.2011.09.049 | 16 | 21 | Maybe Stop (MS)/No Stop (NS): Maybe stop_go > noStop_go | nonverbal | Monitoring |
| Bonnet et al., 2009 | 1 | https://doi.org/10.1002/hbm.20575 | 20 | 3 | Go/no-go: Initial go/no-go > tonic alertness | nonverbal | Monitoring |
| Bonnet et al., 2009 | 2 | https://doi.org/10.1002/hbm.20575 | 20 | 5 | Go/no-go : Complex go/no-go > tonic altertness | nonverbal | Monitoring |
| Braver et al., 2001 | 1 | https://doi.org/10.1093/cercor/11.9.825 | 14 | 11 | Go/no-go: Response inhibition > go | verbal | Monitoring |
| Evers et al., 2006 | 1 | https://doi.org/10.1007/s00213-006-0411-6 | 13 | 17 | Modified Go/no-go: No-go > go | verbal | Monitoring |
| Falconer et al., 2008 | 1 | https://www.jpn.ca/content/jpn/33/5/413.full.pdf | 23 | 3 | Go/no-go: No-go > go | nonverbal | Monitoring |
| Horn et al., 2003 | 1 | https://doi.org/10.1016/S0028-3932(03)00077-0 | 21 | 13 | Go/no-go: No-go > go | nonverbal | Monitoring |
| Kaladjian et al. (2009b) | 1 | https://doi.org/10.1016/j.pscychresns.2008.08.003 | 20 | 16 | Go/no-go: Correct no-go > correct go | nonverbal | Monitoring |
| Konishi et al., 1999 | 1 | https://doi.org/10.1093/brain/122.5.981 | 6 | 2 | Go/no-go: No-go > go | nonverbal | Monitoring |
| Matsuda et al., 2004 | 1 | https://doi.org/10.1016/j.pscychresns.2003.12.007 | 21 | 12 | Anti-saccade: Anti-saccade > visually guided saccade | nonverbal | Monitoring |
| Durston et al., 2003 | 1 | https://doi.org/10.1016/S0006-3223(02)01904-2 | 7 | 7 | Go/no-go: No-go > go | nonverbal | Monitoring |
| Tamm et al., 2002 | 1 | https://doi.org/10.1097/00004583-200210000-00013 | 19 | 4 | Go/no-go: No-go > go | verbal | Monitoring |
| Binter et al., 2019 | 1 | https://doi.org/10.1016/j.envint.2019.105163 | 71 | 21 | Go/no-go: Successful no-go > successful go | nonverbal | Monitoring |
| Booth et al., 2003 | 1 | https://doi.org/10.1016/S1053-8119(03)00404-X | 12 | 13 | Go/no-go: Adults- No-go > go | nonverbal | Monitoring |
| Cascio et al., 2015 | 1 | https://doi.org/10.1162/jocn_a_00693 | 37 | 8 | Go/no-go: Correct No-go > go (all participants) | verbal | Monitoring |
| Fitzgerald et al., 2008 | 1 | https://doi.org/10.1111/j.1469-7610.2008.01906.x | 11 | 12 | Anti-saccade: Correct anti-saccade > correct pro-saccade | nonverbal | Monitoring |
| Gooskens et al., 2019 | 1 | https://doi.org/10.1016/j.dcn.2018.11.004 | 53 | 11 | Stop-signal : Successful stopping > go | nonverbal | Monitoring |
| Lei et al., 2012 | 1 | https://doi.org/10.1002/hbm.21411 | 22 | 14 | Go/no-go: Go/no-go > go | verbal | Monitoring |

| Reference and year | Contrast # | DOI | N | Foci | Task name, brief description and contrast | Input modality | Subcomponent |
|---|---|---|---|---|---|---|---|
| Mechelli et al., 2009 | 1 | https://doi.org/10.1002/hbm.20818 | 69 | 8 | Go/no-go: No-go > go | nonverbal | Monitoring |
| Singh et al., 2010 | 1 | https://doi.org/10.1089/cap.2009.0004 | 22 | 2 | Go/no-go: No-go > go | verbal | Monitoring |
| Spielberg et al., 2015 | 1 | https://doi.org/10.1002/hbm.22838 | 63 | 7 | Go/no-go: No-go > go | verbal | Monitoring |
| Aichert et al., 2012 | 1 | https://doi.org/10.1111/j.1469-8986.2011.01306.x | 54 | 12 | Anti-saccade: Anti-saccade > pro-saccade | nonverbal | Monitoring |
| Boehler et al., 2010 | 1 | https://doi.org/10.1016/j.neuroimage.2010.04.276 | 15 | 30 | Stop-signal: Successful stop > go (stop-relevant blocks) | nonverbal | Monitoring |
| Brown et al., 2007 | 1 | https://doi.org/10.1152/jn.00460.2007 | 11 | 11 | Anti-saccade: Anti-saccade response > pro-saccade response | nonverbal | Monitoring |
| Brown et al., 2012 | 1 | https://doi.org/10.1016/j.neuroimage.2012.06.056 | 20 | 17 | Go/no-go: No-go > go | nonverbal | Monitoring |
| Burke et al., 2011 | 1 | https://doi.org/10.1162/jocn_a_00025 | 11 | 17 | Go/no-go: No-go > go | nonverbal | Monitoring |
| Cai & Leung, 2009 | 1 | https://doi.org/10.1016/j.brainres.2008.12.073 | 12 | 8 | Stop-signal: Successful stop > go (colour task) | nonverbal | Monitoring |
| Cai & Leung, 2009 | 2 | https://doi.org/10.1016/j.brainres.2008.12.073 | 12 | 14 | Stop-signal: Successful stop > go (orientation task) | nonverbal | Monitoring |
| Cai & Leung, 2011 | 1 | https://doi.org/10.1371/journal.pone.0020840 | 23 | 21 | Stop-signal: Successful stop > go (SST) | nonverbal | Monitoring |
| Chen et al., 2015 | 1 | https://doi.org/10.1111/pcn.12224 | 15 | 7 | Go/no-g: No-go > go | nonverbal | Monitoring |
| Chikazoe et al., 2008 | 1 | https://doi.org/10.1093/cercor/bhn065 | 25 | 51 | Go/no-go: No-go > frequent go | nonverbal | Monitoring |
| Chikazoe et al., 2008 | 2 | https://doi.org/10.1093/cercor/bhn065 | 25 | 52 | Go/no-go: No-go > infrequent go | nonverbal | Monitoring |
| Chiu & Egner, 2015 | 1 | https://doi.org/10.1523/JNEUROSCI.0519-15.2015 | 21 | 8 | Go/no-go : No-go > go | nonverbal | Monitoring |
| Congdon et al., 2010 | 1 | https://doi.org/10.1016/j.neuroimage.2010.06.062 | 126 | 19 | Stop-signal: StopInhibit > go | nonverbal | Monitoring |
| Coxon et al., 2016 | 1 | https://doi.org/10.1093/cercor/bhu165 | 20 | 28 | Stop-signal: Young- StopInhibit > go | nonverbal | Monitoring |
| Cummins et al., 2012 | 1 | https://doi.org/10.1038/mp.2011.104 | 50 | 5 | Stop-signal: Stop (successful inhibition) > go | verbal | Monitoring |
| Eijsker et al., 2019 | 1 | https://doi.org/10.3389/fpsyt.2019.00765 | 21 | 1 | Stop-signal: Successful inhibition > correct go | nonverbal | Monitoring |

| Reference and year | Contrast # | DOI | N | Foci | Task name, brief description and contrast | Input modality | Subcomponent |
|---|---|---|---|---|---|---|---|
| Fedota et al., 2014 | 1 | https://doi.org/10.1016/j.neuropsychologia.2013.12.022 | 16 | 9 | Go/no-go: No-go > go | verbal | Monitoring |
| Fuentes-Claramonte et al., 2016 | 1 | https://doi.org/10.1016/j.biopsycho.2016.01.001 | 57 | 16 | Go/no-go: No-go > frequent go | nonverbal | Monitoring |
| Fuentes-Claramonte et al., 2016 | 2 | https://doi.org/10.1016/j.biopsycho.2016.01.001 | 57 | 8 | Go/no-go: No-go > infrequent go | nonverbal | Monitoring |
| Gavazzi et al., 2019 | 1 | https://doi.org/10.1111/ejn.14301 | 36 | 18 | Go/no-go: No-go repetitions > go repetitions | verbal | Monitoring |
| Hendrick et al., 2010 | 1 | https://doi.org/10.1371/journal.pone.0013155 | 60 | 18 | Stop-signal: Stop > go | verbal | Monitoring |
| Hughes et al., 2014 | 1 | https://doi.org/10.1111/ejn.12497 | 12 | 15 | Stop-signal: Stop > go | verbal | Monitoring |
| Iannaccone et al., 2015 | 1 | https://doi.org/10.1016/j.neuroimage.2014.10.028 | 15 | 13 | Flanker: High- and low conflict correct > no conflict correct | nonverbal | Monitoring |
| Jahfari et al., 2011 | 1 | https://doi.org/10.1523/JNEUROSCI.5253-10.2011 | 20 | 7 | Combined Simon and Stop-signal: Successful stop > go | nonverbal | Monitoring |
| Jahfari et al., 2012 | 1 | https://doi.org/10.1523/JNEUROSCI.0902-12.2012 | 16 | 8 | Stop-signal: Successful stop (high, low) > go none | nonverbal | Monitoring |
| Jahfari et al., 2015 | 1 | https://doi.org/10.1162/jocn_a_00792 | 23 | 5 | Stop-signal: Successful stop > go | nonverbal | Monitoring |
| Köhler et al., 2018 | 1 | https://doi.org/10.1016/j.neuropsychologia.2018.08.003 | 33 | 23 | Go/no-go: Correct no-go > correct go | nonverbal | Monitoring |
| Kolodny et al., 2017 | 1 | https://doi.org/10.1016/j.cortex.2016.12.012 | 20 | 5 | Go/no-go: No-go > go (rare no-go condition) | nonverbal | Monitoring |
| Lavallee et al., 2014 | 1 | https://doi.org/10.1371/journal.pone.0096159 | 21 | 9 | Stop-signal: Stim1 > go1 | nonverbal | Monitoring |
| Leung & Cai, 2007 | 1 | https://doi.org/10.1523/JNEUROSCI.2837-07.2007 | 12 | 5 | Stop-signal: Stop > go (saccade and manual) | nonverbal | Monitoring |
| Marco-Pallares et al., 2008 | 1 | https://doi.org/10.1162/jocn.2008.20117 | 10 | 10 | Combined Flanker and Stop-signal : Successful inhibition trials > correct responses | verbal | Monitoring |
| McNab et al., 2008 | 1 | https://doi.org/10.1016/j.neuropsychologia.2008.04.023 | 11 | 6 | Go/no-go: No-go > go | nonverbal | Monitoring |

| Reference and year | Contrast # | DOI | N | Foci | Task name, brief description and contrast | Input modality | Subcomponent |
|---|---|---|---|---|---|---|---|
| Messel et al., 2019 | 1 | https://doi.org/10.1016/j.neuropsychologia.2019.107220 | 28 | 8 | Stop-signal: Stop > go | nonverbal | Monitoring |
| Ness & Beste, 2013 | 1 | https://doi.org/10.1016/j.neuropsychologia.2013.09.032 | 13 | 4 | Stop-change: SCD 0 > go | nonverbal | Monitoring |
| Ness & Beste, 2013 | 2 | https://doi.org/10.1016/j.neuropsychologia.2013.09.032 | 13 | 11 | Stop-change: SCD 300 > go | nonverbal | Monitoring |
| Penfold et al., 2015 | 1 | https://doi.org/10.1016/j.pscychresns.2014.11.005 | 20 | 30 | Go/no-go: No-go > go | verbal | Monitoring |
| Rothmayr et al., 2011 | 1 | https://doi.org/10.1016/j.neuroimage.2010.12.052 | 12 | 4 | Go/no-go: No-go > go | nonverbal | Monitoring |
| Sagaspe et al., 2011 | 1 | https://doi.org/10.1016/j.neuroimage.2011.01.027 | 12 | 25 | Stop-signal: (StopInhibit + StopRespond) > go | nonverbal | Monitoring |
| Sebastian et al., 2012 | 1 | https://doi.org/10.1016/j.pscychresns.2012.02.010 | 24 | 19 | Go/no-go: No-go > go | verbal | Monitoring |
| Sebastian et al., 2013 | 1 | https://doi.org/10.1016/j.neuroimage.2012.09.020 | 21 | 17 | Hybrid response inhibition (combined Simon, Go/no-go, Stop-signal) : No-go > congruent go | nonverbal | Monitoring |
| Sebastian et al., 2013 | 2 | https://doi.org/10.1016/j.neuroimage.2012.09.020 | 24 | 25 | Go/no-go: No-go > go | verbal | Monitoring |
| Sebastian et al., 2016 | 1 | https://doi.org/10.1007/s00429-015-0994-y | 28 | 13 | Stop-signal: Stop > go | nonverbal | Monitoring |
| Shafritz et al., 2015 | 1 | https://doi.org/10.1016/j.pnpbp.2015.03.001 | 15 | 5 | Go/no-go (standard version): ‘X’ no-go > letter go; | verbal | Monitoring |
| Shafritz et al., 2015 | 2 | https://doi.org/10.1016/j.pnpbp.2015.03.001 | 15 | 6 | Go/no-go (emotional version): Happy no-go > neutral go | nonverbal | Monitoring |
| Shafritz et al., 2015 | 3 | https://doi.org/10.1016/j.pnpbp.2015.03.001 | 15 | 5 | Go/no-go (emotional version): Fear no-go > neutral go | nonverbal | Monitoring |
| Shafritz et al., 2015 | 4 | https://doi.org/10.1016/j.pnpbp.2015.03.001 | 15 | 8 | Go/no-go (emotional version): Happy no-go > fear go | nonverbal | Monitoring |
| Shafritz et al., 2015 | 5 | https://doi.org/10.1016/j.pnpbp.2015.03.001 | 15 | 7 | Go/no-go (emotional version): Fear no-go > happy go | nonverbal | Monitoring |
| Tabu et al., 2011 | 1 | https://doi.org/10.1016/j.neures.2011.03.007 | 13 | 6 | Stop-signal: Stop-success > go | nonverbal | Monitoring |
| Tabu et al., 2012 | 1 | https://doi.org/10.1016/j.neuroimage.2011.10.092 | 13 | 15 | Stop-signal: Stop-success > go (hand task) | nonverbal | Monitoring |
| Tabu et al., 2012 | 2 | https://doi.org/10.1016/j.neuroimage.2011.10.092 | 13 | 14 | Stop-signal: Stop-success > go (foot task) | nonverbal | Monitoring |

| Reference and year | Contrast # | DOI | N | Foci | Task name, brief description and contrast | Input modality | Subcomponent |
|---|---|---|---|---|---|---|---|
| Talanow et al., 2020 | 1 | https://doi.org/10.1007/s11682-018-9972-3 | 19 | 3 | Go/no-go: No-go > go | nonverbal | Monitoring |
| Talanow et al., 2020 | 2 | https://doi.org/10.1007/s11682-018-9972-3 | 21 | 17 | Anti-saccade: Certain anti-saccade > certain pro-saccade (pro-saccade/anti-saccade task) | nonverbal | Monitoring |
| Townsend et al., 2012 | 1 | https://doi.org/10.1111/j.1399-5618.2012.01020.x | 30 | 24 | Go/no-go: No-go > go | verbal | Monitoring |
| Van Eijk et al., 2015 | 1 | https://doi.org/10.1016/j.pscychresns.2015.09.017 | 18 | 5 | Go/no-go: No-go > go | verbal | Monitoring |
| Van Eijk et al., 2015 | 2 | https://doi.org/10.1016/j.pscychresns.2015.09.017 | 25 | 6 | Hybrid response inhibition task (combined Simon, Go/no-go, and Stop-Signal) : No-go > go | nonverbal | Monitoring |
| Walther et al., 2010 | 1 | https://doi.org/10.1097/WNR.0b013e328335640f | 17 | 15 | Go/no-go : No-go > go (across both auditory and visual modalities- i.e., conjunction analysis) | nonverbal | Monitoring |
| Weafer et al., 2019 | 1 | https://doi.org/10.1016/j.neuroimage.2019.04.021 | 44 | 15 | Stop-signal: StopInhibit > go | nonverbal | Monitoring |
| Xu et al., 2015 | 1 | https://doi.org/10.3389/fnhum.2015.00034 | 18 | 14 | Stop-signal: Stop-inhibit > StGo | nonverbal | Monitoring |
| Xue et al., 2008 | 1 | https://doi.org/10.1093/cercor/bhm220 | 15 | 13 | Stop-signal: StopInhibit > go (manual) | verbal | Monitoring |
| Xue et al., 2008 | 2 | https://doi.org/10.1093/cercor/bhm220 | 15 | 12 | Stop-signal: StopInhibit > go (letter naming) | verbal | Monitoring |
| Xue et al., 2008 | 3 | https://doi.org/10.1093/cercor/bhm220 | 15 | 8 | Stop-signal: StopInhibit > go (PW naming) | verbal | Monitoring |
| Zhao et al., 2019 | 1 | https://doi.org/10.1007/s11682-018-9868-2 | 20 | 48 | Stop-signal: StopInhibit > go (rested wakefulness) | nonverbal | Monitoring |
| Anderson et al., 2005 | 1 | https://doi.org/10.15288/jsa.2005.66.323 | 46 | 2 | Go/no-go: No-go > go | nonverbal | Monitoring |
| Bennett et al., 2009 | 1 | https://doi.org/10.1016/j.ntt.2009.03.003 | 11 | 8 | Go/no-go: Correct no-go > correct go | verbal | Monitoring |
| Carrion et al., 2008 | 1 | https://doi.org/10.1002/da.20346 | 14 | 31 | Go/no-go: No-go > go | verbal | Monitoring |
| Durston et al., 2006 | 1 | https://doi.org/10.1016/j.biopsych.2005.12.020 | 11 | 9 | Go/no-go: No-go > go | nonverbal | Monitoring |
| Hummer et al., 2010 | 1 | https://doi.org/10.1080/15213261003799854 | 22 | 2 | Go/no-go: No-go > go | verbal | Monitoring |
| Passarotti et al., 2010 | 1 | https://doi.org/10.1016/j.pscychresns.2009.07.002 | 15 | 5 | Stop-signal: Stop > go | nonverbal | Monitoring |

| Reference and year | Contrast # | DOI | N | Foci | Task name, brief description and contrast | Input modality | Subcomponent |
|---|---|---|---|---|---|---|---|
| Querne et al., 2008 | 1 | https://doi.org/10.1016/j.brainres.2008.07.066 | 10 | 14 | Go/no-go: Go/no-go > go | verbal | Monitoring |
| Rubia et al., 2006 | 1 | https://doi.org/10.1002/hbm.20237 | 23 | 11 | Go/no-go: Adults- No-go > go | nonverbal | Monitoring |
| Rubia et al., 2013 | 1 | https://doi.org/10.1016/j.neuroimage.2013.06.078 | 66 | 4 | Stop-signal: Successful inhibition > go | nonverbal | Monitoring |
| Schulz et al., 2004 | 1 | https://doi.org/10.1176/appi.ajp.161.9.1650 | 9 | 5 | Go/no-go: No-go > go | verbal | Monitoring |
| Ware et al., 2015 | 1 | https://doi.org/10.1016/j.bbr.2014.09.037 | 21 | 2 | Stop-signal: All-stop > go | verbal | Monitoring |
| Altshuler et al., 2005 | 1 | https://doi.org/10.1016/j.biopsych.2005.09.012 | 13 | 4 | Go/no-go: No-go > go | verbal | Monitoring |
| Asahi et al., 2004 | 1 | https://doi.org/10.1007/s00406-004-0488-z | 17 | 11 | Go/no-go: No-go > go | verbal | Monitoring |
| Bellgrove et al., 2004 | 1 | https://doi.org/10.1016/j.neuropsychologia.2004.05.007 | 42 | 19 | Go/no-go: Response inhibition > go | verbal | Monitoring |
| Chevrier et al., 2007 | 1 | https://doi.org/10.1002/hbm.20355 | 14 | 3 | Stop-signal: Successful stop > go | verbal | Monitoring |
| Dambacher et al., 2014 | 1 | https://doi.org/10.1093/scan/nsu077 | 15 | 3 | Go/no-go: No-go > go | nonverbal | Monitoring |
| Durston et al., 2002 | 1 | https://doi.org/10.1006/nimg.2002.1074 | 10 | 9 | Go/no-go: No-go > go | nonverbal | Monitoring |
| Fassbender et al., 2004 | 1 | https://doi.org/10.1016/j.cogbrainres.2004.02.007 | 18 | 4 | SART: Random SART correct inhibition > fixed SART correct inhibition | nonverbal | Monitoring |
| Garavan et al., 2006 | 1 | https://doi.org/10.1016/j.brainres.2006.03.029 | 71 | 20 | Go/no-go: Stop > go | verbal | Monitoring |
| Jamadar et al., 2010a | 1 | https://doi.org/10.1016/j.neuropsychologia.2009.12.034 | 18 | 43 | Go/no-go: No-go > informatively cued go | verbal and nonverbal | Monitoring |
| Kaladjian et al., 2007 | 1 | https://doi.org/10.1016/j.schres.2007.07.033 | 21 | 11 | Go/no-go: Correct no-go > correct go | verbal | Monitoring |
| Kaladjian et al., 2009 | 1 | https://doi.org/10.1016/j.pscychresns.2008.08.003 | 20 | 16 | Go/no-go: Correct no-go > correct go | verbal | Monitoring |
| Kiehl et al., 2000 | 1 | https://doi.org/10.1111/1469-8986.3720216 | 14 | 8 | Go/no-go: Correct rejects > go | verbal | Monitoring |
| Konishi et al., 2003 (Exp. 1) | 1 | https://doi.org/10.1523/JNEUROSCI.23-21-07776.2003 | 36 | 16 | Modified WCST: Inhibition > control | nonverbal | Monitoring |
| Lawrence et | 1 | https://doi.org/10.1002/hbm.20564 | 21 | 2 | Go/no-go: No-go > go | nonverbal | Monitoring |

| Reference and year | Contrast # | DOI | N | Foci | Task name, brief description and contrast | Input modality | Subcomponent |
|---|---|---|---|---|---|---|---|
| al., 2009 | | | | | | | |
| Liddle et al., 2001 | 1 | https://doi.org/10.1002/1097-0193(200102)12:2<100::AID-HBM1007>3.0.CO;2-6 | 16 | 23 | Go/no-go: Correct no-go > go | verbal | Monitoring |
| Maguire et al., 2003 | 1 | https://doi.org/10.1016/S1053-8119(03)00402-6 | 6 | 6 | Go/no-go: Go/no-go > go | nonverbal | Monitoring |
| Manoach et al., 2007 | 1 | https://doi.org/10.1523/JNEUROSCI.3662-06.2007 | 21 | 14 | Anti-saccade: Anti-saccade > pro-saccade (current) | nonverbal | Monitoring |
| Mazzola-Pomietto et al., 2009 | 1 | https://doi.org/10.1016/j.jpsychires.2008.05.004 | 16 | 7 | Warned equiprobable Go/no-go: Correct no-go > correct go | verbal | Monitoring |
| Meffert et al., 2016 | 1 | https://doi.org/10.1016/j.neuroimage.2015.11.029 | 22 | 15 | Go/no-go: No-go > go | nonverbal | Monitoring |
| Menon et al., 2001 | 1 | https://doi.org/10.1002/1097-0193(200103)12:3<131::AID-HBM1010>3.0.CO;2-C | 14 | 13 | Go/no-go: Go/no-go > go | verbal | Monitoring |
| Page et al., 2009 | 1 | https://doi.org/10.1016/j.pscychresns.2009.05.002 | 11 | 11 | Go/no-go: No-go > go | nonverbal | Monitoring |
| Pierce & McDowell, 2017 | 1 | https://doi.org/10.1016/j.bandc.2017.03.003 | 35 | 4 | Anti-saccade: Single anti-saccade > pro-saccade | nonverbal | Monitoring |
| Pierce & McDowell, 2017 | 2 | https://doi.org/10.1016/j.bandc.2017.03.003 | 35 | 2 | Anti-saccade: Mixed anti-saccade > pro-saccade | nonverbal | Monitoring |
| Rodríguez-Pujadas et al., 2014 | 1 | https://doi.org/10.1016/j.bandl.2014.03.003 | 33 | 7 | Stop-signal: Stop > go | verbal | Monitoring |
| Roth et al., 2007 | 1 | https://doi.org/10.1016/j.biopsych.2006.12.007 | 14 | 13 | Go/no-go: No-go > go | nonverbal | Monitoring |
| Wager et al., 2005 | 1 | https://doi.org/10.1016/j.neuroimage.2005.01.054 | 14 | 13 | Go/no-go: No-go > go | verbal | Monitoring |
| Zheng et al., 2008 | 1 | https://doi.org/10.1162/jocn.2008.20100 | 18 | 8 | Go/no-go: Successful inhibition (no-go) > go | verbal | Monitoring |
| Schmitz et al., 2006 | 1 | https://doi.org/10.1016/j.biopsych.2005.06.007 | 12 | 11 | Go/no-go: No-go > go | nonverbal | Monitoring |
| Pierce & McDowell, 2017 | 1 | 10.1016/j.bandc.2017.03.003 | 35 | 3 | Task switching: Switch > repeat | nonverbal | Task-setting |
| Sylvester et al., 2003 | 1 | 10.1016/S0028-3932(02)00167-7 | 14 | 14 | Counter-switching: Switch > non-switch (Experiment 1) | nonverbal | Task-setting |

| Reference and year | Contrast # | DOI | N | Foci | Task name, brief description and contrast | Input modality | Subcomponent |
|---|---|---|---|---|---|---|---|
| Rubia et al., 2006 | 1 | 10.1002/hbm.20237 | 22 | 7 | Task switching: Switch task: Adults- Switch > Repeat | nonverbal | Task-setting |
| Wendelken et al., 2012 | 1 | 10.1016/j.dcn.2012.02.001 | 36 | 6 | Rule switching: Switch > repeat | verbal instruction nonverbal stimuli | Task-setting |
| Aizawa et al., 2012 | 1 | 10.1053/j.gastro.2012.07.104 | 30 | 14 | WCST: First error feedback > correct feedback | nonverbal | Task-setting |
| Armbruster et al., 2012 | 1 | 10.1162/jocn_a_00286 | 20 | 15 | Task switching: Task switching > baseline | nonverbal | Task-setting |
| Becker et al., 2014 | 1 | 10.1016/j.neuroimage.2014.08.058 | 8 | 9 | Visual search: Dimension change > repeat | nonverbal | Task-setting |
| Becker et al., 2014 | 2 | 10.1016/j.neuroimage.2014.08.058 | 8 | 4 | Visual search: feature change > repeat | nonverbal | Task-setting |
| Brass & Cramon, 2004 | 1 | 10.1162/089892904323057335 | 14 | 4 | Oddball task: Cue switch > cue repetition | nonverbal | Task-setting |
| Christakou et al., 2009 | 1 | 10.1016/j.neuroimage.2009.06.070 | 63 | 4 | Meiran switch task: Switch > repeat | nonverbal | Task-setting |
| Crone et al., 2005 | 1 | 10.1093/cercor/bhi127 | 20 | 2 | visual cue switching: Univalent switches > repetitions | nonverbal | Task-setting |
| Crone et al., 2005 | 2 | 10.1093/cercor/bhi127 | 20 | 23 | visual cue switching: bivalent switches > repetitions | nonverbal | Task-setting |
| Dang et al., 2012 | 1 | 10.1162/jocn_a_00252 | 16 | 6 | flexible item selection: Object shift > no shift | nonverbal | Task-setting |
| De Baene & Brass, 2011 | 1 | 10.3758/s13415-011-0055-9 | 19 | 5 | cue and task switching: Task-switch > cue-repeat | nonverbal | Task-setting |
| De Baene et al., 2015 | 1 | 10.1162/jocn_a_00817 | 32 | 7 | language switching: Task switching > repeat | verbal | Task-setting |
| DiGirolamo et al., 2001 | 1 | 10.1097/00001756-200107030-00054 | 8 | 67 | task switching: Younger- Switch > non-switch | verbal | Task-setting |
| Dove et al., 2000 | 1 | 10.1016/S0926-6410(99)00029-4 | 16 | 13 | set switching: Task switch > task repetition | nonverbal | Task-setting |
| Dreher & Grafman, 2003 | 1 | 10.1093/cercor/13.4.329 | 8 | 13 | task switching: Task switching > baseline | verbal | Task-setting |
| Eich et al., 2016 | 1 | 10.1016/j.neuropsychologia.2016.08.009 | 62 | 2 | cued task switching: Young- Dual-task > single task (task switching and go/no-go) | verbal | Task-setting |
| Fuentes- | 1 | 10.1371/journal.pone.0123073 | 28 | 20 | nonlinguistic switching: Switch > repeat | nonverbal | Task-setting |

| Reference and year | Contrast # | DOI | N | Foci | Task name, brief description and contrast | Input modality | Subcomponent |
|---|---|---|---|---|---|---|---|
| Claramonte et al., 2015 | | | | | | | |
| Gazes et al., 2010 | 1 | 10.1016/j.bbr.2010.02.036 | 56 | 3 | dual task: Dual > single-task tracking | nonverbal | Task-setting |
| Gazes et al., 2012 | 1 | 10.1016/j.neuropsychologia.2012.09.039 | 47 | 6 | cued task switching: Task-switch > single-task | verbal | Task-setting |
| Graham et al., 2009 | 1 | 10.1016/j.neuroimage.2008.12.040 | 18 | 17 | set shifting: WCST: RSNF (negative feedback) > RSPF (positive feedback) | nonverbal | Task-setting |
| Greenberg et al., 2010 | 1 | 10.1523/JNEUROSCI.4248-09.2010 | 8 | 6 | attention shifting: Shift-colour and shift-location > hold (conjunction analysis) | nonverbal | Task-setting |
| Hedden & Gabriel, 2010 | 1 | 10.1016/j.neuroimage.2010.01.089 | 18 | 23 | global-local task: Incongruent shifting (IS) > neutral non-shifting (NN) | nonverbal | Task-setting |
| Hedden & Gabriel, 2010 | 2 | 10.1016/j.neuroimage.2010.01.089 | 18 | 21 | global-local task: neutral shifting (NS) > incongruent non-shifting (IN) | nonverbal | Task-setting |
| Heinen et al., 2017 | 1 | 10.1016/j.neuropsychologia.2017.02.024 | 14 | 21 | spatial switching: Shift > stay | nonverbal | Task-setting |
| Hodgson et al., 2015 | 1 | 10.3389/fnhum.2015.00486 | 15 | 9 | rule switching: Flip > hold (feedback event) | nonverbal | Task-setting |
| Ikkai & Curtis, 2008 | 1 | https://doi.org/10.1093/cercor/bhm171 | 14 | 16 | attention shifting: Shift > baseline (covert) | nonverbal | Task-setting |
| Ikkai & Curtis, 2008 | 2 | https://doi.org/10.1093/cercor/bhm171 | 14 | 16 | attention shifting: shift > baseline (overt) | nonverbal | Task-setting |
| Jamadar et al., 2010 | 1 | 10.1016/j.neuropsychologia.2009.12.034 | 18 | 9 | task switching: Switch > repeat | verbal | Task-setting |
| Jamadar et al., 2010b | 1 | https://doi.org/10.1016/j.neuroimage.2010.01.090 | 12 | 10 | task switching: Informatively cued switch > repeat | nonverbal | Task-setting |
| Konishi et al., 2002 | 1 | 10.1073/pnas.122644899 | 16 | 9 | WCST: Switch > non-switch | nonverbal | Task-setting |
| Korb et al., 2017 | 1 | 10.1523/JNEUROSCI.3289-16.2017 | 22 | 3 | Selective control Task: Switch > non-switch (task-set control) | nonverbal | Task-setting |
| Korb et al., 2017 | 2 | 10.1523/JNEUROSCI.3289-16.2017 | 22 | 3 | Selective control Task: alternate > repeat (response-set control) | nonverbal | Task-setting |
| Kuptsova et al., 2015 | 1 | https://doi.org/10.1134/S0362119715050084 | 70 | 12 | rule switching: Switch > control | nonverbal | Task-setting |
| Le et al., 1998 | 1 | 10.1152/jn.1998.79.3.1535 | 8 | 12 | Attention shifting: Attention shifting > sustained attention | nonverbal | Task-setting |
| Leber et al., 2008 | 1 | 10.1073/pnas.0805423105 | 21 | 20 | cued switching: Switch slope > repeat slope | nonverbal | Task-setting |

| Reference and year | Contrast # | DOI | N | Foci | Task name, brief description and contrast | Input modality | Subcomponent |
|---|---|---|---|---|---|---|---|
| Liu et al., 2003 | 1 | 10.1093/cercor/bhg080 | 13 | 5 | serial visual presentation (with switching component): sCM > hC and sMC > hM (conjunction analysis) | nonverbal | Task-setting |
| Liu et al., 2015 | 1 | 10.1016/j.neuropsychologia.2015.07.019 | 12 | 6 | WCST: Switching > rule application | nonverbal | Task-setting |
| Liu et al., 2015 | 1 | 10.1016/j.neuropsychologia.2015.07.019 | 12 | 23 | WCST: switching > hypothesis testing | nonverbal | Task-setting |
| Loose et al., 2006 | 1 | 10.1016/j.brainres.2006.03.039 | 14 | 4 | figural shifting task: Shift > non-shift (figural) | nonverbal | Task-setting |
| Loose et al., 2006 | 2 | 10.1016/j.brainres.2006.03.039 | 14 | 6 | verbal shifting task: shift > non-shift (verbal) | verbal | Task-setting |
| Luks et al., 2002 | 1 | 10.1006/nimg.2002.1210 | 11 | 2 | cue-target: Neutrally cued switch > repeat targets | verbal | Task-setting |
| Luks et al., 2002 | 2 | 10.1006/nimg.2002.1210 | 11 | 2 | cue-target: switch > repeat targets | verbal | Task-setting |
| Moll et al., 2002 | 1 | 10.1590/S0004-282X2002000600002 | 7 | 7 | TMT: vTMTB (switch) > vTMTA (nonswitch) | verbal | Task-setting |
| Monchi et al., 2001 | 1 | https://doi.org/10.1523/JNEUROSCI.21-19-07733.2001 | 11 | 4 | WCST: Matching after negative feeback > control matching | nonverbal | Task-setting |
| Monchi et al., 2001 | 2 | https://doi.org/10.1523/JNEUROSCI.21-19-07733.2001 | 11 | 5 | WCST: matching after positive feeback > control matching | nonverbal | Task-setting |
| Monchi et al., 2006 | 1 | 10.1523/JNEUROSCI.21-19-07733.2001 | 10 | 8 | montreal card sorting: Retrieval with shift > retrieval without shift | nonverbal | Task-setting |
| Muhle-Karbe et al., 2014 | 1 | 10.1016/j.neuroimage.2014.05.058 | 22 | 12 | Task switching: Full switch > repeat | verbal | Task-setting |
| Muhle-Karbe et al., 2014 | 2 | 10.1016/j.neuroimage.2014.05.058 | 22 | 13 | Task switching: task switch > repeat | verbal | Task-setting |
| Muhle-Karbe et al., 2014 | 3 | 10.1016/j.neuroimage.2014.05.058 | 22 | 10 | Task switching: SR switch > repeat (main effect of transition) | verbal | Task-setting |
| Nagano-Saito et al., 2008 | 1 | 10.1523/JNEUROSCI.3921-07.2008 | 19 | 29 | WCST: set-shifting: negative feedback > control feekback | nonverbal | Task-setting |
| Nagano-Saito et al., 2008 | 2 | 10.1523/JNEUROSCI.3921-07.2008 | 19 | 2 | WCST: set-shifting: positive feedback > control feedback | nonverbal | Task-setting |
| Orr et al., 2014 | 1 | 10.1016/j.neuroimage.2013.08.047 | 28 | 19 | Task switching: Switch > repeat | verbal | Task-setting |
| Orr et al., 2019a | 1 | 10.3758/s13415-019-00689-0 | 19 | 10 | Task switching: Switch > repeat | verbal | Task-setting |
| Orr et al., | 1 | 10.1016/j.neuroimage.2019.116133 | 19 | 15 | Task switching: Proactive switch > | verbal | Task-setting |

| Reference and year | Contrast # | DOI | N | Foci | Task name, brief description and contrast | Input modality | Subcomponent |
|---|---|---|---|---|---|---|---|
| 2019b | | | | | proactive repeat; | | |
| Orr et al., 2019b | 2 | 10.1016/j.neuroimage.2019.116133 | 19 | 8 | Task switching: reactive switch > reactive repeat | verbal | Task-setting |
| Page et al., 2009 | 1 | 10.1016/j.pscychresns.2009.05.002 | 11 | 3 | Task switching: Switch > repeat (switch task) | nonverbal | Task-setting |
| Parris et al., 2007 | 1 | 10.1162/jocn.2007.19.1.13 | 22 | 20 | Rule switching: Flip > hold (Experiment 1) | nonverbal | Task-setting |
| Pessoa et al., 2009 | 1 | 10.1016/j.brainres.2008.10.010 | 20 | 8 | Cue task: Switch > non-switch | nonverbal | Task-setting |
| Peters et al., 2015 | 1 | 10.1523/JNEUROSCI.3795-14.2015 | 20 | 6 | Attention shifting : Shift > hold | nonverbal | Task-setting |
| Philipp et al., 2013 | 1 | 10.1002/hbm.22036 | 23 | 2 | Task switching: switch > repeat (stimulus categorisation) | nonverbal | Task-setting |
| Philipp et al., 2013 | 2 | 10.1002/hbm.22036 | 23 | 1 | Task switching: switch > repeat (response modality) | nonverbal | Task-setting |
| Pollmann et al., 2006 | 1 | 10.1016/j.neuroimage.2005.09.013 | 20 | 5 | Task switching: Change > stay (dimension) | nonverbal | Task-setting |
| Pollmann et al., 2006 | 2 | 10.1016/j.neuroimage.2005.09.013 | 20 | 15 | Task switching: change > stay (response) | nonverbal | Task-setting |
| Provost & Monchi, 2015 | 1 | https://doi.org/10.1111/ejn.12821 | 15 | 25 | Set shifting: Continuous shift > control | nonverbal | Task-setting |
| Rodehacke et al., 2014 | 1 | 10.1371/journal.pone.0088957 | 29 | 7 | Switching task: Switch > repeat | nonverbal | Task-setting |
| Rushworth et al., 2002 | 1 | 10.1152/jn.2002.87.5.2577 | 10 | 4 | Visual switching: Switch > stay (RS task) | nonverbal | Task-setting |
| Rushworth et al., 2002 | 2 | 10.1152/jn.2002.87.5.2577 | 8 | 2 | Response switching: Switch > stay (VS task) | nonverbal | Task-setting |
| Sali et al., 2016 | 1 | 10.1523/JNEUROSCI.2323-15.2016 | 20 | 10 | Attention shifting: Shift > hold | verbal | Task-setting |
| Sali et al., 2020 | 1 | 10.1162/jocn_a_01541 | 23 | 3 | Attention shifting: Shift > hold | verbal | Task-setting |
| Schmitz et al., 2006 | 1 | 10.1016/j.biopsych.2005.06.007 | 12 | 9 | Task switching: Switch > repeat (switch task) | nonverbal | Task-setting |
| Sekutowicz et al., 2016 | 1 | 10.1016/j.neuroimage.2016.07.046 | 108 | 22 | Rule switching: Switch > repetition | verbal | Task-setting |
| Shi et al., 2018 | 1 | 10.1002/hbm.23878 | 32 | 21 | Cued task switching: Task switching > task repetition | verbal | Task-setting |
| Shomstein & | 1 | 10.1523/JNEUROSCI.4408-05.2006 | 12 | 4 | Attention shifting: Shift > hold (spatial | verbal | Task-setting |

| Reference and year | Contrast # | DOI | N | Foci | Task name, brief description and contrast | Input modality | Subcomponent |
|---|---|---|---|---|---|---|---|
| Yantis, 2006 spatial | | | | | experiment) | | |
| Shomstein & Yantis, 2006 nonspatial | 2 | 10.1523/JNEUROSCI.4408-05.2006 | 13 | 8 | Attention shifting: Shift > hold (non-spatial experiment) | verbal | Task-setting |
| Smith et al., 2004 | 1 | 10.1002/hbm.20007 | 20 | 10 | WCST: Switch > repeat | nonverbal | Task-setting |
| Stoppel et al., 2013 | 1 | 10.1093/cercor/bhs116 | 16 | 7 | Task switching: Switch locations > hold and switch objects > hold (conjunction analysis) | nonverbal | Task-setting |
| Swainson et al., 2003 | 1 | 10.1162/089892903322370717 | 12 | 3 | Rule switching: Switch > non-switch (WAIT task) | nonverbal | Task-setting |
| Swainson et al., 2003 | 2 | 10.1162/089892903322370717 | 12 | 2 | Rule switching: switch > non-switch (GO task) | nonverbal | Task-setting |
| Townsend et al., 2006 | 1 | 10.1016/j.neuroimage.2006.01.045 | 10 | 14 | Attention shifting: Younger- Shift > focus | nonverbal | Task-setting |
| Trempler et al., 2016 | 1 | 10.1162/jocn_a_01040 | 20 | 5 | Digit sequence: Switch > drift | nonverbal | Task-setting |
| Uehara et al., 2019 | 1 | 10.1016/j.brainres.2019.146365 | 32 | 10 | Sequential finger tapping: Switch > repeat | nonverbal | Task-setting |
| Vallesi et al., 2012 | 1 | 10.1002/hbm.21312 | 12 | 1 | Rule switching: switch > no-switch (cue-related) | nonverbal | Task-setting |
| Vallesi et al., 2012 | 1 | 10.1002/hbm.21312 | 12 | 1 | Rule switching: switch > no-switch (target-related) | nonverbal | Task-setting |
| Vallesi et al., 2015 | 1 | 10.1016/j.cortex.2015.01.016 | 31 | 10 | verbal and spatial switching: Switch > single (spatial task) | verbal, nonverbal | Task-setting |
| Vandenberghe et al., 2001 | 1 | 10.1006/nimg.2001.0860 | 11 | 4 | Attention shifting: Shift > maintain (Experiment 1) | nonverbal | Task-setting |
| Wang et al., 2015 | 1 | 10.1371/journal.pone.0140731 | 16 | 8 | WCST: WCST: 2-NF > 0-NF and 1-NF > 0-NF (conjunction analysis) | nonverbal | Task-setting |
| Witt & Stevens, 2012 | 1 | 10.1016/j.neuroimage.2012.05.007 | 134 | 23 | set shifting: Switch > non-switch | verbal | Task-setting |
| Witt & Stevens, 2013 | 1 | 10.1016/j.neuroimage.2012.09.072 | 83 | 23 | set shifting: Switch > non-switch | nonverbal | Task-setting |
| Woodcock et al., 2010 | 1 | 10.1016/j.brainres.2010.09.093 | 8 | 10 | task-switching: Switch > non-switch | nonverbal | Task-setting |
| Wylie et al., 2004 | 1 | 10.1002/hbm.20003 | 12 | 13 | task switching: Switch 3 > switch 1 (Group 1) | nonverbal | Task-setting |

| Reference and year | Contrast # | DOI | N | Foci | Task name, brief description and contrast | Input modality | Subcomponent |
|---|---|---|---|---|---|---|---|
| Wylie et al., 2004 | 2 | 10.1002/hbm.20003 | 12 | 17 | task switching: Switch 3 > switch 1 (Group 2) | nonverbal | Task-setting |
| Wylie et al., 2005 | 1 | 10.1093/cercor/bhi118 | 13 | 19 | Rule switching: Switch > repeat targets (colour) | nonverbal | Task-setting |
| Wylie et al., 2005 | 2 | 10.1093/cercor/bhi118 | 13 | 4 | task switching: switch > repeat targets (speed) | nonverbal | Task-setting |
| Xu et al., 2015 | 1 | 10.3389/fnhum.2015.00034 | 18 | 10 | task switching: Switch > SwitchGo | nonverbal | Task-setting |
| Zanolie et al., 2008 | 1 | 10.1016/j.cortex.2007.11.005 | 18 | 17 | Set shifting: intradimensional negative feedback > negative feedback | nonverbal | Task-setting |
| Zanolie et al., 2008 | 2 | 10.1016/j.cortex.2007.11.005 | 18 | 4 | Set shifting: intradimensional negative feedback > positive feedback | nonverbal | Task-setting |
| Yoon et al., 2021 | 1 | 10.1016/j.nicl.2021.102590 | 26 | 14 | WCST: WCST: RNF > RPF (healthy) | nonverbal | Task-setting |
| Zuhlsdorff et al., 2023 | 1 | 10.1093/cercor/bhac431 | 40 | 26 | "Change your mind task" - switching under uncertainty, without rule-based learning: changing > repeating | nonverbal | Task-setting |
| Hummos et al., 2022 | 1 | 10.1371/journal.pcbi.1010500 | 28 | 16 | probabilistic inference task: switching > staying | nonverbal | Task-setting |
| Nee & Jonides, 2014 | 1 | 10.1523/JNEUROSCI.0130-14.2014 | 26 | 14 | Attention switching: Focus switch > focus repeat | verbal | Task-setting |
| Nee & Jonides, 2014 | 2 | 10.1523/JNEUROSCI.0130-14.2014 | 26 | 6 | Behavioural switching: active switch > active repeat | verbal | Task-setting |
| Goel et al., 2000 | 1 | https://doi.org/10.1006/nimg.2000.0636 | 11 | 13 | verbal syllogistic reasoning: main effect of verbal syllogistic reasoning | verbal | Task-setting |
| Christoff et al., 2001 | 1 | https://doi.org/10.1006/nimg.2001.0922 | 10 | 2 | RPM: RPM 2-relational > 1-relational | nonverbal | Task-setting |
| Parsons and Osherson, 2001 | 1 | https://doi.org/10.1093/cercor/11.10.954 | 10 | 19 | verbal reasoning: probabilistic reasoning (whether the conclusion of an argument was more likely to be true than false) > deduction (distinguish valid from invalid arguments) | nonverbal | Task-setting |
| Kroger et al., 2002 | 1 | https://doi.org/10.1093/cercor/12.5.477 | 8 | 18 | RPM: RPM (relational complexity) | verbal | Task-setting |
| Knauff et al., 2002 | 1 | https://doi.org/10.1016/S0028-3932(03)00016-2 | 12 | 9 | verbal syllogistic reasoning: syllogistic reasoning > baseline | verbal | Task-setting |
| Fangmeier et al., 2006 | 1 | https://doi.org/10.1162/jocn.2006.18.3.320 | 12 | 9 | verbal syllogistic reasoning: syllogistic reasoning problem (reasoning validation phase) > rest | verbal | Task-setting |

| Reference and year | Contrast # | DOI | N | Foci | Task name, brief description and contrast | Input modality | Subcomponent |
|---|---|---|---|---|---|---|---|
| Monti et al., 2007 | 1 | https://doi.org/10.1016/j.neuroimage.2007.04.069 | 10 | 32 | verbal syllogistic reasoning: complex deductions > simple deductions (syllogistic reasoning words and pseudowords; experiment 1) | verbal | Task-setting |
| Monti et al., 2007 | 1 | https://doi.org/10.1016/j.neuroimage.2007.04.069 | 12 | 26 | verbal syllogistic reasoning: complex deductions > simple deductions (syllogistic reasoning with face and house; experiment 2) | verbal | Task-setting |
| Reverberi et al., 2007 | 1 | https://doi.org/10.1016/j.neuroimage.2007.07.060 | 14 | 8 | verbal syllogistic reasoning: deductive inference (syllogistic) > baseline | verbal | Task-setting |
| Masunaga et al., 2008 | 1 | https://doi.org/10.1016/j.intell.2008.01.006 | 18 | 16 | Nonverbal intelligence test: nonverbal intelligence test (Cattell's Culture Fair Intelligence Test - Topology) > scrambled stimuli | nonverbal | Task-setting |
| Wendelken et al., 2008 | 1 | https://doi.org/10.1016/j.neuroimage.2009.06.025 | 20 | 8 | analogical reasoning verbal: compare two related words > match a related word | verbal | Task-setting |
| Wartenburger et al., 2009 | 1 | https://doi.org/10.1016/j.neuroimage.2009.06.025 | 15 | 5 | analogical reasoning nonverbal: difficult geometric analogical reasoning (whether B:B" matches A:A") > less difficult problems | nonverbal | Task-setting |
| Geake and Hansen, 2010 | 1 | https://doi.org/10.1016/j.neuroimage.2009.09.008 | 16 | 34 | analogical reasoning nonverbal: nonsemantic analogical reasoning > baseline | nonverbal | Task-setting |
| Golde et al., 2010 | 1 | https://doi.org/10.1016/j.neuroimage.2009.09.009 | 16 | 12 | RPM: RPM abstract > RPM action | nonverbal | Task-setting |
| Krawczyk et al., 2011 | 1 | https://doi.org/10.1016/j.cortex.2010.04.008 | 20 | 11 | RPM: RPM > baseline | nonverbal | Task-setting |
| Volle et al., 2010 | 1 | https://doi.org/10.1093/cercor/bhq012 | 16 | 25 | analogical reasoning (verbal, less semantic): analogy > matching (alphabet) | verbal | Task-setting |
| Cho et al., 2010 | 1 | https://doi.org/10.1093/cercor/bhp121 | 17 | 9 | analogical reasoning (semantic, nonverbal): people pieces analogy task with cartoon characters (task > baseline) | nonverbal | Task-setting |
| Prado et al., 2010 | 1 | https://doi.org/10.1016/j.neuroimage.2010.03.026 | 13 | 12 | verbal reasoning: integrable arguments > nonintegrable arguments | verbal | Task-setting |
| Crescentini et al., 2011 | 1 | https://doi.org/10.1523/JNEUROSCI.4579-10.2011 | 20 | 12 | rule detection task: nonverbal nonsemantic rule acquisition > rule following | nonverbal | Task-setting |

| Reference and year | Contrast # | DOI | N | Foci | Task name, brief description and contrast | Input modality | Subcomponent |
|---|---|---|---|---|---|---|---|
| Jia et al., 2011 | 1 | https://doi.org/10.1016/j.neuroimage.2011.03.020 | 20 | 16 | rule detection task: arithmetic inductive reasoning (Rule induction > rule judgement; Table 2) | nonverbal | Task-setting |
| Shokri-Kojori et al., 2012 | 1 | https://doi.org/10.1038/srep00411 | 20 | 20 | visuospatial reasoning: visuospatial reasoning task that assembles RPM (3 relational > 1 relational; Table S2) | nonverbal | Task-setting |
| Watson and Chatterjee, 2012 | 1 | https://doi.org/10.1016/j.neuroimage.2011.09.030 | 23 | 3 | nonverbal analogical reasoning: analogical reasoning of shapes (analogy > matching) | nonverbal | Task-setting |
| Reverberi et al., 2012 | 1 | https://doi.org/10.1016/j.neuroimage.2011.08.027 | 21 | 15 | syllogistic reasoning: syllogistic reasoning (integratable > nonintegratable sentences) | verbal | Task-setting |
| Cocchi et al., 2014 | 1 | https://doi.org/10.1093/cercor/bht075 | 21 | 18 | deductive reasoning: rule > null | verbal and nonverbal | Task-setting |
